# A deep learning model captures position-specific effects of plant regulatory sequences and suggests genes under complex regulation

**DOI:** 10.1101/2025.08.30.673246

**Authors:** Kevin C. Rockenbach, Silvia F. Zanini, Alison C. Tidy, Richard J. Morris, Rachel Wells, Agnieszka A. Golicz

## Abstract

Deep neural networks can be trained to predict gene expression directly from genomic sequence, thereby implicitly learning regulatory sequence patterns from scratch, minimizing the bias imposed by prior assumptions. A challenging, yet promising prospect is the extraction of novel insights into gene-regulatory mechanisms, by probing and interpreting such gene expression models. Using a branched convolutional neural network architecture trained on promoter and terminator sequences we predict gene expression for allopolyploid *Brassica napus* and the closely related model organism *Arabidopsis thaliana*. We validate the model by comparing predicted and measured expression across ecotypes. We also show that deep learning models can successfully capture the positional binding preferences of some transcription factor families, without having been trained on transcription factor binding data. Furthermore, we show that our model did not only detect local sequence patterns, but was also able to determine their function based on their positional context. We also found that increased prediction error correlated with additional more distal or epigenetic regulatory input. Our results demonstrate that deep learning can be used to understand the regulatory architecture of gene expression in plants. A better understanding of gene regulation in the context of polyploid genomes is of particular economic importance, due to their prevalence among major crops. In the future, we hope that such models may facilitate the targeted engineering of gene regulation in crops.

## Background

Although complete genome assemblies of eukaryotic organisms have been available since the turn of the millennium [1–3], deciphering the grammar by which regulatory information is encoded in the genome remains challenging [4]. At the level of transcriptional regulation, transcription factors (TFs) are known to play an important role in facilitating the assembly of the transcriptional machinery [5]. TFs are DNA-binding proteins that can selectively bind to cis-regulatory elements (CREs) in the genome based on sequence- and shape-dependent physical interactions with DNA, as well as interactions with other transcription factors and epigenetic factors [6–9]. Genetic variations in CREs that lead to differences in the ability of TFs to bind, have been shown to explain most of the heritable phenotypic variation in maize [10]. CREs tend to be clustered into cis-regulatory modules (CRMs), such as promoters, enhancers, silencers and insulators, but the underlying grammar of the constituent CREs, meaning the set of rules governing their relative positioning and orientation, is not well understood [11]. Active regulatory sequences are typically associated with accessible chromatin, which enables binding of TFs to take place. About 75% of accessible chromatin regions in plants are located outside of transcribed regions but within 2 kb upstream of the transcription start site (TSS) or 1 kb downstream of the transcription termination site (TTS), indicating a much more localized regulation of gene expression compared to animals [12, 13]. This aligns with in vivo profiling of sequences bound by TFs in maize [14], which showed that 69% of TFs were bound to sequences in and around genes and 31% were bound to intergenic regions. Likewise, an in vitro profiling method for TF occupancy performed in *Arabidopsis thaliana* [8] found that CREs were enriched in promoters and 5’ UTRs and moderately depleted in coding sequences. Voichek et al. [15] experimentally confirmed that 5’ UTRs and introns downstream of the TSS play a profound role in TF-mediated transcriptional regulation. In particular, they showed that TFs of the C2C2-GATA family were able exert a strong positive influence on gene expression, independent of the genomic context, across plant species and most tissues, as long as the respective binding motif was located within 500 bp downstream of the TSS [15]. They suggested that the position-dependent activity of plant enhancers is a defining feature that distinguishes them from enhancers in other kingdoms of life. However, there is also increasing evidence for position-dependent regulatory activity of TFs in humans [4, 16] and yeast [17]. Many TF families in plants show characteristic positional binding preferences relative to the TSS [15, 18, 19] and in some cases, such as C2C2-GATA factors, there appears to be a clear functional dependence between their regulatory activity and their binding position relative to the TSS. Nevertheless, it is largely unknown to what extent the function of other TF families depends on their binding position relative to the TSS, since positional preferences can also arise as a byproduct of other dependencies such as the co-occurrence of CREs within CRMs [20].

Convolutional neural networks (CNNs), originally developed for image recognition tasks [21, 22], have since become well-established as a tool for automatically identifying relevant DNA motifs and learning to interpret their local grammar [23–25]. CNNs process input sequences in a layer-wise manner, employing convolutional filters that scan across the input and aggregate local input features into higher-order output features. Convolutional layers are typically interspersed with pooling layers, which successively reduce the spatial dimensionality of the input and, thereby, increase the receptive field of subsequent convolutional layers. Using these operations, standard CNNs can integrate information over a distance of up to 20 kb [26]. Various flavors of CNNs have been successfully used to predict gene expression and other molecular phenotypes across a wide range of organisms and were used to gain insight into the regulatory architecture of gene expression at a genome-wide scale [17, 26–37]. Among them, Wrightsman et al. [32] trained models to predict chromatin accessibility and DNA methylation using data from a wide range of plant species and observed that the models generalized well across species, with performances comparable to models trained within species. Washburn et al. [28] built a model which was able to categorize genes based on their expression level (high or low) in maize based on 1.5 kb of genomic sequence surrounding transcription start and termination sites, respectively. Peleke et al. [34], extended the model and identified distinct expression-predictive motifs (EPMs) associated with gene activation or repression. Importantly, subsequent analysis of the positional distribution of EPMs suggested that their function may be position-dependent. Interestingly, both the Washburn and Peleke models [28, 34] highlighted the importance of 5’ and 3’ untranslated regions (UTRs) for prediction. This was achieved by computing importance (saliency) scores across the sequence regions used for prediction. Washburn et al. and Peleke et al. built classification models assigning low and high expression categories. Opdebeeck et al. [37] published a model for classifying the responsiveness of genes to abscisic acid (ABA) in *A. thaliana* and were able to identify TFs that are likely to be involved in ABA-mediated gene regulation, for which this role was previously unknown. To our knowledge, only four models predicting gene expression values (regression models) in plants have been published thus far [24, 35, 38, 39], but only Li et al. [38] have performed extensive hyperparameter optimization. They built an ensemble model consisting of multiple CNNs. Most recently, Qui et al. [39] adapted an existing architecture to identify putative CREs in four plant species. They were able to explain 41% of the variation in maximum gene expression in *A. thaliana*.

Here, we achieved state-of-the-art performance in predicting gene expression from proximal regulatory sequences using a branched CNN architecture, without the need for ensemble modeling. We applied a multi-step approach to hyperparameter optimization, where we not only optimized the architecture, but also the learning algorithm. With this approach, we were able to explain 53% of the variation in median gene expression, on a test set of unseen gene families in allotetraploid *Brassica napus*, using only the non-coding portions of proximal genomic sequences. We argue that highly effective species-specific models can be created for any species of interest, even complex polyploid species, provided that a genome assembly and appropriate gene expression data are available. We found that the model was able to recapitulate the position-specific upregulation effect of C2C2-GATA TFs recently described by Voichek et al. [15]. The presence of the corresponding motif resulted in higher gene expression predictions only when found downstream of the TSS. Finally, inspection of genes whose expression was poorly predicted by the model revealed an over-representation of genes associated with additional regulatory enhancer elements and polycomb repressive complexes. In essence, the model can help distinguish genes that are under control by proximal DNA sequence from genes that have additional regulatory control. Taken together, our results show that substantial novel biological insight can be derived from a highly optimized CNN model.

## Results

### Deep learning models can be effectively trained on polyploid data

#### *n*emo predicts gene expression from proximal regulatory sequences

We implemented a branched convolutional neural network architecture called *n*emo (***n****apus* **e**xpression **mo**del), for gene expression prediction from promoter and terminator sequences in allotetraploid *B. napus* (Fig. 1). We focused on prediction of median gene expression across tissues, which was supported by overall high correlation across tissues (Fig. S1, S2) and has previously been successfully performed on human and mouse data [30]. The architecture contains separate convolutional branches to independently extract features from the two sequence inputs. The model inputs span 5 kb upstream of the TSS to 1.2 kb downstream of the TSS and 1.2 kb upstream of the TTS to 5 kb downstream of the TTS, respectively. Ther performance of the *n*emo architecture, trained on a fixed 80% split of the *B. napus* data (*n*emo_80_) and tested on a held-out test set (10% of the data) containing unseen gene families, performed better (*r*^2^ = 0.53) than the Xpresso architecture (*r*^2^ = 0.49), when trained and tested on the same data sets (Fig. 2A). When incorporating the validation set into the training set and therefore training the *n*emo architecture on 90% of the data (*n*emo_90_), the performance on the held-out test set slightly increased (*r*^2^ = 0.54). Unlike the Xpresso architecture, the *n*emo architecture does not rely on additional handcrafted sequence features. However, the *n*emo architecture is also much larger, with a little less than 7 · 10^6^ trainable parameters compared to approximately 10^5^ trainable parameters for the Xpresso architecture. The performance of the *n*emo architecture on *B. napus* data was similar to the reported performance of the Xpresso architecture on human data (*r*^2^ = 0.59) [30]. Although these performance estimates cannot be directly compared between studies, due to differences in data preprocessing, this result gives a rough indication that proximal genomic sequences may have a similar potential for explaining gene expression in plants and animals, respectively. This is consistent with Li et al. [38], who reported similar performance estimates in plants, and Zrimec et al. [31], who found that a gene expression model trained on data from *A. thaliana* only performed marginally better than the same model trained on human data, despite large differences in gene density between the two species. Taken together, based on the performance of various proximal sequence-to-expression models including our own, the proportion of proximal gene regulation does not appear to be higher in plants compared to mammals, despite large differences in chromatin organization that would suggest otherwise [12, 40].

**Fig. 1.**
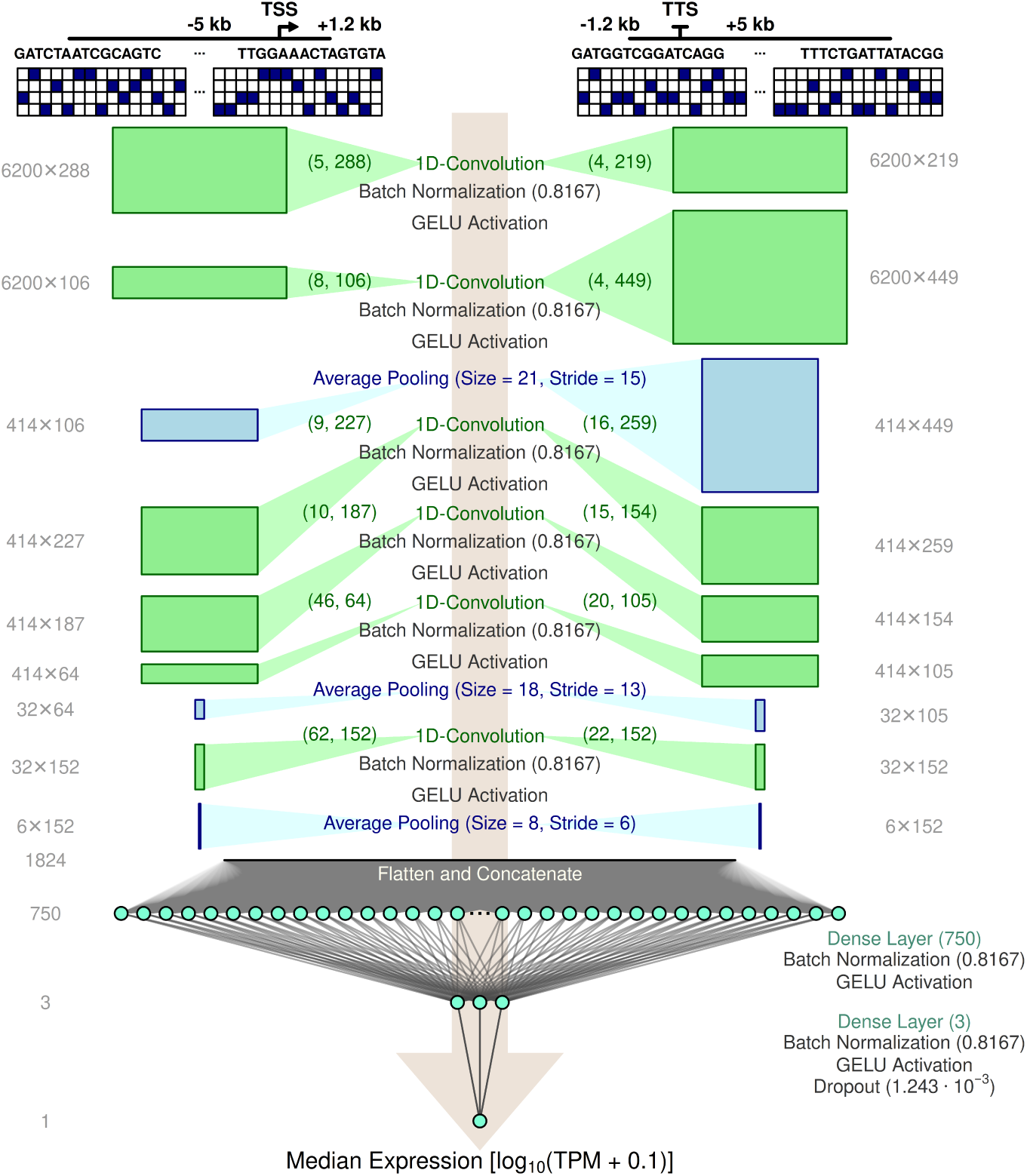
The *n*emo architecture. Optimized hyperparameters of the branched convolutional neural network architecture. Convolutional kernel sizes and the number of filters for each convolutional layer are given in green braces. The dimensions of boxes are scaled to the output dimensions of the respective layer in the respective branch. Numerical values given for batch normalization layers, dense layers and the dropout layer represent values for the momentum of the moving average, the number of neurons and the dropout rate, respectively.

**Fig. 2.**
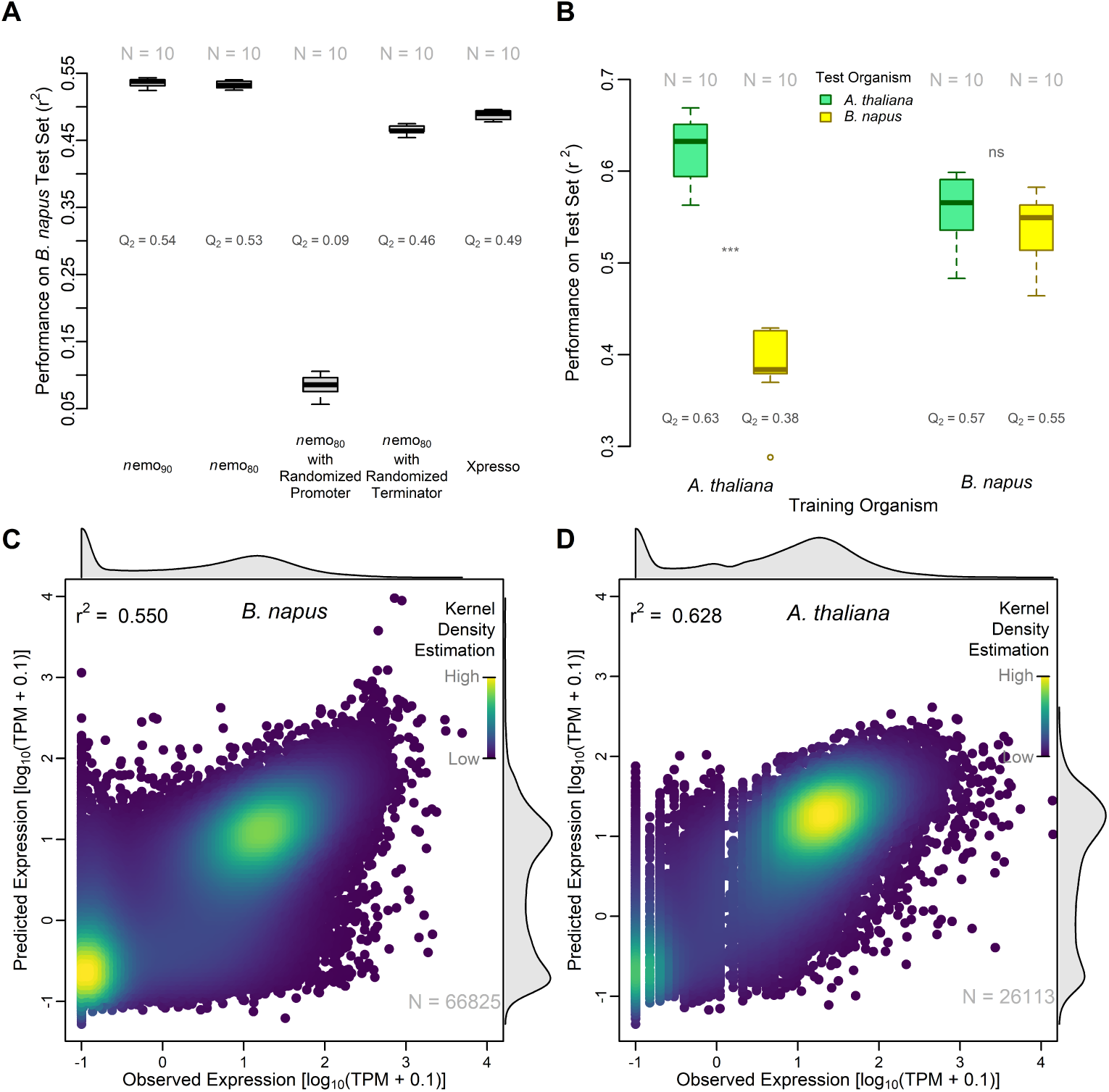
*n*emo robustly predicts median gene expression. (A) Performance comparisons on unseen *B. napus* test data. From left to right: Ten *n*emo_90_ instances, trained on the same training set, consisting of all the data used during hyperparameter optimization, including the validation set; ten *n*emo_80_ instances, trained on the 80% of data used as a training set during hyperparameter optimization; the same ten *n*emo_80_ instances tested against mutated versions of the test set where promoter and terminator sequences were replaced with random sequences, respectively; ten Xpresso instances trained on the same training set and tested on the (non-randomized) test set. (B) Cross-species comparison between *n*emo_90_ trained on *B. napus* data and *n*emo_90_ trained on *A. thaliana* data. The box plots represent test performance across all ten fold permutations. *B. napus*test sets were randomly down-sampled to match the smaller *A. thaliana* test set size. (C, D) Minimally biased predictions on jointly partitioned data. Ten instances of *n*emo_90_ were trained on different permutations of data folds from *B. napus* (C) and *A. thaliana* (D) respectively. Each instance was tested on the respective held-out test set. All ten sets of test predictions for each species were then concatenated and plotted against the observed median expression, to arrive at a minimally biased set of predictions across the entire data set. Colors represent gaussian kernel density estimations (KDE).

To test the contributions of the two input sequences to the predictive performance of the *n*emo architecture, we derived variants of the test set with randomized promoter or terminator sequences, while maintaining the pattern of masked segments, and performed test predictions on these modified inputs. When randomizing the terminator sequence while maintaining the promoter sequence (*r*^2^ = 0.46), *n*emo was able to explain 7% less of the variation in gene expression compared to the unmodified inputs, indicating that the contribution of the terminator sequence to the model’s predictions is non-zero, but fairly marginal. However, when randomizing the promoter sequence, the correlation between predicted and observed gene expression disappeared almost entirely (*r*^2^ = 0.09), suggesting a crucial role of the promoter region, which includes the 5’ UTR, in setting a coarse baseline of expression, which then may be up-or down-regulated by cis-regulatory elements in the terminator and distal enhancers.

#### Polyploid training data increases model robustness

For cross-species performance comparisons, we jointly partitioned data from *B. napus* and *A. thaliana* so that evolutionary dependencies would only occur between the corresponding data folds of the two species. We then performed 10-fold cross-testing where, for each species, we trained ten *n*emo_90_ model instances on 90% of the data. Each model instance was trained on a different permutation of data folds and tested on the remaining data fold of the same species and the corresponding data fold from the other species (Fig. 2B). The predictions made by the ten model instances of each species were aggregated to obtain minimally biased predictions across the complete data sets of both species (Fig. 2C,D). Since predictions made on the jointly partitioned data were also made on gene families that were used during hyperparameter optimization, the cross-species performances should only be evaluated in relation to each other and not in absolute terms. The *n*emo_90_ instances trained on *A. thaliana* data were able to explain about 8% more of the variation in observed median expression in *A. thaliana* (*r*^2^ = 0.63) than the model instances trained on *B. napus* data were able to explain in *B. napus* (*r*^2^ = 0.55). However, while the model instances trained on *B. napus* data showed no significant difference in performance when tested on data from *A. thaliana* (*r*^2^ = 0.57; Wilcoxon rank sum test: *N* = 10, *p* = 0.2475), the model instances trained on data from *A. thaliana* performed significantly worse when tested on data from *B. napus* (*r*^2^ = 0.38; Wilcoxon rank sum test: *N* = 10, *p* = 1.083·10^−5^) (Fig. 2B). This indicates that models trained on data that is based on a manually curated high quality genome annotation may be able to achieve a higher peak performance when tested on data from the same source, since TSSs, TTSs and intron exon boundaries are well defined, but do not perform robustly on data that is based on minimally or non-curated annotations, which may contain less well-defined structural features and a higher rate of pseudogenes. On the other hand, previous studies have also reported immense difficulty in transferring models from simple diploid genomes to complex polyploid genomes [39]. Models trained on polyploid genomes are exposed to a larger quantity of data and in particular, a larger quantity of redundant sequences and, therefore, may be able to implicitly learn how variations between similar sequences affect gene expression. In situations where the number of training examples is limited, data augmentation can help to improve models [37], however, we argue that training models on polyploid organisms provides and inherent form of data augmentation. Gene family-guided partitioning of data not only prevents data leakage between sets, but also maximizes the sequence redundancy that the model is exposed to during training, since each gene family is concentrated in one of the sets, instead of being scattered between sets. The fact that model instances trained and tested on data from *A. thaliana* achieved a higher peak performance than model instances trained and tested on data from *B. napus*, served as an encouraging sign that the high degree of sequence redundancy found in the *B. napus* genome had been accounted for and therefore did not lead to overinflated performance estimates.

#### *n* emo predictions are consistent with real-world gene expression variation

To further validate nemo predictions, we tested whether the predictions agree with differences in genes expression observed across two oilseed rape genotypes: Express617 (winter type) and ZS11 (semi-winter type). Structural gene annotation was transferred from Express617 to ZS11 and gene expression prediction was made for all genes across the two genotypes. We then selected the top 100 and 1000 genes with the most extreme differences in expression levels between the two genotypes, meaning where the predicted Expressed 617 expression was either higher (Express617 *>* ZS11) or lower (Express617 *<* ZS11) than ZS11. Actual (observed) gene expression levels were then compared between genotypes. On average, we confirmed that the model was able to correctly identify the allele with higher expression (Fig. 3A). Recently, it was postulated that structural variants (SVs) play an important role in rewiring gene expression in Brassica napus [41, 42]. We therefore tested if the genes predicted to have different expression across the two genotypes are over-represented in SVs, and indeed observed significant over-representation of this subset (Fig. 3B). Selected examples point to these SVs affecting open chromatin and therefore putative regulatory regions (Fig. 3C). Taken together, these results demonstrate the robust performance of nemo and its ability to predict differences in expression by accounting for real-world variation between genotypes.

**Fig. 3.**
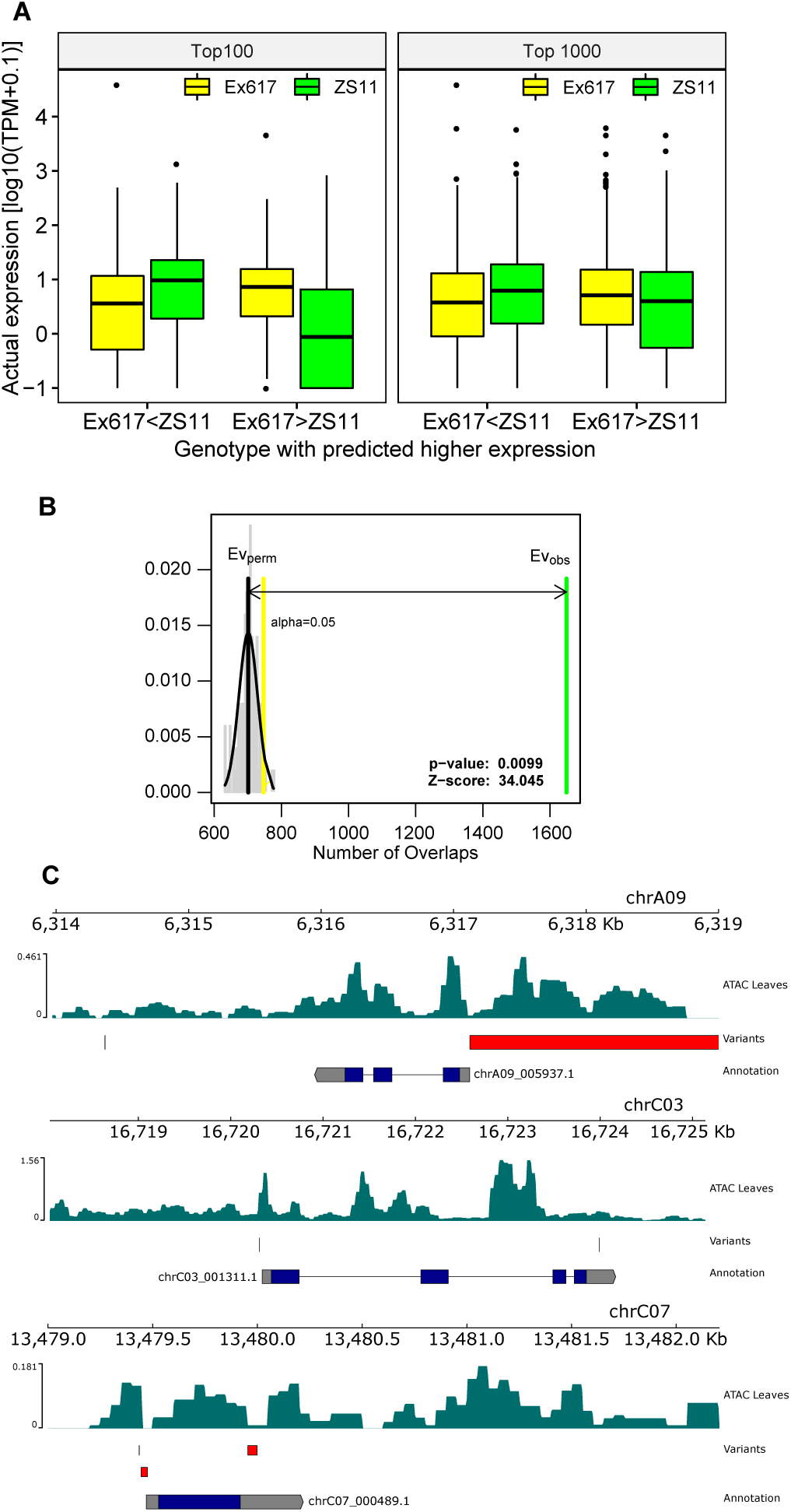
Validation of *n*emo gene expression prediction across Brassica napus ecotypes. (A) *n*emo correctly predicts alleles with higher expression across the two ecotypes. (B) Genes predicted to have difference in expression are over-represented in structural variants. (C) Example of genes which are predicted to have higher expression in Express 617 (reference) compared to ZS11. ZS11 harbors variants (insertions (thin lines) and deletion (red boxes)) in regions adjacent to TSS.

### Convolutional neural networks can learn positional binding preferences of plant TFs

#### Importance scores reflect experimentally derived binding preferences of TFs

In order to assess which sequence features the model predictions were based on, we attributed prediction outcomes to features in the input sequences of specific genes using Shapley valuesand then used the Shapley values to calculate importance scores across genes. To gain an overall picture, we first classified well-predicted genes into three expression bins using the upper and lower quartiles of observed expression values as thresholds (Fig. 4A,C), and calculated importance scores across all genes in the three respective groups (Fig. 4B,D). For all three expression classes, the 5’ and 3’ UTRs were, on average, more important for making predictions than the regions upstream of the TSS and downstream of the TTS. This is in agreement with similar findings from Washburn and colleagues [28], as well as Peleke and colleagues [34]. Due to the additive property of Shapley values [43], highly expressed genes tend to be associated with high importance scores, with the caveat that importance scores are based on absolute Shapley values, making no distinction between positive and negative values, which may cancel each other out. The differences between the three expression classes also tended to be most pronounced in the UTR regions. In addition, in the promoter regions of *B. napus* (Fig. 4B), lowly and moderately expressed genes tended to have slightly higher importance scores than highly expressed genes upstream of the TSS. We hypothesized that some of the observed patterns could be due to differential presence and effect of TF binding sites.

**Fig. 4.**
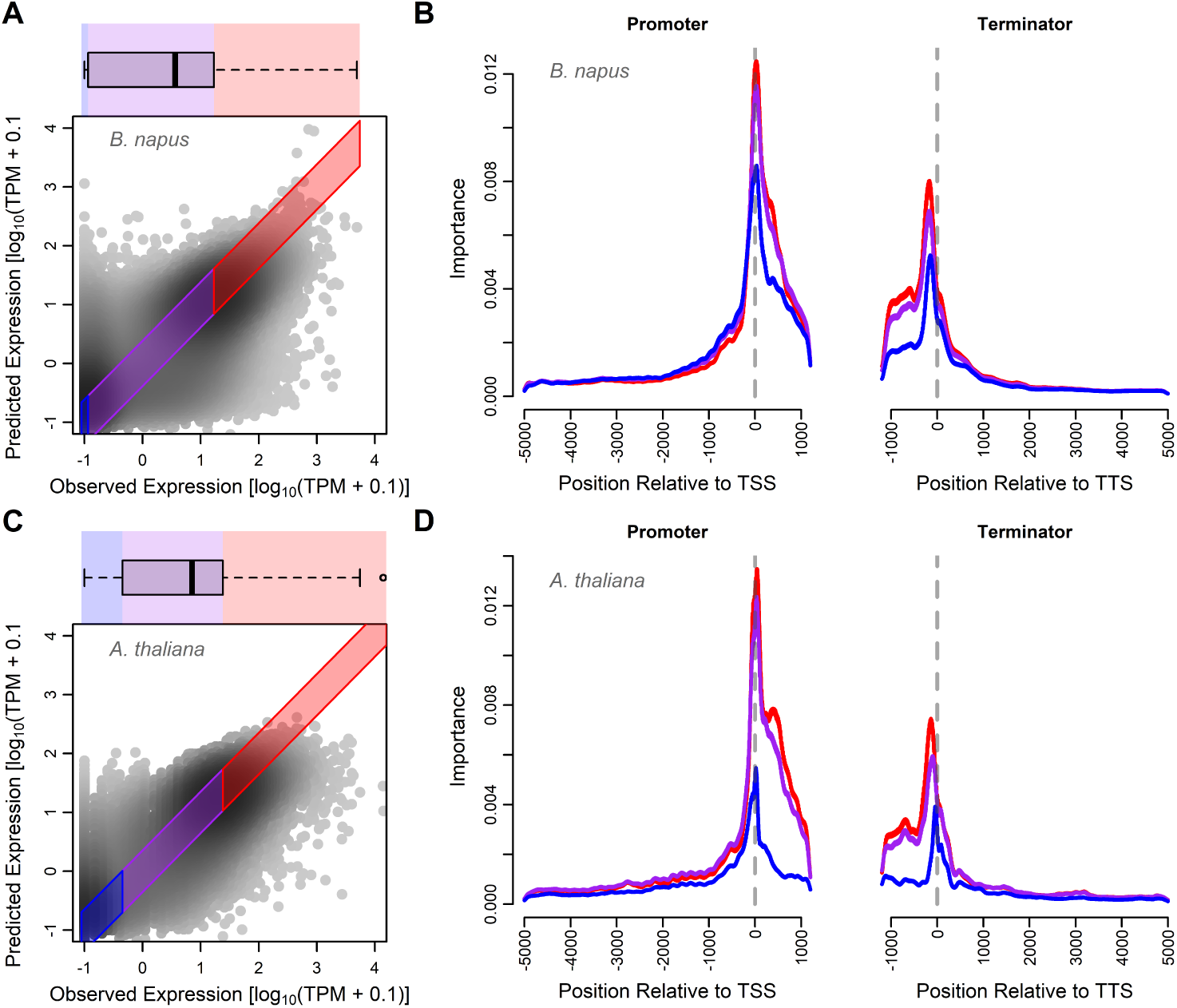
Expression-specific importance. (A, C) Segmentation of well-predicted genes (colored areas) into highly (red), moderately (purple) and lowly (blue) expressed groups, based on the upper and lower quartiles of observed expression values for *B. napus* and *A. thaliana* respectively. (B, D) Meta-gene plots of group specific importances along the two input sequences for *B. napus* and *A. thaliana* respectively.

To investigate the possibility of importance scores reflecting distinct effects of different TF families, *A. thaliana* genes were classified based on experimentally derived information (DAP-Seq [8]) on TF binding to their promoters. Genes of *B. napus* were then classified based on homology to the respective *A. thaliana* genes. Importance scores were calculated for each group and normalized(Fig. S7, S8) to highlight position-specific differences between the groups (Fig. 5B,C). For promoters of *A. thaliana*, the normalized importance scores (Fig. 5C) of some of the groups showed a clear correspondence with the respective DAP-Seq profile (Fig. 5A). For TFs of the MYB-related, Homeobox, C2C2-Dof and MADS families which have a preference for binding up to several hundred base pairs upstream of the TSS, the normalized importance scores to a varying degree, but distinguishably highlighted this region in genes belonging to the respective groups. For TFs of the TCP family, which preferentially bind immediately upstream of the TSS, the corresponding normalized importance scores distinctly highlighted this area. The same was also true for TF families that preferentially bind downstream of the TSS, such as HSF, AP2/EREBP and C2C2-GATA. For *B. napus*, similar regional patterns were visible, especially for Homeobox, MADS, HSF, AP2/EREBP and C2C2-GATA factors. When taking into account the entire interval ± 500 bp around the TSS, we observed particularly strong positive Spearman correlations between the DAP-Seq signal of the three TF families that have the strongest tendency to bind downstream of the TSS (HSF, AP2/EREBP and C2C2-GATA) and the respective normalized importance scores in both species (Fig. S9A,B). Some of the discrepancies between DAP-Seq signals and normalized importance scores may be explained by the presence of other important features, such as co-occurring CREs that bind TFs of other families (Fig. S9C). A sharper boundary between importance scores of upstream and downstream positions was observed in *A. thaliana* (Fig. S8) compared to *B. napus* (Fig. S7), indicating a lower precision of annotated TSSs in *B. napus*.

**Fig. 5.**
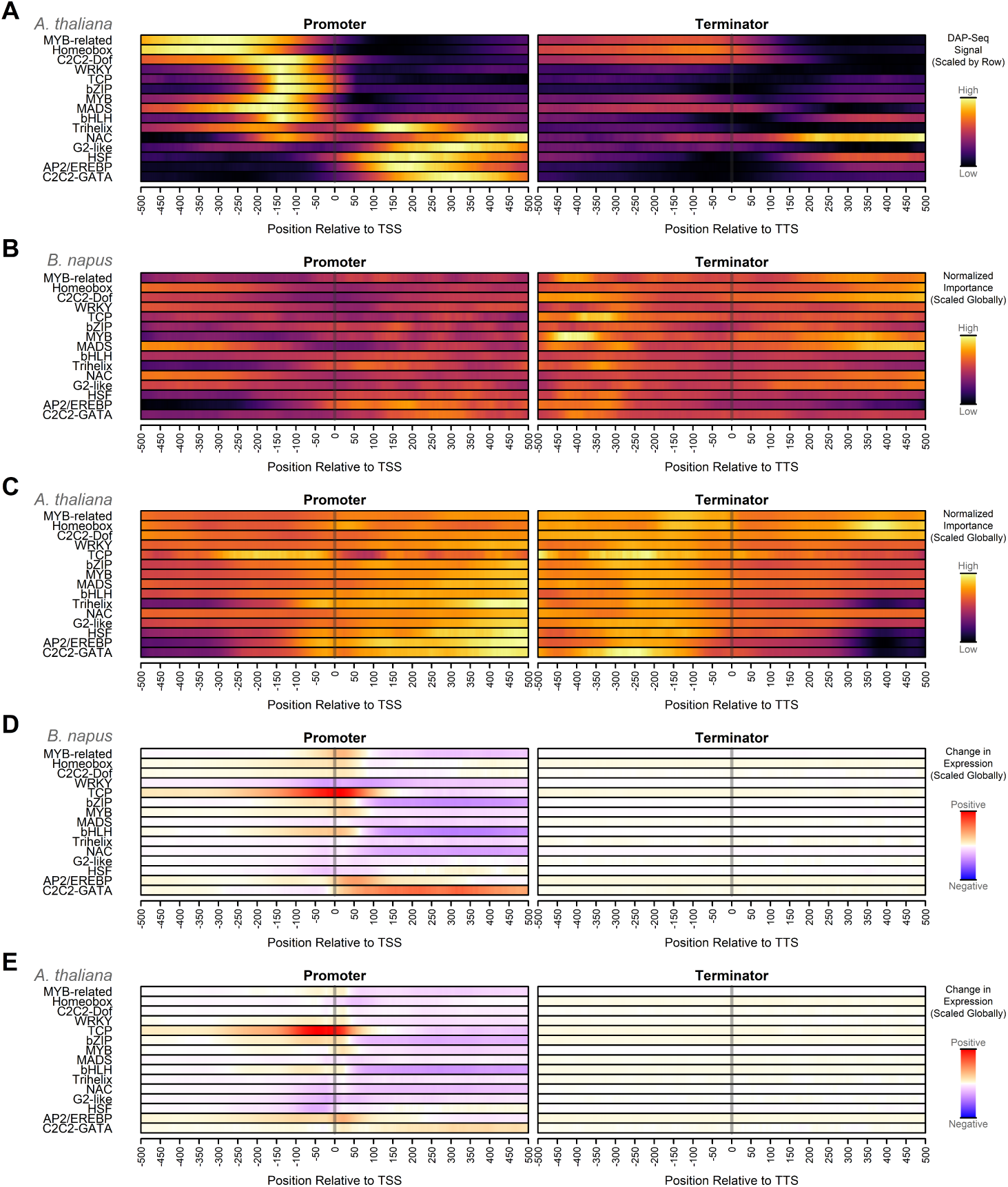
Group specific positional preferences learned by *n*emo. (A) DAP-Seq signals showing TF-binding positions for 15 TF families averaged across genes and scaled to the [0,1] range across the promoter and terminator within each TF family. (B, C) Normalized importance profiles of *n*emo_90_ for well-predicted genes in which a DAP-Seq peak of the respective family intersected within *±* 500 bp up- or downstream of the TSS, for *B. napus* and *A. thaliana* respectively. The normalized group-specific importance scores were scaled to the [0,1] range globally across all TF families and both input sequences. (D, E) Changes in expression relative to baseline (wild type), predicted by *n*emo_100_, with respect to the location of insertion of motifs of the respective TF family, globally scaled to the range [-1,1] by dividing the value at each position by the maximum absolute value across all families and both input sequences, for *B. napus* and *A. thaliana* respectively.

#### In-silico mutagenesis suggests position-specific effect of some TF families

Recent experimental evidence suggest that plant and animal transcription factors can exert position-specific effect on gene expression [4, 15]. We performed in silico mutations to confirm that the model had not only learned to recognize CREs, which happened to be preferentially located in specific regions in the input, but also learned to associate specific motifs and their effect with specific regions. We inserted motifs of each family into random locations in the input sequences of moderately expressed genes whose promoters did not intersect with a DAP-Seq peak of the respective TF family, and measured how the predicted expression changed with respect to the location of the insertion (Fig. 4D,E, Fig. S11-S40). We observed distinct patterns for the various TF families that were mostly consistent between the two species.

To our knowledge, C2C2-GATA TFs are the only family with extensive experimental validation demonstrating a position-specific effect (expression upregulation only if found downstream of TSS) across a number of species [15]. Concordantly, our in silico motif insertion analysis predicted increased expression following C2C2-GATA motif insertion, but only within a few hundred bp downstream of the TSS (Fig 5D,E). A strong positive change in predicted expression was observed when TCP motifs were inserted around the TSS, which is consistent with experimental data pointing to TCP binding in the immediate proximity of TSSs (Fig. 5A). A positive change in expression was also predicted for AP2/EREBP motifs inserted immediately downstream of the TSS. Though the predicted effect size for C2C2-GATA and AP2/EREBP factors was lower than for TCP factors. Interestingly, for bHLH factors, which have a bimodal binding pattern, frequently binding directly upstream of the TSS and less frequently 150+ bp downstream of the TSS (Fig 5A), the model associated a positive change in expression with upstream motifs and a negative change in expression with down-stream motifs (Fig. 5D,E, S13, S14). A similar pattern was predicted for bZIP factors (Fig. S15, S16). The opposite pattern was observed for the HSF family, where insertion positions directly upstream of the TSS were associated with a decrease in expression, while insertions a few hundred bp downstream of the TSS were associated with very modest increases in expression (Fig. S25, S26). NAC factors were predicted to have a predominantly negative influence on gene expression. For MYB-related factors, the highest effect sizes were predicted downstream of the TSS, which was unexpected considering the DAP-Seq signal, suggesting a strong preference to bind upstream. A similar effect was predicted for the insertion of MYB factors. The predicted effect of the remaining TF families was either weak or inconsistent between species.

#### Epigenetics and distal cis-regulation influence prediction outcomes

While inspecting prediction results, we observed distinct groups of genes whose expression was either over- or underpredicted by our model, we hypothesized that those genes could be under additional control either by additional cis-regulatory elements or epigenetic factors. We therefore compared them with identified *B. napus* genes that are associated with enhancer elements and H3K27-trimethylation, respectively. H3K27me3 histone marks are deposited by polycomb proteins and are typically found in stably repressed genes [44]. Super-enhancers (SEs) are clusters of enhancers, which can increase the expression of cognate genes over long distances and typically have a strong effect on surrounding genes [45]. To assess the effect of these factors on prediction errors, *B. napus* genes were categorized based on the magnitude and directionality of their prediction error (predicted expression − observed expression). Up to an error of ± the median absolute error across all genes (Fig. 6C), predictions were considered good. If the error was above this range, the respective gene was considered overpredicted. If it was below this range, it was considered underpredicted (Fig. 6A). By definition, well-predicted genes make up 50% of the entire data set. The other 50% are distributed among the other two error-based classes.

**Fig. 6.**
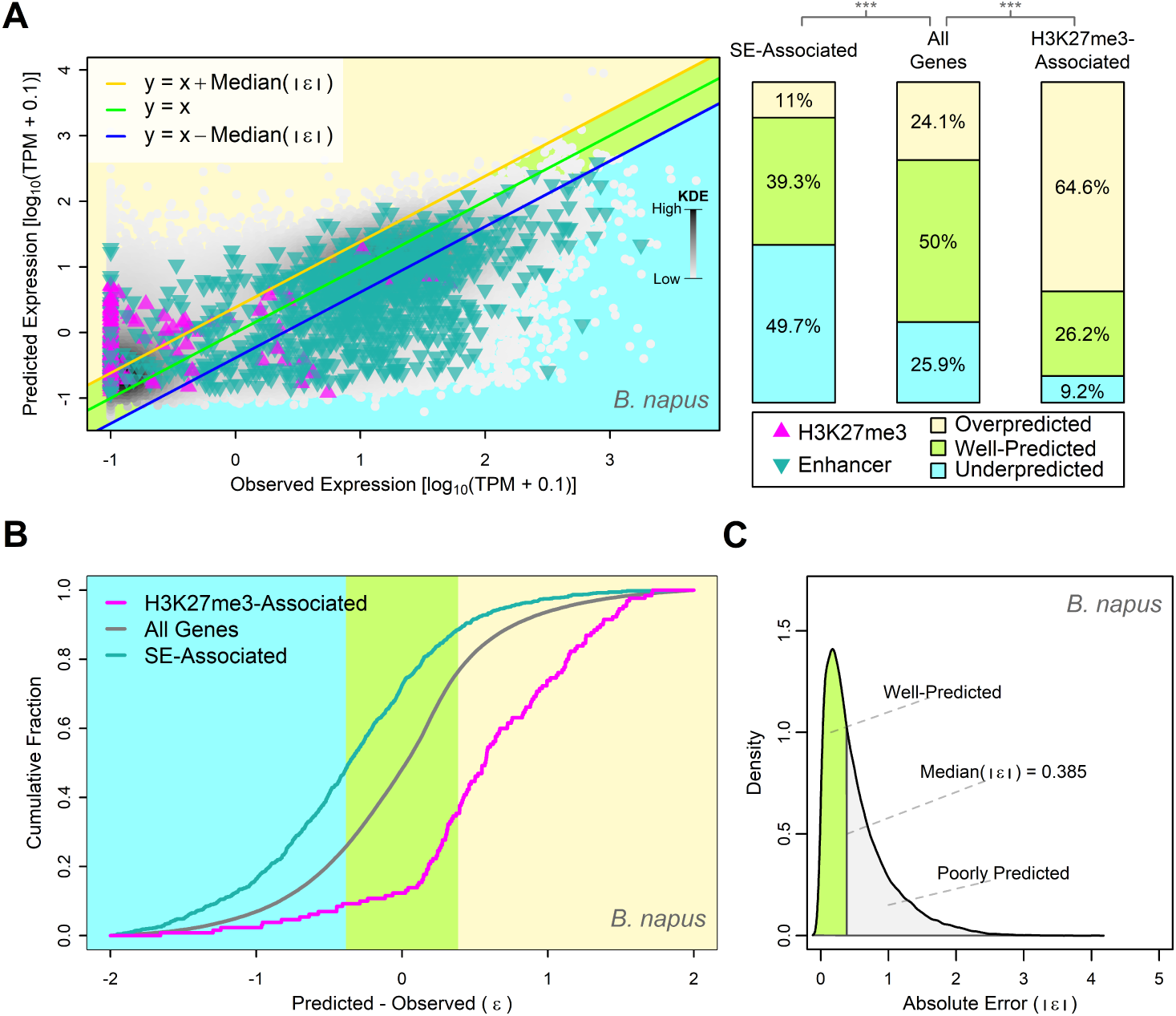
Regulatory features not captured by the model. (A) Relative proportions of prediction error-based gene classifications for genes associated with H3K27me3 histone marks or near-by SEs, compared to the overall distribution. (B) Empirical cumulative distribution function of prediction errors for H3K27me3-associated genes, SE-associated genes and all genes respectively. (C) Distribution of absolute prediction errors.

When comparing the overall distribution of the error-based classes with the distributions of genes associated with SEs and H3K27-trimethylation, there was a clear enrichment of underpredicted genes among the subset of genes that were associated with SEs (*χ*^2^-test, *N*_1_ = 956, *N*_2_ = 66, 825, *p <* 2.2 · 10^−16^) and an enrichment of overpredicted genes among genes associated with H3K27me3 histone marks (*χ*^2^-test, *N*_1_ = 130, *N*_2_ = 66, 825, *p <* 2.2 · 10^−16^). Importantly, the SE-associated genes [46] and underpredicted genes (Fig. S41) were no more tissue-specific than well-predicted genes, suggesting that the prediction error for underpredicted genes was not a result of more tissues-specific expression patterns. This suggests that the model was not able to properly predict the influence of epigenetic and potentially more distal and complex cis-regulatory features. This could be because the enhancer elements were found too far away or the fact that the model clearly focused on features proximal to the TSS and TTS, respectively (Fig. 4B). The outside influence of enhancers may lead to expression levels that are higher than local regulatory features would suggest, leading the model to predict lower expression levels than what is observed. The opposite of this applies to H3K27me3 histone marks, where, based on features in the local DNA sequence, the model may overestimate the expression of genes that are epigenetically silenced.

## Discussion

Deciphering the gene regulatory architecture of plants remains a challenging task, yet deep learning models are remarkably adept at identifying relevant patterns and predicting gene expression from genomic sequence. We were able to show that a well-optimized CNN model is able to explain more than 50% of the variation in median gene expression, even in complex polyploid plants such as *B. napus*, using only proximal non-coding sequence information. We have therefore created a single CNN model that is able to match the performance of a recently published transformer-based architecture for gene expression prediction from proximal sequences in plants, as well as an ensemble architecture consisting of 27 CNNs from the same study [38].

Besides being a versatile and economically important crop, *B. napus* is also exceptionally well-suited for the study of complex polyploid genomes, due to its close evolutionary relationship with the model plant *A. thaliana* and many other important crops in the mustard family (Brassicaceae), as well as its very recent history of polyploidization [47, 48]. However, a challenge in applying deep learning to polyploid crops is the high degree of sequence redundancy. Not only does sequence redundancy complicate the acquisition of high-quality training data due to multi-mapped sequencing reads [49–51], but it can also lead models to overfit on irrelevant evolutionary artifacts if homoeologous and paralogous sequences are not properly accounted for when partitioning data into training, validation and test sets [28]. A better understanding of gene regulation in the context of polyploid genomes is of high economic importance, due to their prevalence among major crops. We have tackled the issue of sequence redundancy in four major ways. (1) We used the EAGLE-RC pipeline [50] to obtain accurate homoeolog-specific expression values. (2) We discarded homoeologs with identical CDS sequences from the data set, due to the impossibility of obtaining accurate expression values from short RNA-Seq reads. (3) We used gene family-guided data partitioning based on CDS similarity [28, 52], to minimize evolutionary relationships between data sets used for training, validation and testing. (4) Since homology-based methods depend on arbitrary thresholds, we hard-masked coding sequences to further minimize any potentially remaining dependencies that the model could exploit. The decision to mask coding sequences is supported by the observation that, while it marginally reduces model performance, it also reduces the difference in performance between gene family-guided partitioning and random partitioning (Fig. S43). In doing so, we hope to establish best practices for training deep learning models on data from polyploid species.

By randomizing the promoter and terminator inputs, respectively, we found that despite using a highly flexible architecture that can learn separate motifs and patterns for the promoter and terminator, most of the model’s performance depended on the promoter input alone. However, since the 3’ UTR has been hypothesized to be important in fine-tuning gene expression levels [28], this finding may not hold up for predictions of tissue- or condition-specific gene expression. Although it is possible to train performant regression models for tissue-agnostic gene expression across species [38], they are ultimately limited to general patterns that are transferable between species. Species-specific models have the potential to yield insights that are more relevant to the species of interest. Furthermore, even though general regulatory patterns may be similar, the structure and function of tissues and the adaptive strategies to environmental cues may differ significantly between species, making it difficult to train condition- or tissue-specific models across species. The proportion of variation in gene expression that can be explained based on proximal sequences seems to be comparable, or perhaps somewhat lower in plants than in mammals [30]. Although accessible chromatin regions in plants are highly enriched in close proximity to genes [12] and chromatin interactions seem to mostly occur locally [40], proximal gene regulation does not appear to be much more prevalent in plants compared to mammals, where 75% of open chromatin regions are located more than 10 kb away from genes [13].

Nevertheless, by calculating importance scores and devising a normalization strategy to highlight group-specific regional differences in importance, we were able to partially recapitulate the proximal positional binding preferences of TF families. Genes regulated by specific TF families showed specific regional importance patterns that were largely reproducible between models trained on data from *B. napus* and *A. thaliana*, respectively, and roughly coincided with transcription factor binding profiles gathered from DAP-Seq data in *A. thaliana*. Interestingly, we observed that TFs, whose effect appears strongly upregulating, tend to bind downstream of TSSs (Fig. S10), which may be the reason for the high importance of UTR regions implied by the deep learning models. Since CRMs typically contain multiple CREs that bind specific combinations of TFs, single motifs may not be predictive of gene expression outcomes [15]. Therefore, it is not surprising that other regions, in addition to the observed TF-binding regions of the respective TF family, were often also important for predicting the expression of the respective genes. In addition, other regulatory patterns not related to TF-binding may have also been learned by the model.

By inserting individual TF binding motifs in various locations around the TSS and TTS and measuring changes in prediction, we were able to show that the model associated motifs of the C2C2-GATA family with a positive influence on gene expression, but only if they were inserted downstream of the TSS, demonstrating that the model was able to learn biologically relevant positional dependencies that had been demonstrated experimentally [15]. This gives some credence to positional dependencies that the model associated with other TF families. Most notably, the model predicted TCP factors to have a very strong positive position-dependent influence on gene expression. While CREs binding TCP factors are highly enriched upstream of the TSS in close proximity to the core promoter, downstream binding of TCP factors is comparatively negligible [8, 53] (Fig. 5A) and likely unfavorable due to competition with or steric hinderance of the core transcriptional machinery [4, 54]. Additionally, the disruption of core promoter elements would be expected to negatively impact transcriptional activity [55–57]. This indicates that the model was only able to identify rough positional dependencies at a coarse grained resolution, as, in general, insertions around TSSs were associated with an increase in expression. Nevertheless, several studies have highlighted the extraordinarily important role that TCP factors play in various aspects of gene regulation, such as promoter strength [54], the accessibility of chromatin [32, 58] and even large-scale chromatin topology [59, 60]. TCP factors are involved in basic biological functions such as plant growth and development across many different tissues and developmental stages [61]. The identification of TCP factors as important regulators under control conditions is therefore not surprising and further demonstrates the model’s ability to identify biologically relevant features. Interestingly, the model predicted bHLH and bZIP factors to have a positive effect on gene expression when binding upstream and around the TSS and a negative effect when binding downstream. DAP-Seq signals indicate that bHLH factors bind both upstream and downstream of the TSS in a bimodal pattern (Fig. 5A) and bZIP factors are known to often interact with bHLH factors in plants [62]. The human TFs NRF1 (a member of the mammalian bZIP family), YY1 and NFY have also been shown to act as activators or repressors, depending on the binding position relative to the TSS [4].

As discussed above, single motifs are typically not predictive of gene expression outcomes, therefore, predicting the position-dependent influence of TF families by inserting single motifs comes with several limitations. For one, the consensus motifs of most TFs would be expected to randomly occur at a high frequency throughout the genome, but in vivo TF binding site selection is highly specific and typically context-dependent [63]. On the other hand, some TF families are much more diverse than others, not only in terms of recognized binding motifs, but also in terms of their influence on gene expression. While some TFs may be less reliant on specific motifs and more reliant on DNA-shape or co-regulating factors, other TFs may bind to very specific motifs, but their function depends on other co-regulators [63]. The interpretation is especially difficult for genes that are related to stress responses, since they would often be expected to be lowly expressed under control conditions. Therefore, the motifs of TF families associated with stress responses may also be associated with low expression by the model, even if the members of the respective families primarily function as activators. Since the model was trained to predict median gene expression, it is unsurprising that the dispersion of prediction errors was lowest for genes with low tissue-specificity (Fig. S41C,D). Tissue- or condition-specific control of gene expression often depends on more complex regulatory mechanisms involving distal or epigenetic factors [64], which may be difficult for this type of model architecture to capture. Despite these limitations, we found that the errors made by the model can serve as a valuable source of information for identifying genes under complex regulation. We found that silenced genes tended to be overpredicted, whereas enhanced genes tended to be underpredicted (Fig. 6A,B). The magnitude and directionality of errors made by the model can therefore give useful hints towards the mechanisms underlying the regulation of any given gene.

## Conclusion

Highly optimized sequence-to-expression models are useful and versatile tools in the field of regulatory genomics. They can be used to independently validate and complement experimental findings and generate new hypotheses. We found that proximal gene regulation does not appear to play a greater role in plants than in animals, despite published differences in chromatin organization that might suggest otherwise. We further demonstrated that CNNs can not only recognize relevant motifs, but are also able to capture the spatial context of motifs in relation to their function. We were able to show that genes that may be subject to more complex regulatory mechanisms can be identified by comparing model predictions with observed expression. Having successfully trained a state-of-the-art model on the allotetraploid *B. napus* genome, we hope to lay the foundation for future studies of deep learning in complex crop species, which may be used to elucidate condition- or tissue-specific gene regulation and uncover the genetic and epigenetic factors underlying phenotypic variation and trait determination. The availability of robust species-specific models that can be probed and used to test the effects of mutations at low cost is invaluable for precision genome editing efforts. Far from being impenetrable black boxes, we have demonstrated their usefulness for generating hypotheses and gaining novel insight into gene regulation. In the future, we anticipate such models to aid in the creation of the next generation of resilient crop varieties that are capable of ensuring global food security.

## Methods

### Genomic data

The *B. napus* Express617 v1 assembly [65] was used as the genomic reference. A structural genome annotation containing UTR information was generated using Helixer [66, 67] with the land plant v0.3 a 0100 model. Gene models were filtered using Gffread v0.12.7 [68] with the options -CJV, to retain only protein-coding genes with a valid open reading frame. Genes located on unplaced scaffolds were excluded from the data set.

For *A. thaliana* nuclear chromosomes of the TAIR10 genome assembly [69] were used together with the TAIR10 genome annotation [70], which was likewise filtered for protein-coding genes.

### Gene expression data

For *B. napus*, a total of 36 RNA-seq samples from the cultivar Express617 with 1 to 3 biological replicates each, covering 8 tissues at various developmental stages, were used to quantify gene expression (Tab. S1). The RNA-Seq data included samples from 2 studies [71, 72], as well as thus far unpublished samples.

The reference genome was divided into subgenome-specific references containing only chromosome-scale scaffolds of the respective subgenome. Adapters were trimmed from RNA-seq reads using trimmomatic v0.39 [73]. Trimmed reads were then aligned to each subgenome using STAR v2.7.10b [74] with options –outSAMstrandField intronMotif, –outFilterIntronMotifs RemoveNoncanonicalUnan-notated, –outSJfilterCountUniqueMin 3 2 2 2, –outMultimapperOrder Random, and –outFilterType BySJout. The resulting BAM files were further processed using the EAGLE-RC pipeline [50] to obtain the most likely mapping locations, using Gffread v0.12.7 [68] to extract transcript sequences, LAST v1447 [75] to call subgenome-specific variants, and EAGLE v1.1.3 [76] for subgenome-specific read assignment. Transcripts per million (TPM) were estimated for each gene using Stringtie v2.2.1 [77]. Expression values were averaged across biological replicates within each sample before calculating the median expression across all 36 samples. Non-expressed genes, which had a maximum expression of 0 across samples, were excluded from the data set. Likewise, highly similar homoeologs of *B. napus* (100% identical coding sequence) were removed, due to difficulties in obtaining accurate gene expression values, since RNA-Seq reads that cannot be unambiguously assigned to a specific homoeolog are discarded in the EAGLE-RC pipeline.

For *A. thaliana*, a precomputed TPM-normalized gene-level expression matrix for the Araport11 genome annotation, containing data from 56 RNA-seq experiments covering various tissues, developmental stages and growth conditions, was obtained from the EMBL-EBI Expression Atlas [78, 79]. TAIR10 gene models were used instead of Araport11 gene models, as TAIR10 had been used to calculate TF binding-site distributions with respect to the TSS and TTS by Voichek et al [15] and TAIR10 transcripts have substantially smaller UTRs on average. Median expression values were calculated across 54 of 56 samples. Two samples were discarded due to low spearman correlation with the other samples (Fig. S2). Non-expressed genes (max. TPM of zero across the 54 included samples) were discarded.

### Plant growth, RNA extraction and sequencing

#### Time series data (leaf, apex, floral organs)

Brassica napus cv. Express617 was sown in cereals mix (40% medium grade peat, 40% sterilized soil, 20% horticultural grit, 1.3 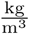 PG mix 14-16-18 + Te base fertilizer, 1 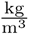 Osmocote Mini 16-8-11 2 mg + Te 0.02% B, wetting agent, 3 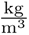 maglime, 300 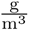 Exemptor). Material was grown in a Conviron MTPS 144 controlled environment room with Valoya NS1 LED lighting (250 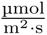) 18°C : 15°C, day : night, 70% relative humidity with a 16 h day. At day 21, plants were placed into vernalizing conditions at 5°C (8 h day) for 6 weeks before return to warm, long day conditions. Plants were sampled at day 7 (seedling) 21 (3 leaf stage, pre-cold treatment), day 42 (4-5 leaf stage, 3 weeks - mid vernalization treatment), day 63 (5 leaf stage, 6 weeks vernalization treatment), day 64 (5 leaf stage, one day returned to warm, long day), day 71 (8-9th true leaf, apex vegetative) and day 98 (BBCH51, floral buds present). At each sampling timepoint three replicates samples containing either three whole seedings (day 7) or three dissected apices were collected. One leaf sample was collected at each timepoint. Sampling of bud and anther stages was performed at 131d post sowing. Three replicate samples of three buds from a single inflorescence were collected at Sanders stages 6-7 (Pollen Mother Cell (PMC) meiosis and tetrad stage), Sanders stages 10-11 (mitosis, polarised microspore to bicellular stage). Three replicates of anthers and gynoecia were collected comprising the largest two unopened buds and the next opening flower per inflorescence for two inflorescences per replicate (36 anthers and six gynoecia in total per replicate). Samples were ground in LN2 to a fine powder before RNA extraction and DNase treatment were performed following the method provided with the EZNA^®^ Plant RNA Kit (Omega Biotek Inc., http://omegabiotek.com/store/). RNA samples were processed at Novogene (Beijing, China). Complementary DNA (cDNA) libraries were constructed using NEBNext Ultra Directional Library Kit (New England Biolabs Inc., Ispwich, MA, USA). Sequencing was performed using Illumina HiSeq X, resulting in 150 bp paired end reads.

#### Broad tissue data (leaf, root, seedling, silique, immature flower bud)

*Brassica napus* cv. Express617 was grown under long day conditions in growth chambers and vernalized for 10 weeks at a constant 5°C. Five sample types were selected to represent thee vegetative and two reproductive growth stages: leaves, roots, seedlings, immature flower buds and siliques. RNA was extracted with Quick-RNA™ Plant Miniprep (Zymo Research), quantity and purity were assessed with Qubit RNA BR Assay Kit and Nanodrop (ThermoFisher Scientific), RNA integrity with the RNA 6000 Nano Kit for 2100 Bioanalyzer Systems (Agilent). Samples were sent to Novogene for library preparation and sequencing with strand specific Illumina 150PE. For further information see [46].

### DNA affinity purification sequencing data analysis

DNA affinity purification sequencing (DAP-Seq) narrow peaks for various TFs of 15 TF families (MYB-related, Homeobox, C2C2-Dof, WRKY, TCP, bZIP, MYB, MADS, bHLH, Trihelix, NAC, G2-like, HSF, AP2/EREBP, C2C2-GATA) were obtained from NCBI GEO [8, 80, 81]. Peak coverages along the *A. thaliana* TAIR 10 reference assembly were calculated using the coverage tool of Bedtools v2.30.0 [82] with the -d option and summed across TFs of each respective family. For each TF family, peak coverages within promoter regions (TSS ± 500 bp) and terminator regions (TTS ± 500 bp) were summed across all primary TAIR 10 transcripts. Orthologs in *B. napus* were identified based on protein similarity using BLASTp v2.12.0 [83].

### Identification of genes associated with super enhancers and H3K27-trimethylation

SE-associated genes in B. napus were identified using SE locations provided by Zanini et al. [46] and applying the function bedtools closest (v2.30.0). A set of Polycomb-silenced genes in *A. thaliana*, which are associated with H3K27me3 histone marks, were taken from the literature [84]. Orthologs in *B. napus* were identified based on protein similarity using BLASTp 2.12.0 [83].

### Data preprocessing

#### Partitioning of genes based on CDS similarity

Genes were partitioned into ten evolutionarily independent sets of roughly equal size using GraphPart v1.0.2 [52], which implements gene family-guided partitioning [28]. Genes were classified into one of three classes based on their expression levels, using the upper and lower quartile as boundaries to categorize genes as lowly, moderately, or highly expressed. These classes were used to guide partitioning, ensuring that the distribution of expression levels was roughly equal between data folds. GraphPart uses sequence similarity between genes to determine their optimal partitioning to reduce evolutionary dependencies between sets, which could otherwise lead to overinflated performance estimations due to the model being able to overfit to evolutionary artifacts [28]. Partitioning was based on CDS similarity using GraphPart’s mmseqs2needle function and the options -th 0.3 -re 0.2 -nu -tr.

For hyperparameter optimization and model performance evaluation, *B. napus* genes were partitioned by themselves. For cross-species comparisons of model performance and the calculation of attribution scores, *B. napus* and *A. thaliana* genes were partitioned jointly, using six classes based on all possible combinations of expression level and organism of origin to guide partitioning, so that proportional numbers of genes for each expression level and organism landed in each joint data fold. Afterwards, genes were split back up, based on the respective organism of origin, resulting in two sets of data folds, one for each organism, which evolutionarily correspond between organisms.

#### Sequence data

CDS features were hard-masked from the reference assemblies using the maskfasta tool from Bedtools v2.31.1. This was done to further mitigate potential overfitting due to spurious correlations between homologs that might have been missed by GraphPart.

For each gene, the TSS and TTS were identified as the outer boundaries of the outermost annotated exons of the longest annotated isoform. Sequence intervals ranging from -15 kb to +5 kb around the TSS and -5 kb to +15 kb around the TTS were extracted from the respective reference assembly. Sequence intervals that extended beyond the outer boundaries of the respective chromosome were padded with Ns. For genes annotated on the minus strand, the reverse complement sequence was extracted accordingly.

#### Xpresso gene features

To train the Xpresso model [30], eight additional features (lengths and GC-contents of 5’ UTRs, 3’ UTRs and total CDS, the total intron length and the number of CDS-exons per kilobase of CDS sequence) were extracted from the reference assembly and genome annotation.

#### Data scaling

The median expression data, as well as all the numeric input features for the Xpresso model, except for GC-content, were log_10_ transformed after adding an offset of 0.1.

For hyperparameter optimization and performance evaluation, two data folds were reserved as test and validation sets. In order to prevent information leakage, a standard scaler was fit only to the numeric data in the eight remaining data folds, which served as the training set, before applying it to all data folds. For cross-species comparisons of model performance and the calculation of attribution scores, each data fold was held out once as a test fold, while training a model instance on the remaining nine data folds, respectively. This strategy resembles a 10-fold cross-validation scheme, as is often used in machine learning [85]. This was possible because the training regimen was fixed to six epochs on a fixed learning rate schedule, which obviated the need for a separate validation set that would typically be used to stop model training based on the validation loss. Accordingly, standard scalers were fitted to the corresponding nine training folds, preventing data leakage to the respective test fold. For the final predictive model, a standard scaler was fitted to the entire data set.

### Hyperparameter optimization

Hyperparameters were optimized in a multi-step process using Bayesian optimization over large search spaces with Optuna v3.6.0 [86]. In the first set of steps, the hyperparameters that govern the general model architecture were tuned using validation loss (mean squared error) as the performance metric. In subsequent steps, the hyperparameters that govern the learning algorithm itself, including the loss function, were optimized, using the coefficient of determination (*r*^2^) on the validation set as the performance metric. Hyperparameters were optimized with respect to a single data-fold configuration, setting fold 0 aside as the test set and not using it at all during hyperparameter optimization. Fold 1 was used as the validation set to evaluate model performance for different hyperparameter settings. The remaining eight folds were used to train the model in each round of optimization.

(1) As a first step, the hyperparameters governing the overall architecture of the model were optimized using the Adam optimizer [87] with default parameters and selecting hyperparameters that minimize the mean squared error on the validation set. The hyperparameter search space for the model architecture (Tab. S2) was mostly a superset of published architectures that were used successfully for similar tasks [28, 30, 88]. In addition, the search space included convolutional strides larger than 1, overlapping pooling windows, average-pooling and a large array of activation functions. Overlapping pooling windows (pooling strides smaller than the window size) were used in an effort to improve the ability of the network to resolve the spatial organization of motifs by increasing the overlap of receptive fields, thus mitigating the occurrence of multifaceted neurons [25, 89]. In the case of image classification, the use of overlapping pooling windows has also been reported to increase performance and reduce overfitting [22, 90]. Features from promoter and terminator sequences were extracted in separate convolutional branches, so that convolutional kernel sizes could be optimized independently during hyperparameter tuning and the corresponding model weights could be optimized independently during training. This was done to allow for some flexibility between branches, since relevant features may differ between promoter and terminator sequences. However, this flexibility comes at the cost of an increased number of model parameters. Pooling window sizes, pooling strides and the number of convolutional filters in the last convolutional layer were kept equal between both branches to ensure that their respective output feature spaces have equal dimensions.
(2) A selection of hyperparameters found in the first round of optimization, related to the convolutional layers, was validated by switching hyperparameter values between the two convolutional branches or using default values instead (Tab. S3). The hyperparameters that had been found for the promoter and terminator branches, respectively, were tested in all possible combinations, using them for either or both branches. The best performance was seen when the hyperparameters of the promoter branch were used for both branches. The type of pooling layer had been optimized in the first round between max-pooling and average-pooling for each individual pooling layer. To validate this decision, the optimized configuration of pooling types was tested against using only max-pooling or only average-pooling. The best performance was seen when using only average-pooling, instead of the optimized configuration. The use of convolutional strides larger than one and overlapping pooling windows was tested against the respective default. The best performance was seen when using default strides for the convolutional operations and optimized strides (smaller than or equal to the pooling window size) for the pooling operations. The architecture validation was performed by grid search, testing all possible permutations of the above-mentioned aspects.
(3) The model architecture was then fine-tuned, using insights from the first two rounds as guidance to narrow down the search space (Tab. S4).
(4) Next, the training algorithm and the loss function were optimized, keeping the model architecture constant (Tab. S5). Instead of using Adam with default parameters, in this step, Nadam [91], a variation of the Adam optimizer that uses Nesterov momentum, was used, treating the function parameters as hyperparameters to optimize. Instead of using a fixed learning rate, at this stage a one-cycle learning rate schedule was implemented, consisting of a gradual warm-up phase, followed by a high peak learning rate and a subsequent cool-down phase, allowing for fast and consistent model convergence that generalizes well, by smoothing out the loss landscape and stabilizing gradient descent [92, 93]. The length and shape of the cycle, as well as the initial, final, and peak learning rate, were optimized. The momentum was cycled similarly, but in the opposite direction as the learning rate, optimizing the maximum and minimum momentum (Fig. S4A). High peak learning rates provide a regularizing effect [94], which was balanced by setting the batch size to a value of 130, which was the highest possible batch size, given the memory constraints of the GPU. Weight decay [95] and dropout rates were optimized [96] together with the learning rate schedule [94] in order find an optimal combination of regularizing influences. The loss function itself was also optimized, using Huber loss instead of mean squared error and optimizing its delta parameter. Since loss values could not be directly compared between trials, instead the coefficient of determination (*r*^2^) on the validation set was maximized. The tuning of the Nadam optimizer and the corresponding hyperparameter search space was inspired by Choi et al. [97], who argued that more sophisticated optimizers should always perform better or equally as well as less sophisticated optimizers, given adequate parameter settings.
(5, 6) Finally, the model training was further fine-tuned in two final stages, progressively reducing the hyperparameter search spaces based on results from previous stages (Tab. S6, S7), in order to arrive at the final set of hyperparameters (Tab. S8, Fig. 1, S3).

### Model training and evaluation

Before training a given model instance, the necessary data folds were loaded into memory and numeric data were scaled using the appropriate standard scaler. The training folds were concatenated to form the training set. The order of samples within each data set was randomly shuffled, and sequences were one-hot encoded.

The models were trained on NVIDIA V100 GPUs with 32 GB of VRAM and implemented using the functional API of Tensorflow v2.14.0 [98]. Xpresso was re-implemented in Tensorflow v2.14.0 and trained on the same data as *n*emo.

#### Training for performance evaluation

For the purpose of evaluating *n*emo against Xpresso, which served as a strong baseline regression model for gene expression prediction from proximal regulatory sequences, both models were trained on the same training set used during hyperparameter optimization, encompassing 80% of the entire data set. The same validation set that had been used to assess model performance during hyperparameter optimization was used to monitor validation loss during training. Instances of *n*emo that were trained on 80% of the data are subsequently referred to as *n*emo_80_. Both models were trained 10 times on the same training data, due to the stochastic nature of the model training procedure, to capture the variability in performance across training runs (Fig. 2A).

The Xpresso model was trained using the Adam optimizer with parameters *learning_rate* = 0.001, *beta*_1 = 0.9, *beta*_2 = 0.999, *epsilon* = 10^−8^ and *weight_decay* = 0.0 and mean squared error as the loss function. Early stopping was used with a patience of 10 epochs. The model weights that had achieved the lowest validation loss were saved. The original Xpresso model architecture [30] had to be slightly altered due to issues with model convergence, likely caused by the “dying ReLU” phenomenon [99]. The authors of the original paper [30] had reported achieving convergence in 9 out of 10 training runs. For our data, the rate of convergence was much lower. Using the leaky ReLU activation function [100] with a slope of 0.1 solved the convergence issues. In addition, the Tensorflow callback function ReduceLROnPlateau was used to further stabilize convergence.

To train *n*emo, the legacy Nadam optimizer of Tensorflow v2.14.0 with parameters *beta*_2 = 0.9982, *epsilon* = 1.1330 · 10^−6^, *weight_decay* = 3.2094 · 10^−5^ was used to minimize Huber loss with *delta* = 2.9466. The learning rate and momentum (*beta*_1) were scheduled using a one-cycle policy [92, 94, 101] that reliably achieved convergence within a fixed training regimen of six epochs. The validation loss was monitored, but was not used to stop training. Since the validation set was not needed to stop the training of the *n*emo model, another set of 10 model instances was created by incorporating the validation set into the training set and training the model on 90% of the data, subsequently referred to as *n*emo_90_. Importantly, for the purpose of general performance evaluation (Fig. 2A) these instances were only trained on the 90% of *B. napus* data used during hyperparameter optimization and tested on the same unseen test set as *n*emo_80_.

#### Training for model interpretation and cross-species comparisons

Using data folds derived from the joint partitioning of *A. thaliana* and *B. napus* genes, 10 *n*emo_90_ instances were trained for each species. Each model instance was trained on a different training set, holding out one of the data folds as a test set to calculate attributions on, and training on the remaining nine data folds, similar to 10 fold cross-validation [85]. Importantly, the respective test sets remained separate from the training set of each model instance, enabling minimally biased predictions and attributions for the entire dataset when combining the predictions of all ten model instances on their respective test sets. Due to the joint partitioning of data between species, corresponding data folds contained homologous gene families (e.g. fold 0 of *A. thaliana* contained gene families related to genes in fold 0 of *B. napus*, but practically unrelated to genes in all other data folds), therefore corresponding test sets could be used interchangeably between species. Apart from the cross-species comparison shown in Fig. 2B, models were always trained and tested on the same organism.

#### Training a final predictive model for mutational analyses

A final instance of *n*emo was trained on the entire data set, to release the model weights as a tool to make predictions on other genotypes and other *Brassica* crops. This model instance, subsequently referred to as *n*emo_100_, was used to predict position-dependent effects of inserting TF motifs into wild-type promoter and terminator sequences.

#### Model predictions

For each trained model instance, predictions were made only on held-out test data, complementary to the data on which the respective model instance was trained on. The test set used to evaluate general model performance (Fig. 2A) was not used at any point during hyperparameter optimization or model training and contained gene families never before seen by the model. To assess the contributions of promoters and terminators to the total prediction, the same *n*emo_80_ instances used for general performance evaluation were used to make predictions on mutated versions of the test set genes, where the promoter and terminator sequences were replaced by random sequences, respectively. For calculating attributions and cross-species performance comparisons based on *n*emo_90_, every data fold was held out from training once, to obtain minimally biased predictions for every gene in the entire set. Furthermore, for the cross-species comparison, the *B. napus* test sets were randomly down-sampled to match the size of the *A. thaliana* test sets for better comparability, however, the training sets were not down-sampled.

Genes were defined as well-predicted if the absolute error (|*ε*|) was lower or equal to the median absolute error across all genes (Fig. 6C). Accordingly, genes for which the signed error fell above or below the range of well-predicted genes were considered over-predicted or under-predicted, respectively (Fig. 6A).

### Attribution

Attribution scores for the input sequences with respect to the model output were calculated using the GradientExplainer from SHAP v0.42.0 [43]. This explainer implements the expected gradients algorithm [102], which is an extension of the integrated gradients algorithm [103]. It calculates robust attribution scores by sampling multiple reference sequences instead of using a single reference. For each input sequence, a set of ten reference sequences was generated by (1) splitting the sequence into fragments that contained nucleotide information and non-informative (masked) fragments that contained only Ns. (2) Sequence fragments containing nucleotide information were then shuffled 10 times using the dinuc shuffle function from DeepLIFT v0.6.13.0 [104]. The masked sequence fragments were copied ten times and (3) concatenated back together with the shuffled fragments (Fig. S5). This approach made sure that the references stay true to actual input sequences by containing contiguous stretches of shuffled nucleotides, interspersed with masked stretches that are unimportant by design. In the masked regions, all ten references were identical to the respective input, therefore, the resulting attribution was automatically zero in those regions. Since the model exhibits some degree of overfitting (higher performance on the training set compared to the validation set (Fig. S4B)), all attribution maps were calculated based on predictions made on the respective test set. For each test set, a separate *n*emo_90_ instance had been trained and used to calculate the attributions for the respective held-out test set. Attributions for all ten test sets were then concatenated together to arrive at a complete set of minimally biased attributions for each gene in the data set.

### Global and local importance scores

The sum of absolute attribution values across the 4 bases was used as a measure of local importance at a given sequence position and gene (Eq. 1). For a group of genes, the local importance scores were averaged at each position to give group-specific importance scores (Eq. 2). Global importance scores across all genes (Eq. 3) were used as a background reference value for normalization, together with the global standard deviation (Eq. 4).

Since coding sequences had been masked and attribution scores were set to zero by design in the respective regions, the number of genes in which a given sequence position was masked was accounted for when calculating averages and standard deviations of local importance scores across genes. This was done to minimize the effect that masking had on the resulting scores. Only well-predicted genes, for which the absolute prediction error was smaller than or equal to the median absolute prediction error, were used to calculate group-specific importance scores. Since attributions had only been calculated for unseen test data, it was assumed that selecting only well-predicted genes increased the likelihood that non-zero importance scores would coincide with generalizable patterns instead of artifacts from overfitting on the training set. The global importance scores, on the other hand, were calculated across all genes, including those that had been poorly predicted.

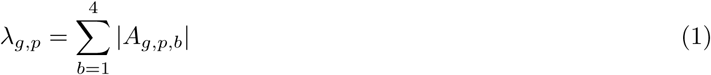

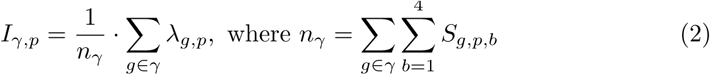

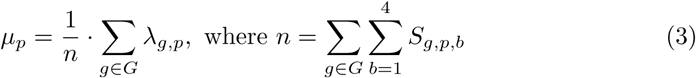

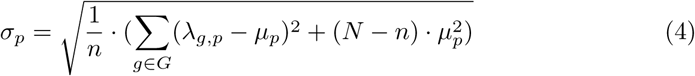

Where *A_g,p,b_*∈ ℝ is the Shapley value, measuring the attribution of base *b*, at position *p* of gene *g*, *S_g,p,b_* ∈ {0, 1} is the corresponding value of the one-hot encoded input sequence and *λ_g,p_*∈ ℝ^+^ = {*x* ∈ ℝ | *x* ≥ 0} is the local importance of gene *g* at position *p*. *I_γ,p_* ∈ ℝ^+^ is the group-specific importance of position *p* for group *γ* ⊂ *G*. *G* is the set of all genes, with *g* ∈ *G* referring to any gene in the data set. *µ_p_* ∈ ℝ^+^ is the global background importance calculated across all genes in the data set and *σ_p_*is the global standard deviation of local importance scores at position *p* across all genes in the data set. *N* is the total number of genes in the entire data set, *n* is the total number of genes that are *not* masked at position *p*, while *n_γ_* is the number of genes in group *γ* that are *not* masked at position *p*. Note that *λ_g,p_* is equal to zero at masked positions.

### Smoothing and normalization of importance scores

Due to the use of overlapping pooling windows, the input regions in which the overlap occurs are connected to a greater number of neurons in the deeper layers. Therefore, changes in these regions have a greater potential to have an impact on the output of the model. This inherent architectural bias is reflected in the importance scores as a periodic signal, with a period equal to the stride *s* of the respective pooling layer and, accordingly, a frequency equal to ^1^. A Fourier analysis of the global background importance confirmed a large peak at the expected frequency for the first pooling layer, as well as some of its harmonic frequencies, but also showed that the effect of deeper pooling layers was negligible. Applying a Savitzky-Golay filter [105] with a kernel width of 45 (three times the period of the noise signal) and a polynomial order of 1, resulted in a smooth signal that was comparable to the signal that could be achieved by directly filtering the identified noise frequencies (Fig. S6).

In order to highlight group-specific differences in the importance landscape (Fig. 5B,C) importance scores were normalized to make them more comparable between different regions in the input and different groups of genes. The normalization was done in a four-step process (Fig. S7, S8). First, Savitzky-Golay filters were applied to the global importance, the global standard deviation of local importances and the group-specific importances, as described above, using the R-package gsignal 0.3-7 [106]. Second, importance scores were mean-normalized by subtracting the mean importance across the promoter and terminator for each respective group. This was done due to the additivity of Shapley values [43], which means that the attributions of a given sample add up to the difference of the predicted value from a reference value. Due to this property, the sum of the importance scores of a given gene is correlated with the predicted expression of that gene, which, depending on the context, leads to potential bias when comparing importance scores between groups of genes that may vary in their average expression. Third, group-specific residual importance scores were calculated by subtracting the mean-normalized global importance from the mean-normalized group-specific importances. This step highlights how the importance scores of each group differ from the global background importance. Lastly, the group-specific residual importance scores were divided by the global standard deviation of local importance scores, in order to emphasize differences in areas with low variability and de-emphasize differences in areas with high variability.

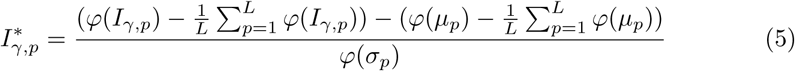

Where 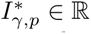 is the normalized importance of group *γ* at the sequence position *p*, *φ*(.) denotes the Savitzky-Golay filter and *L* = 12400 is the combined length of the two input sequences.

### In silico motif insertion analysis

For each TF family, the motif position weight matrix of each member of the respective family (from *A. thaliana* or *Arabidopsis lyrata*) was gathered from the JASPAR database [107]. If there were multiple versions of the motif, the latest version was chosen. The position weight matrices were converted into position probability matrices. For each TF of a given TF family, 20 motifs were generated according to the probabilities in the corresponding position probability matrix. Using these 20 motifs, 1000 randomly selected moderately expressed genes, not associated with the respective TF family, were mutated by inserting the motifs in random locations in the promoter (TSS ± 1 kb) and terminator (TTS ± 1 kb), respectively, thus creating 20,000 mutated promoters and 20,000 mutated terminators, each containing a single inserted motif. Motifs were not inserted into masked regions. Mutated promoters were then used together with wild-type terminators and vice versa to make predictions using *n*emo_100_. Baseline predictions made on the wild-type inputs were subtracted from the predictions made on the corresponding mutated inputs to arrive at changes in predicted expression, corresponding to a given motif inserted at a given position of the promoter or terminator, respectively. Changes in predicted expression resulting from insertions at each position were averaged for each TF and then across TFs for each TF family. The average changes in expression were smoothed by applying Savitzky-Golay filters of size 45 and polynomial order 1.

### Model validation with real-world data

Gene expression (RNA-Seq) data for young pre-vernalization leaves for Express617 and semi-winter oilseed rape ZS11 were obtained from public databases [41] and the BRAVO project from the John Innes Centre (JIC) [71]. To ensure maximum comparability across annotations, structural gene annotation was lifted from Express617 onto ZS11 using Liftoff v1.6.3 [108]. Gene expression was predicted using nemo. RNA-Seq data was preprocessed and quantified in the same manner as for model training. Genes were pre-filtered to only keep genes where expression in one of the genotypes was higher across all the samples. Differences in the predicted gene expression between Express617 and ZS11 was calculates and genes were sorted by this value. Top 100 and top 1000 genes where (Express 617 predicted expression) *>* (ZS11 predicted expression) and (ZS11 predicted expression) *>* (Express 617 predicted expression) were extracted and actual observed levels of expression were compared. To detect structural variants between the two ecotypes, the ZS11 genome [109] was aligned to the Express 617 genome v1 using minimap2 v2.24 [110]. Then, structural variants (SV) were called with svim-asm v1.0.2 [111]. RegioneR [112] was used to test the over-representation of SVs in extended genes bodies (1 kb upstream and downstream).

## Declarations

## Acknowledgements

This work was supported by the de.NBI Cloud within the German Network for Bioinformatics Infrastructure (de.NBI) and ELIXIR-DE (Forschungszentrum Jülich and W-de.NBI-001, W-de.NBI-004, W-de.NBI-008, W-de.NBI-010, W-de.NBI-013, W-de.NBI-014, W-de.NBI-016, W-de.NBI-022) and Justus Liebig University Bioinformatics Core Facility (BCF). The authors acknowledge the BBSRC BRAVO project (UK) as source of part of the RNAseq data use. We thank Andris Finkbeiner for suggesting the use of Savitzky-Golay filters. We thank Dr. Yoav Voichek for the constructive discussion on the possible approaches for model validation.

## Code availability

The source code for this project is available at https://github.com/GoliczGenomeLab/nemo.

## Author contribution

AAG conceptualized the project. AAG and KCR designed the project. KCR performed the data preprocessing, designed and trained the model, performed downstream data analysis, plotted the figures and drafted the manuscript. SFZ and RW performed the RNA sequencing and wrote the respective methods sections. ACT generated data used in this study. AAG contributed to the data preprocessing and performed the DAP-Seq data analysis. SFZ and AAG identified genes associated with SEs and H3K27me3. AAG, SFZ and RJM refined and substantially improved the manuscript. All authors reviewed and approved the manuscript. AAG acquired the funding and supervised the project.

