## Supplementary material for "A deep learning model captures position-specific effects of plant regulatory sequences and suggests genes under complex regulation": Fig S

### Supplemental figures 1: A deep learning model captures position-specific effects of plant regulatory sequences and suggests genes under complex regulation

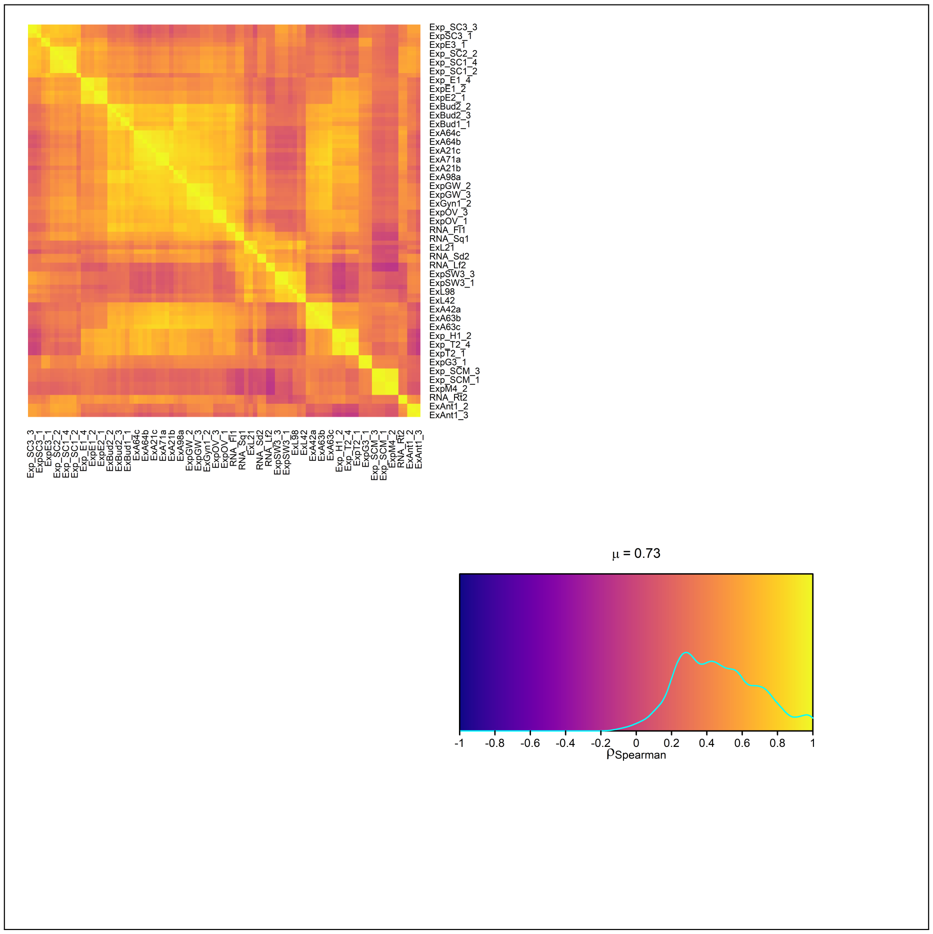


**Fig. S1.** Pair-wise Spearman correlations of TPM-normalized expression values, calculated from *B. napus* RNA-Seq samples. The 5000 most highly expressed genes across samples were used to compute correlation coefficients. The overall distribution of correlation coefficients is depicted in the bottom right.

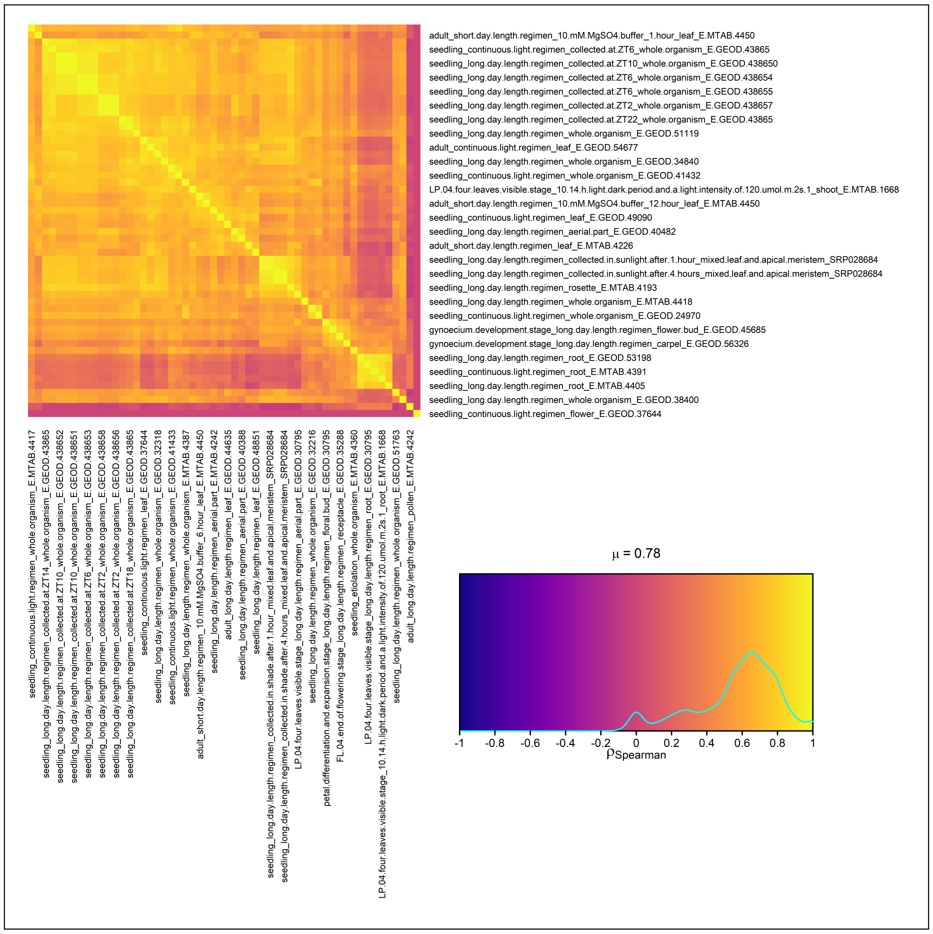


**Fig. S2.** Pair-wise Spearman correlations of TPM-normalized expression values gathered from a pre-computed expression matrix of *A. thaliana* RNA-Seq samples. The 5000 most highly expressed genes across samples were used to compute correlation coefficients. The overall distribution of correlation coefficients is depicted in the bottom right. Due to their low overall correlation, two samples ("E.GEOD.37644", "E.MTAB.4242") were discarded.

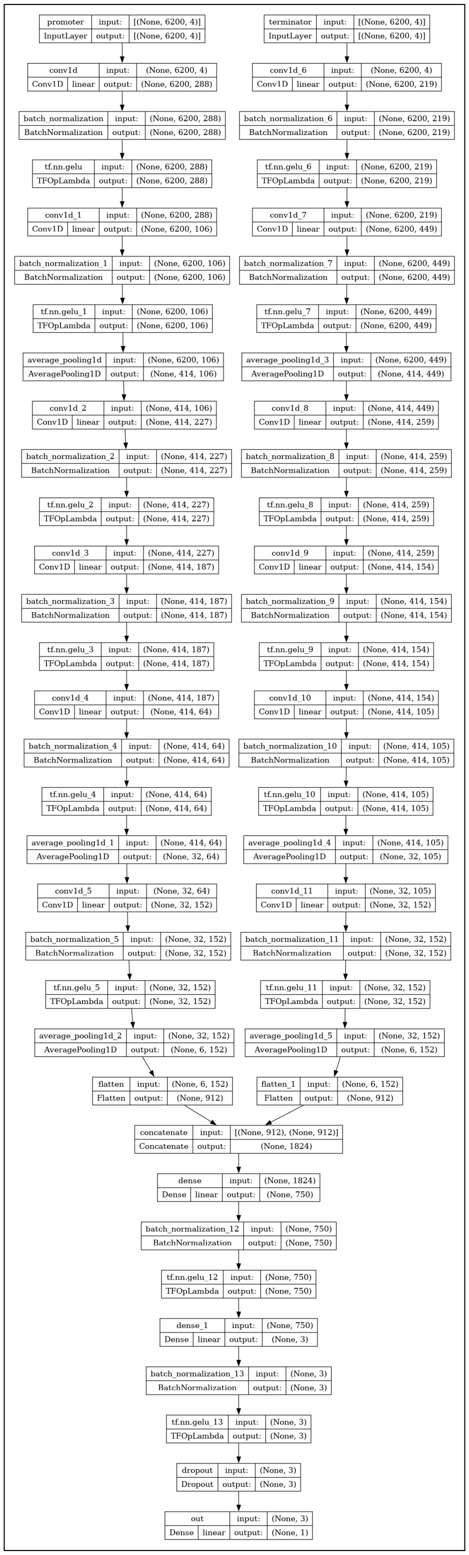


**Fig. S3.** The final n*e*mo architecture. After hyperparameter optimization, the architecture contains a stack of six convolutional layers in each convolutional branch, interspersed by batch normalization, GELU activation and average pooling. Information processing is done in two dense layers interspersed by batch normalization and GELU activation before a final dropout layer and the output.

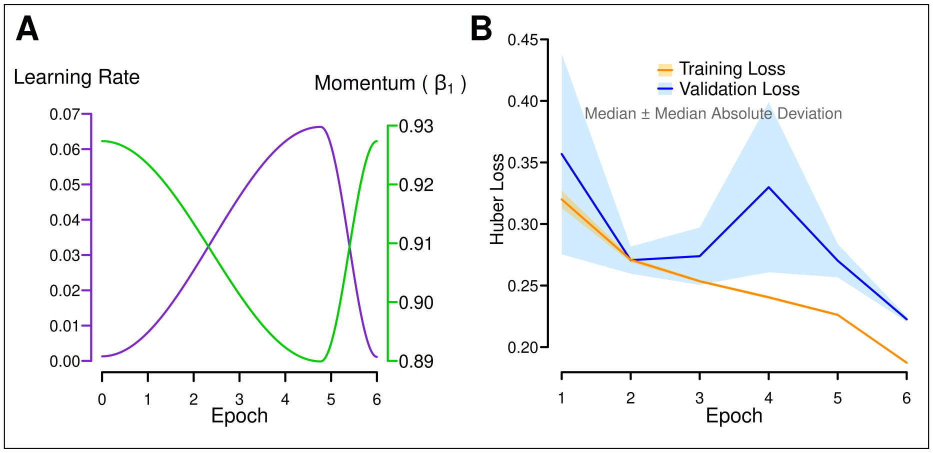


**Fig. S4.** Model training using one-cycle learning rate schedule. (A) The one-cycle schedules that are applied to the learning rate (left axis) and momentum (right axis). (B) Training and validation loss dynamics during training. Lines and shaded areas represent the median $\pm$ the median absolute deviation at each training epoch for 10 training runs using the same training set.

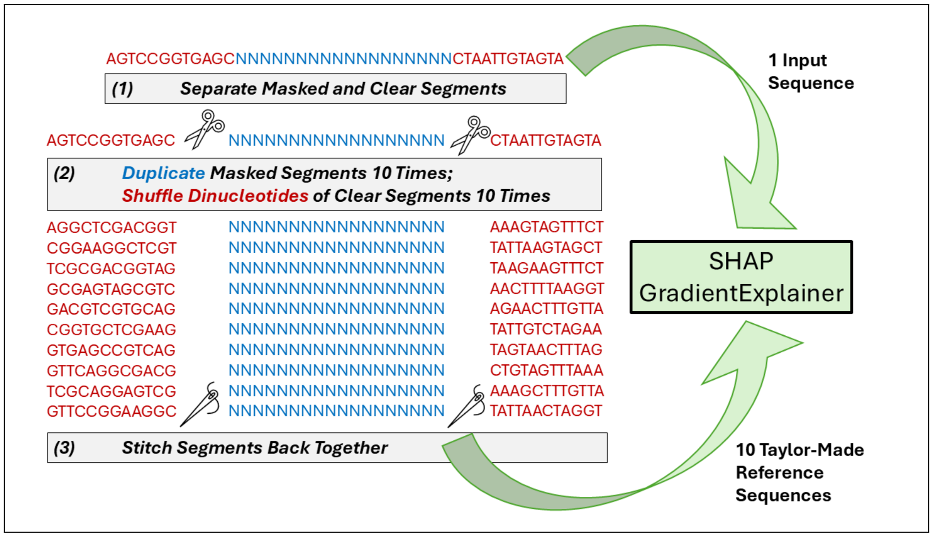


**Fig. S5.** Calculation of attribution scores. For each input sequence, a set of 10 reference sequences was randomly generated by shuffling dinucleotides of non-coding (clear) sequences. Coding (masked) sequences were maintained in the references, making them unimportant by definition. The input sequences and the corresponding sets of custom reference sequences were used to calculate attributions using the SHAP GradientExplainer.

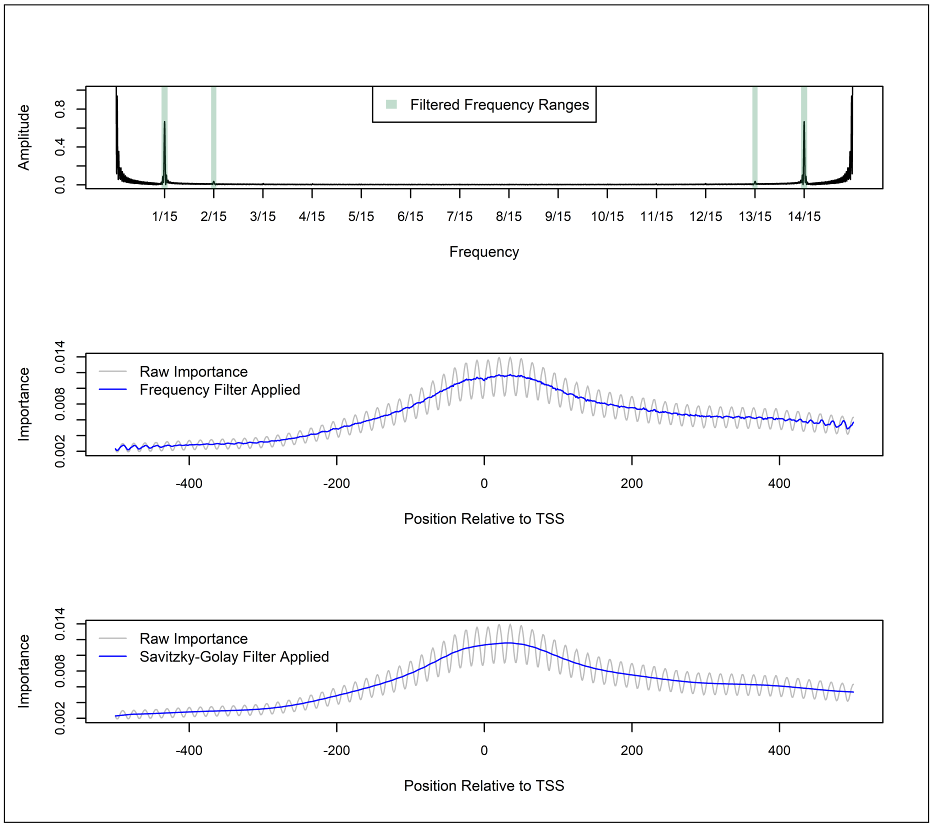


**Fig. S6.** Fourier analysis of importance scores. (Top) Fourier transformation of importance scores calculated across all genes. The periodic signal with a frequency of $\frac{1}{15}$ (plus harmonics) is caused by overlapping pooling windows with a stride of 15. Shaded areas show frequency bands filtered out by setting them to 0 (Middle), resulting in much smoother importance scores. Using a Savitzky-Golay filter with a polynomial order of 1 and a window size of 75, achieves similar results, but easier and cleaner (Bottom).

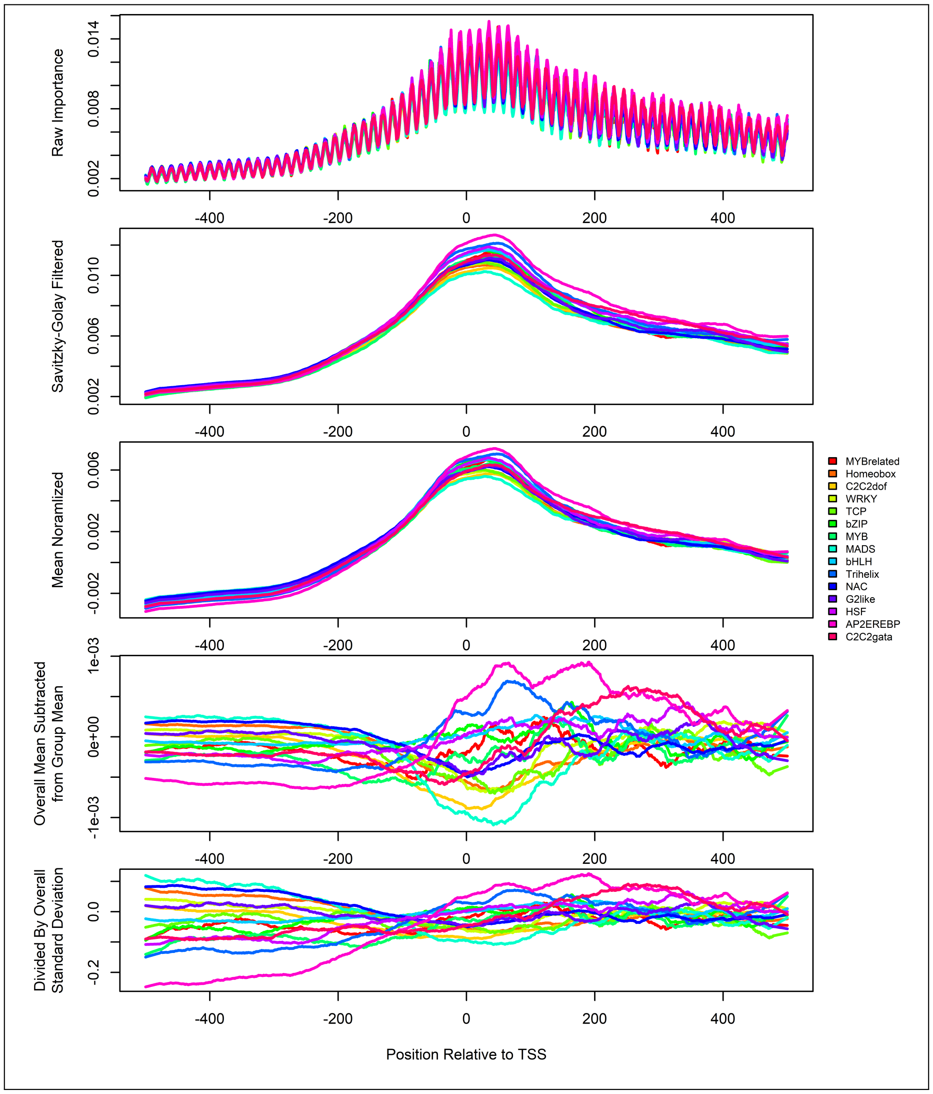


**Fig. S7.** Normalization of group-specific importance scores (*B. napus*). Savitzky-Golay filters are applied to filter out the periodic noise signal caused by overlapping pooling windows. Filtered importance scores are first mean normalized by subtracting the average importance across the sequence from the scores at each position. This mitigates bias due to differences in average expression between groups. Then, at each position, the (mean-normalized) average importance across all genes is subtracted from the (mean-normalized) group-specific importance in order to arrive at group-specific residual scores. Lastly, at each position, the residual score is divided by the standard deviation across all genes, in order to emphasize differences in less-variable regions and de-emphasize differences in highly variable regions.

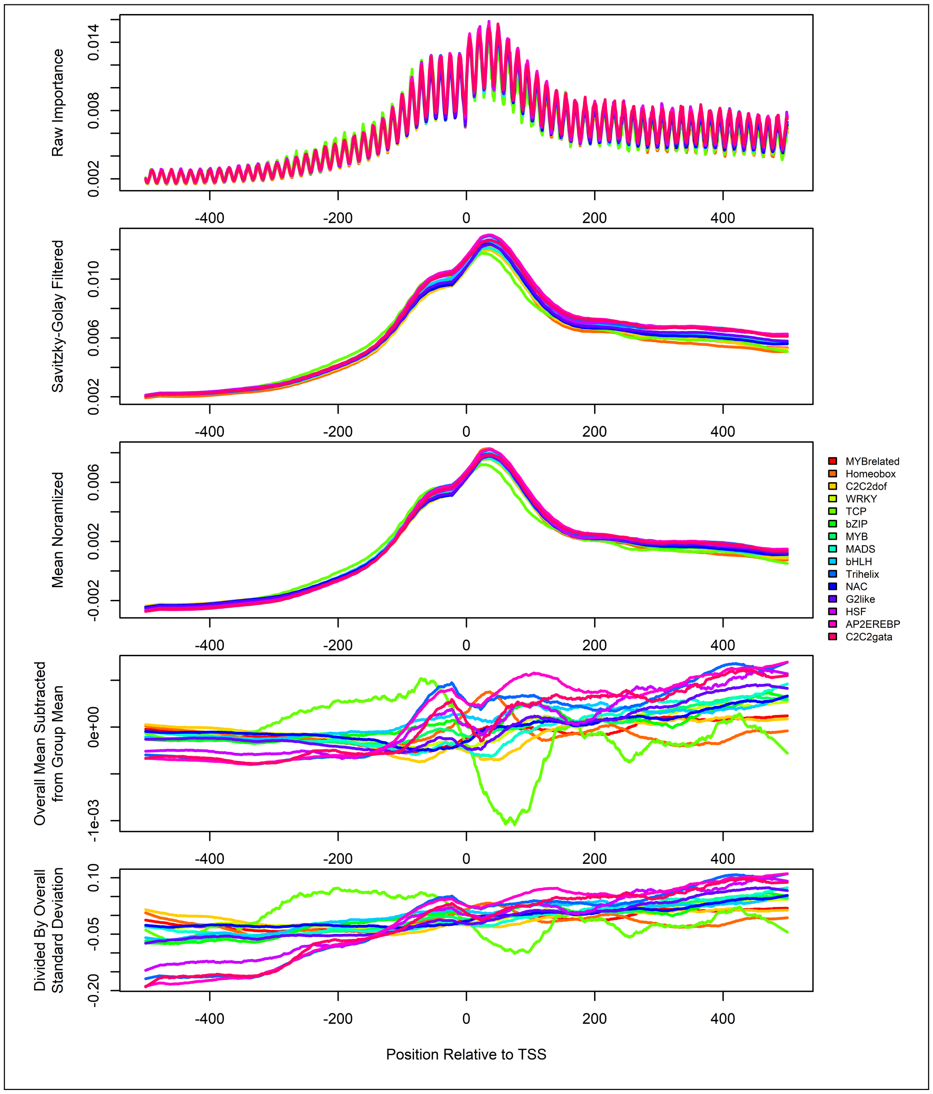


**Fig. S8.** Normalization of group-specific importance scores (*A. thaliana*). Savitzky-Golay filters are applied to filter out the periodic noise signal caused by overlapping pooling windows. Filtered importance scores are first mean normalized by subtracting the average importance across the sequence from the scores at each position. This mitigates bias due to differences in average expression between groups. Then, at each position, the (mean-normalized) average importance across all genes is subtracted from the (mean-normalized) group-specific importance in order to arrive at group-specific residual scores. Lastly, at each position, the residual score is divided by the standard deviation across all genes, in order to emphasize differences in less-variable regions and de-emphasize differences in highly variable regions.

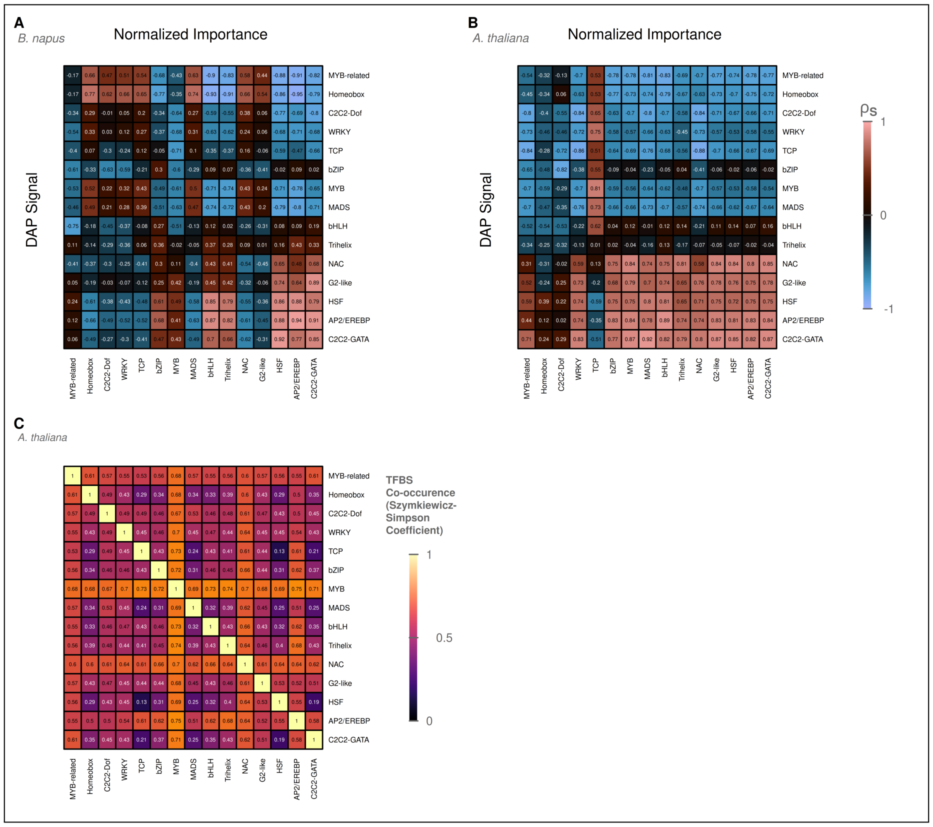


**Fig. S9.** TF co-occurrences and importance score correlations. (A, B) Spearman correlations between DAP-Seq signals and normalized (group-spceific) importance scores for *B. napus* (A) and *A. thaliana* (B). The interval ±500 bp around the TSS was used to calculate correlation coefficients. (C) Co-occurences of TFBSs measured by the Szymkiewicz-Simpson coefficient based on DAP-Seq peaks from different TF families overlapping the same promoter region. DAP-Seq data was only available for *A. thaliana*.

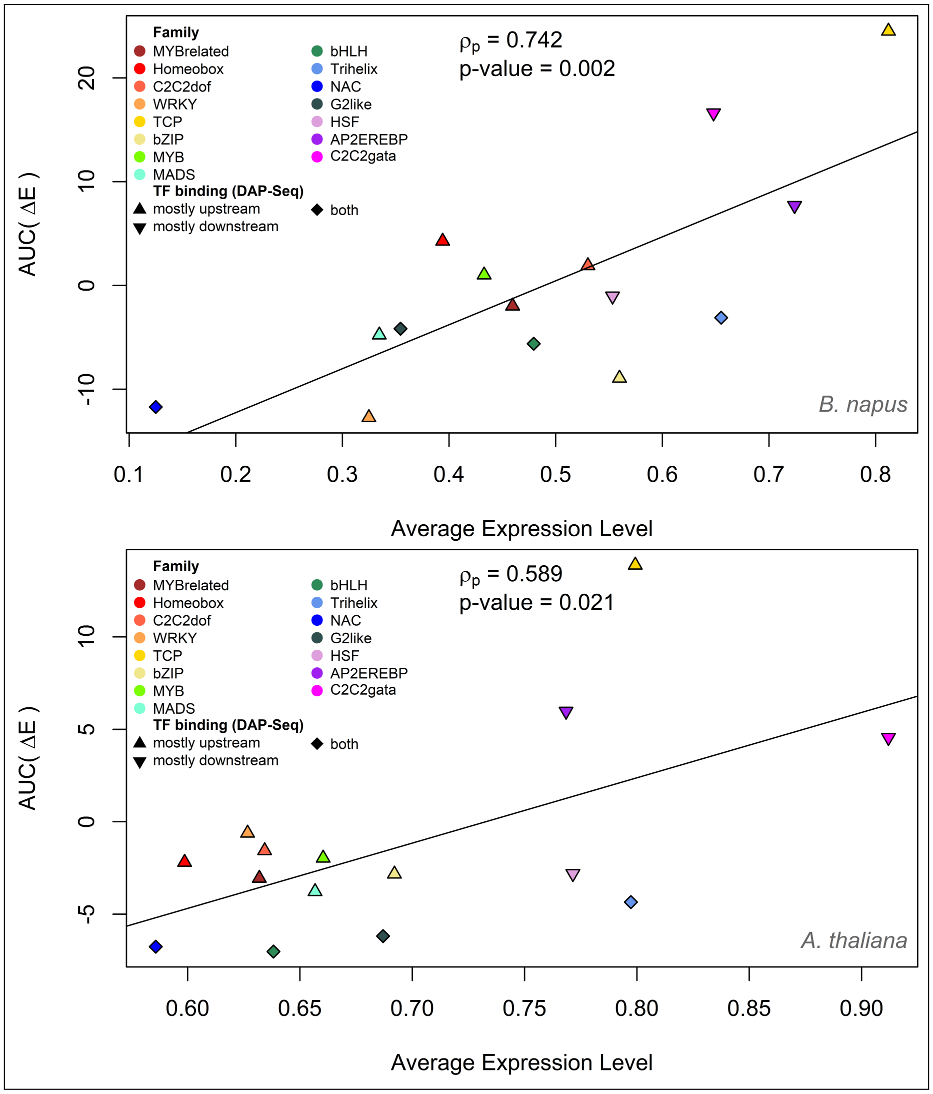


**Fig. S10.** Correlation of expression levels and response to motif insertion. For each group of genes associated with a particular TF family, the area under the curve of average predicted changes in expression across the interval spanning ±500 bp around the TSS plotted against the average gene expression of genes in the respective group. TF families are qualitatively grouped with respect to their binding preference relative to the TSS, based on visual inspection of DAP-Seq signals.

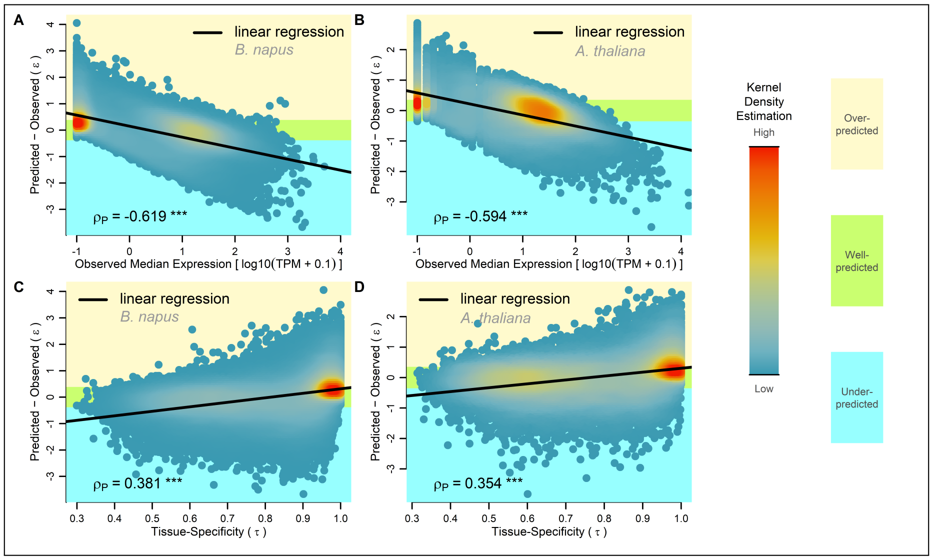


**Fig. S41.** Prediction-based categorization of *A. thaliana* genes. (A) Genes are classified as over-, under- and well-predicted based on the prediction error, using the median absolute error as the threshold. Predictions were made for the entire data set using via cross validation. The coloration of points is based on Gaussian kernel density estimations. (B) Overall distribution of absolute prediction errors.

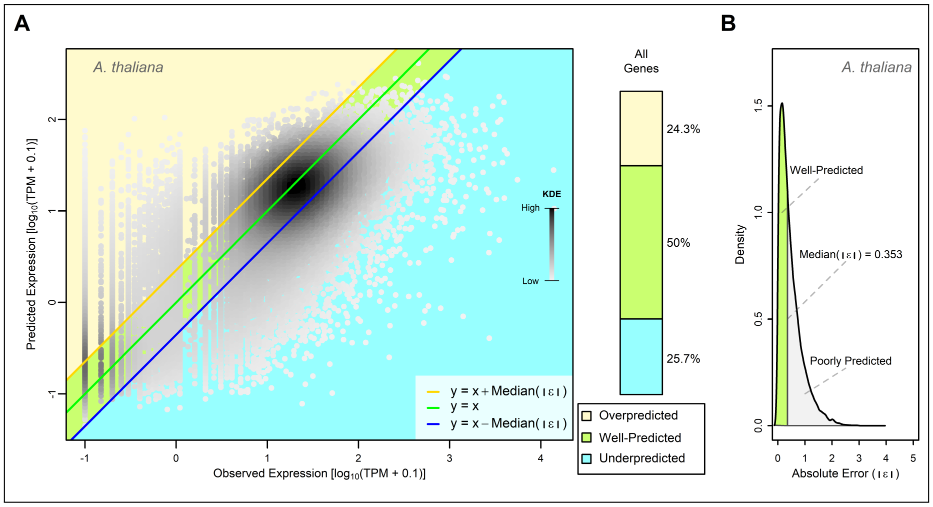


**Fig. S42.** Correlation of expression level and tissue-specificity on *A. thaliana* predictions. (A, B) Prediction errors are plotted against the median expression, showing Pearson correlations. (C, D) Prediction errors are plotted against the tissue-specificity calculated using the $\tau$ metric, showing Pearson correlations. Predictions were made for the entire data set via cross validation. The coloration of points is based on Gaussian kernel density estimations.

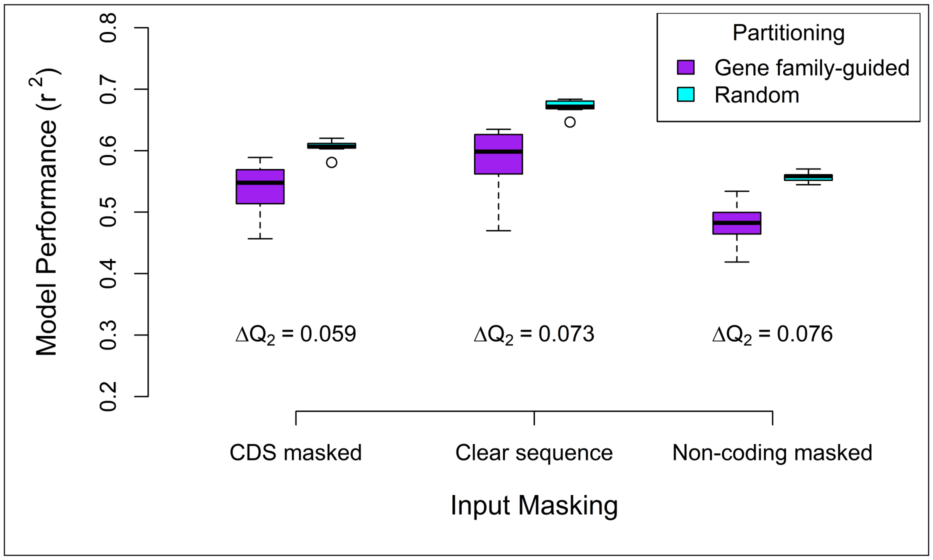


**Fig. S43.** The effect of CDS masking on overinflated performance estimations in *B. napus*. Differences in median performance between gene family-guided partitioning (purple) and random partitioning (cyan) are lowest when masking coding sequences (left), versus retaining the complete sequence (middle) or masking non-coding sequences (right).
