## Supplementary material for "A deep learning model captures position-specific effects of plant regulatory sequences and suggests genes under complex regulation": Fig S

### Supplemental figures 2: A deep learning model captures position-specific effects of plant regulatory sequences and suggests genes under complex regulation

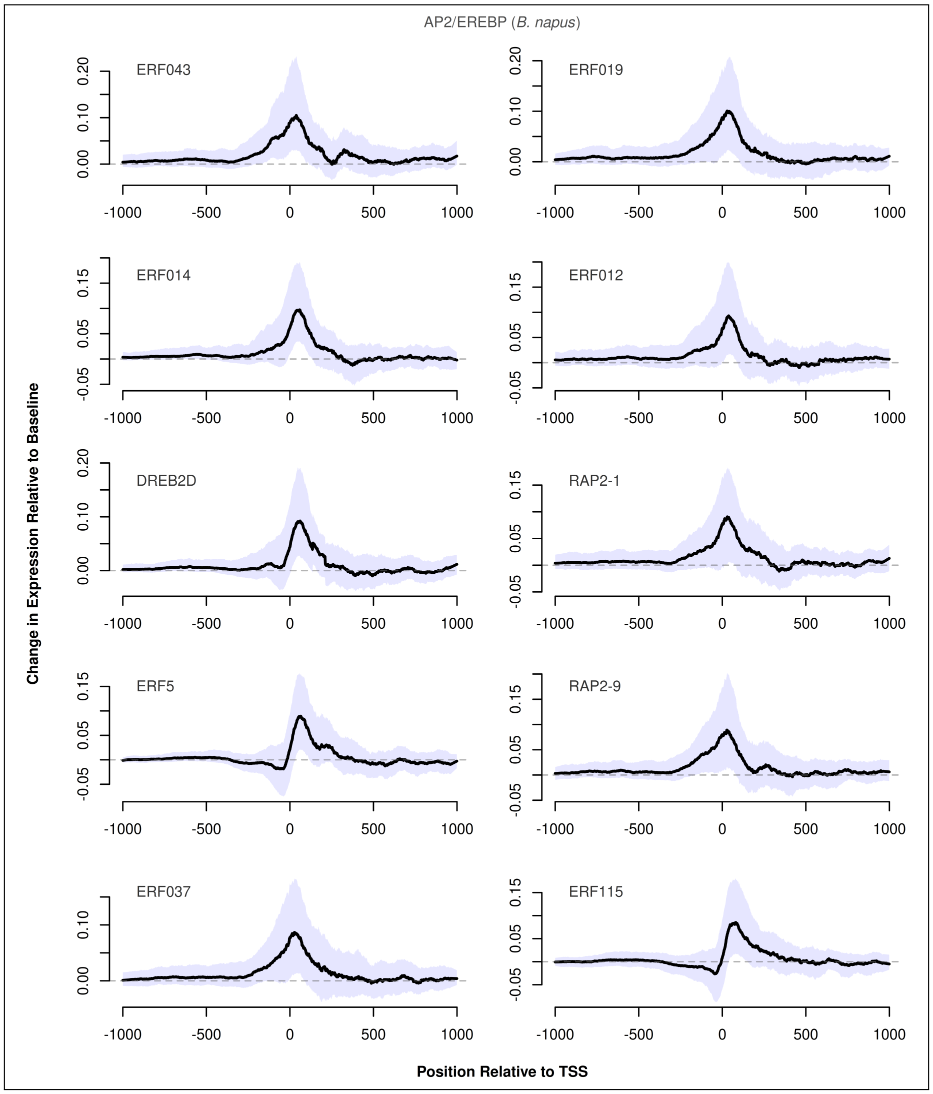


**Fig. S11.** Predicted changes in expression caused by insertion of AP2/EREBP TFs with respect to the insertion position in *B. napus*. The ten TFs for which the maximum absolute change in expression was predicted are shown, sorted by the maximum absolute change in expression from top-left to bottom-right. Black lines show the median change in expression at each position, while the shaded area shows the upper and lower quartiles. Dashed lines represent baseline expression.

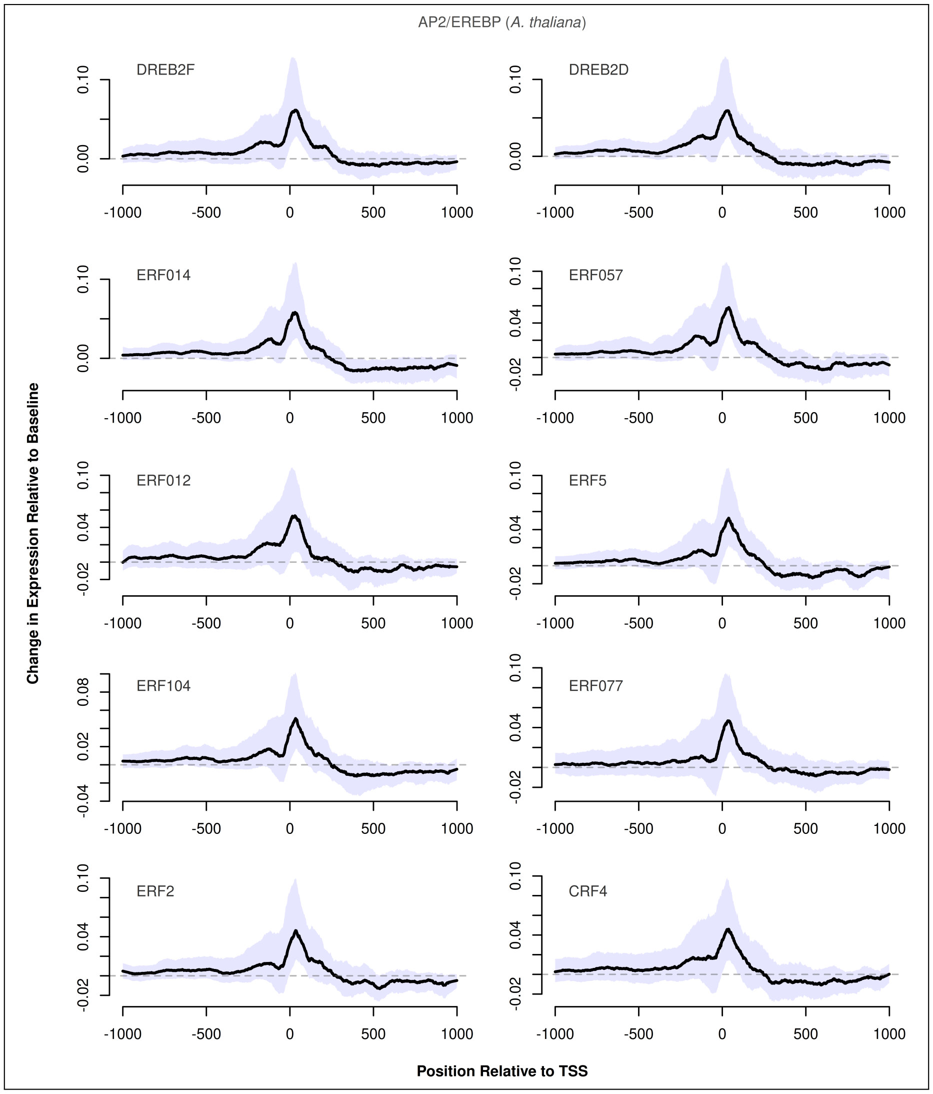


**Fig. S12.** Predicted changes in expression caused by insertion of AP2/EREBP TFs with respect to the insertion position in *A. thaliana*. The ten TFs for which the maximum absolute change in expression was predicted are shown, sorted by the maximum absolute change in expression from top-left to bottom-right. Black lines show the median change in expression at each position, while the shaded area shows the upper and lower quartiles. Dashed lines represent baseline expression.

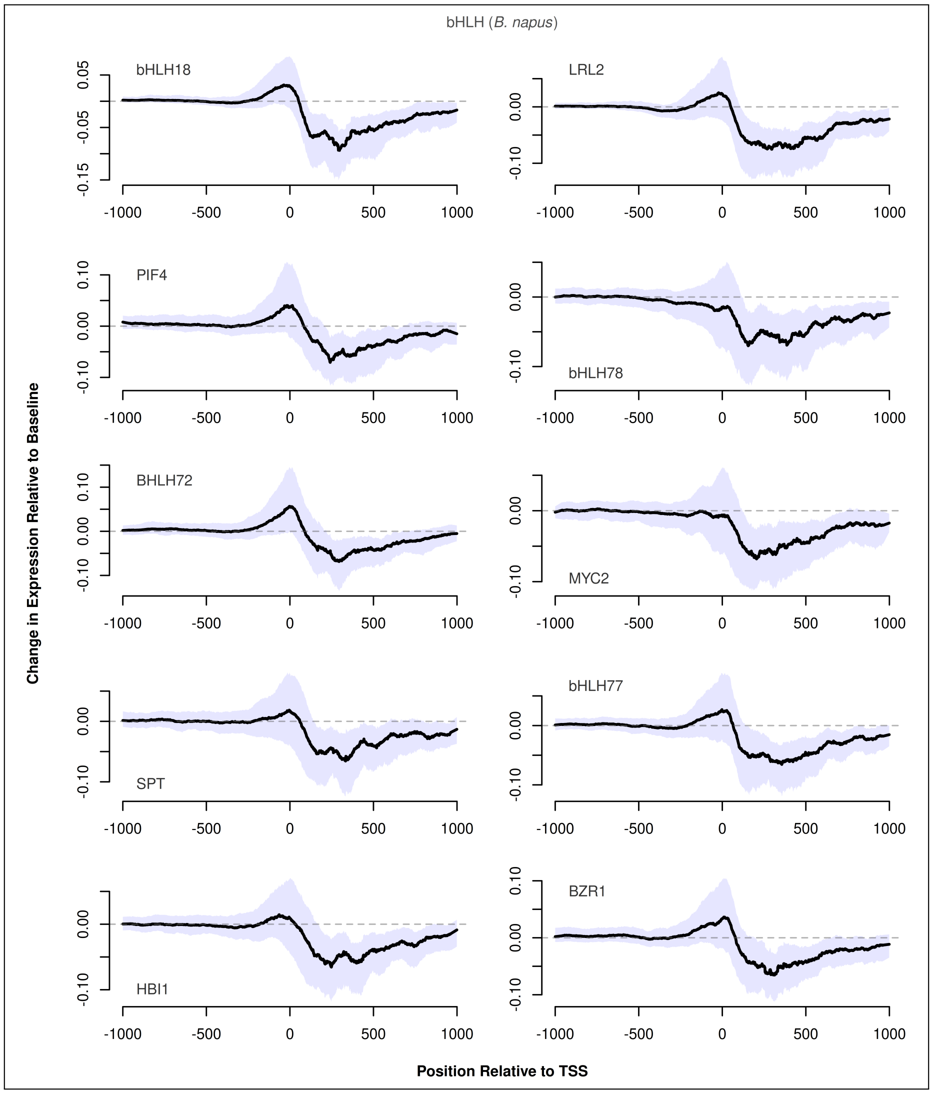


**Fig. S13.** Predicted changes in expression caused by insertion of bHLH TFs with respect to the insertion position in *B. napus*. The ten TFs for which the maximum absolute change in expression was predicted are shown, sorted by the maximum absolute change in expression from top-left to bottom-right. Black lines show the median change in expression at each position, while the shaded area shows the upper and lower quartiles. Dashed lines represent baseline expression.

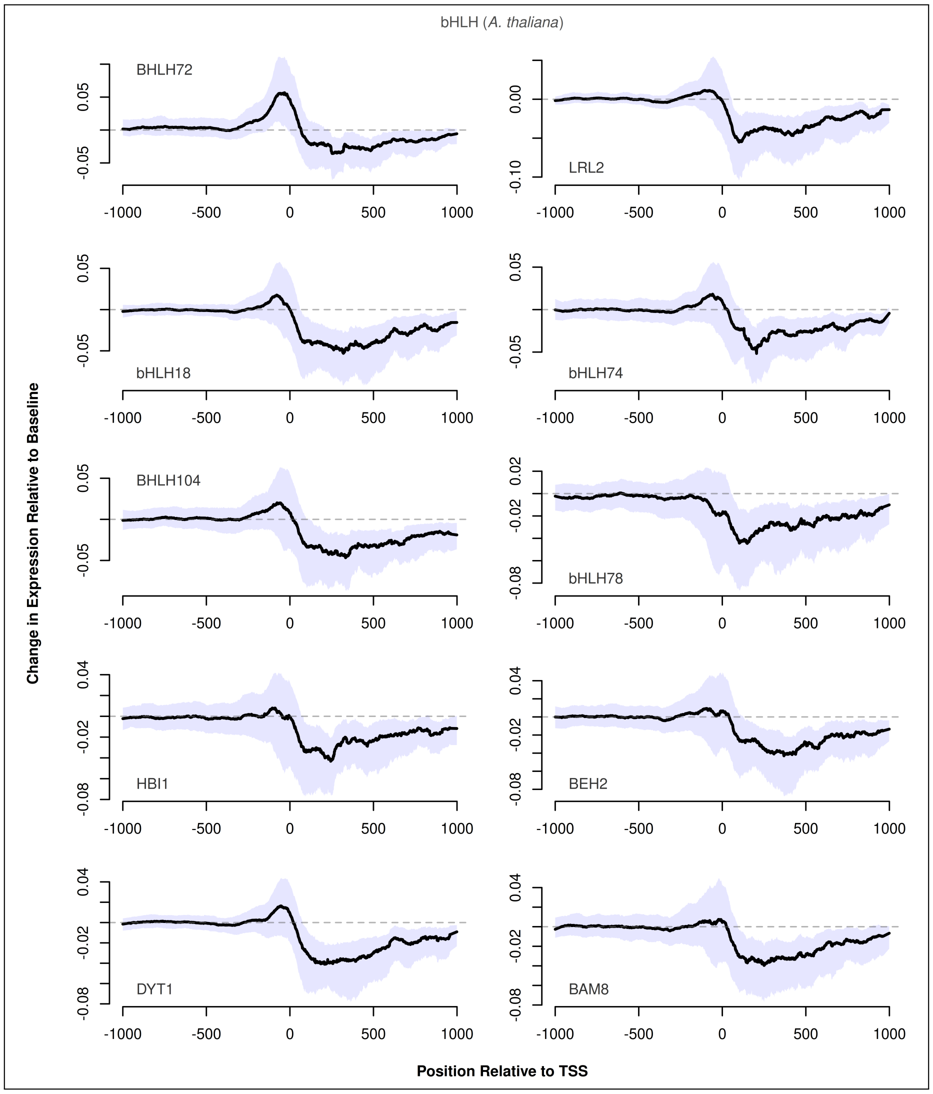


**Fig. S14.** Predicted changes in expression caused by insertion of bHLH TFs with respect to the insertion position in *A. thaliana*. The ten TFs for which the maximum absolute change in expression was predicted are shown, sorted by the maximum absolute change in expression from top-left to bottom-right. Black lines show the median change in expression at each position, while the shaded area shows the upper and lower quartiles. Dashed lines represent baseline expression.

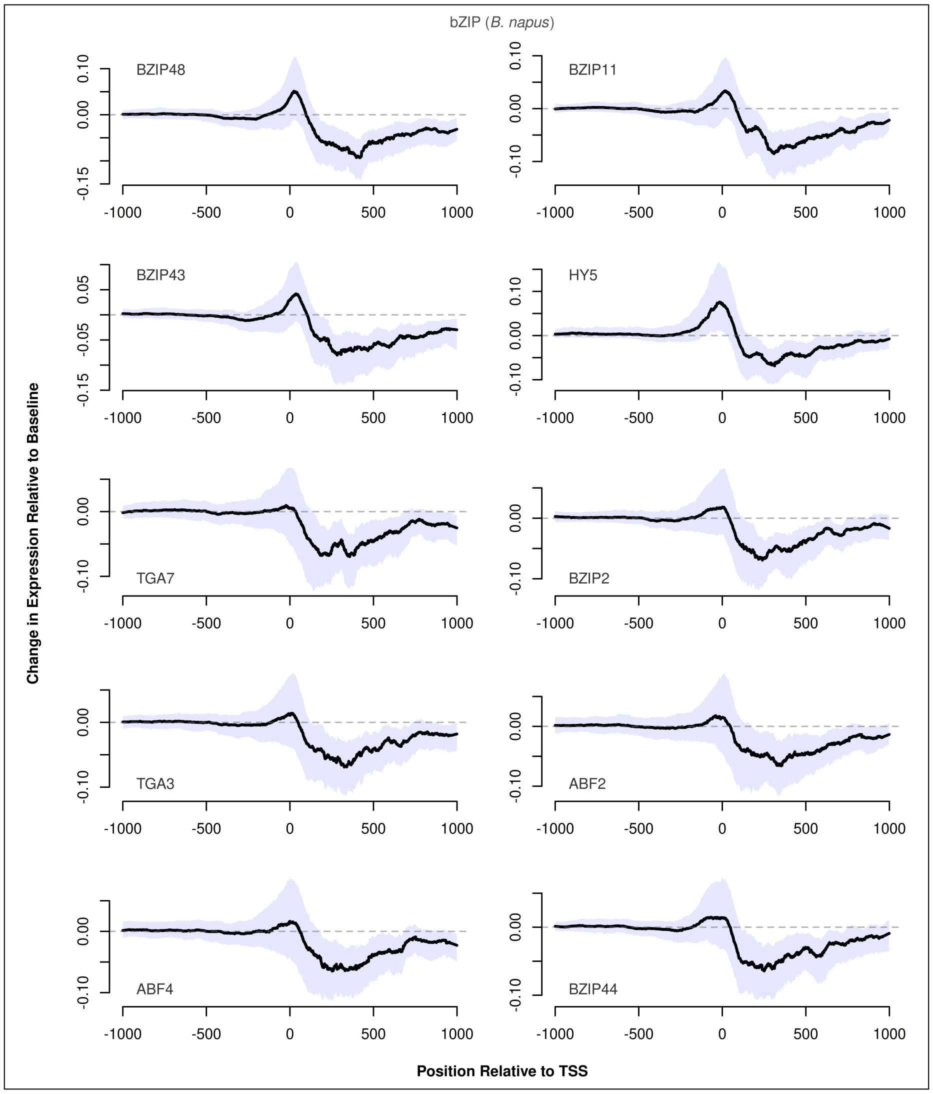


**Fig. S15.** Predicted changes in expression caused by insertion of bZIP TFs with respect to the insertion position in *B. napus*. The ten TFs for which the maximum absolute change in expression was predicted are shown, sorted by the maximum absolute change in expression from top-left to bottom-right. Black lines show the median change in expression at each position, while the shaded area shows the upper and lower quartiles. Dashed lines represent baseline expression.

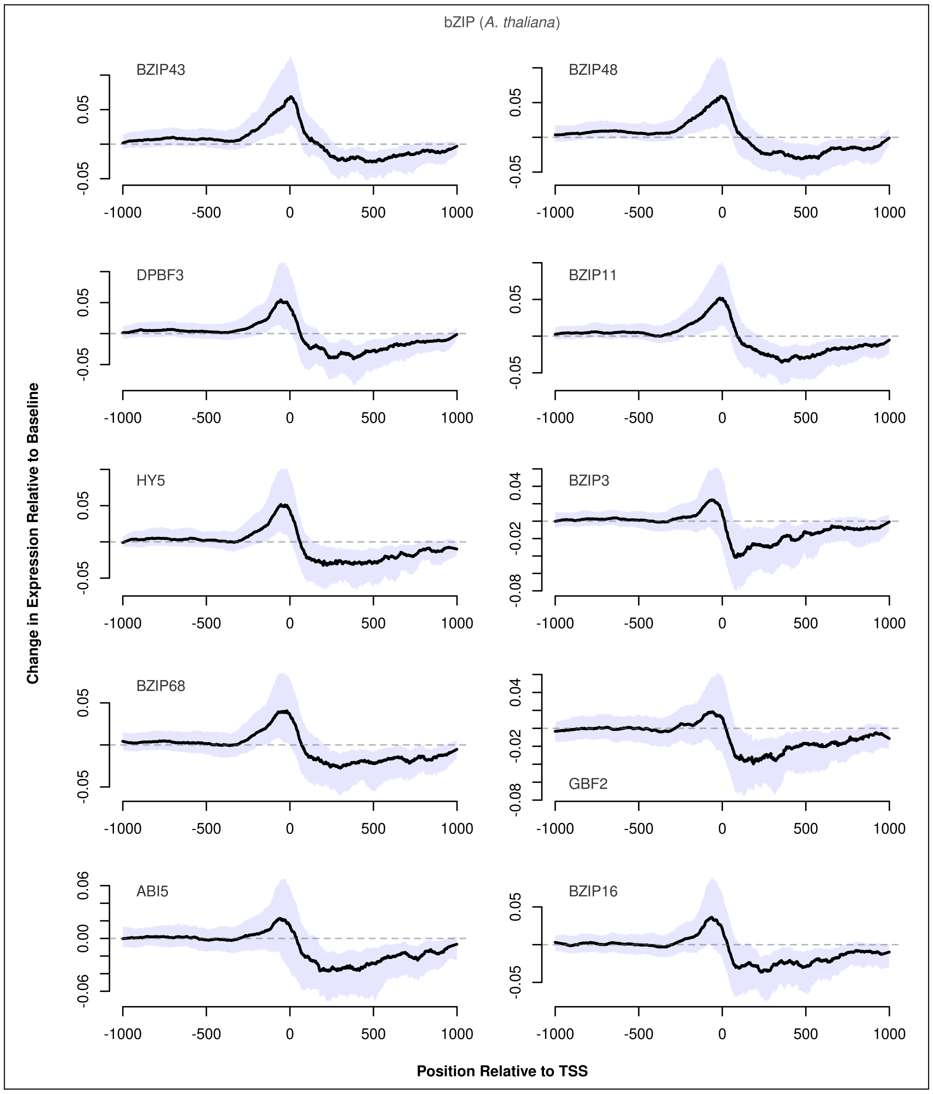


**Fig. S16.** Predicted changes in expression caused by insertion of bZIP TFs with respect to the insertion position in *A. thaliana*. The ten TFs for which the maximum absolute change in expression was predicted are shown, sorted by the maximum absolute change in expression from top-left to bottom-right. Black lines show the median change in expression at each position, while the shaded area shows the upper and lower quartiles. Dashed lines represent baseline expression.

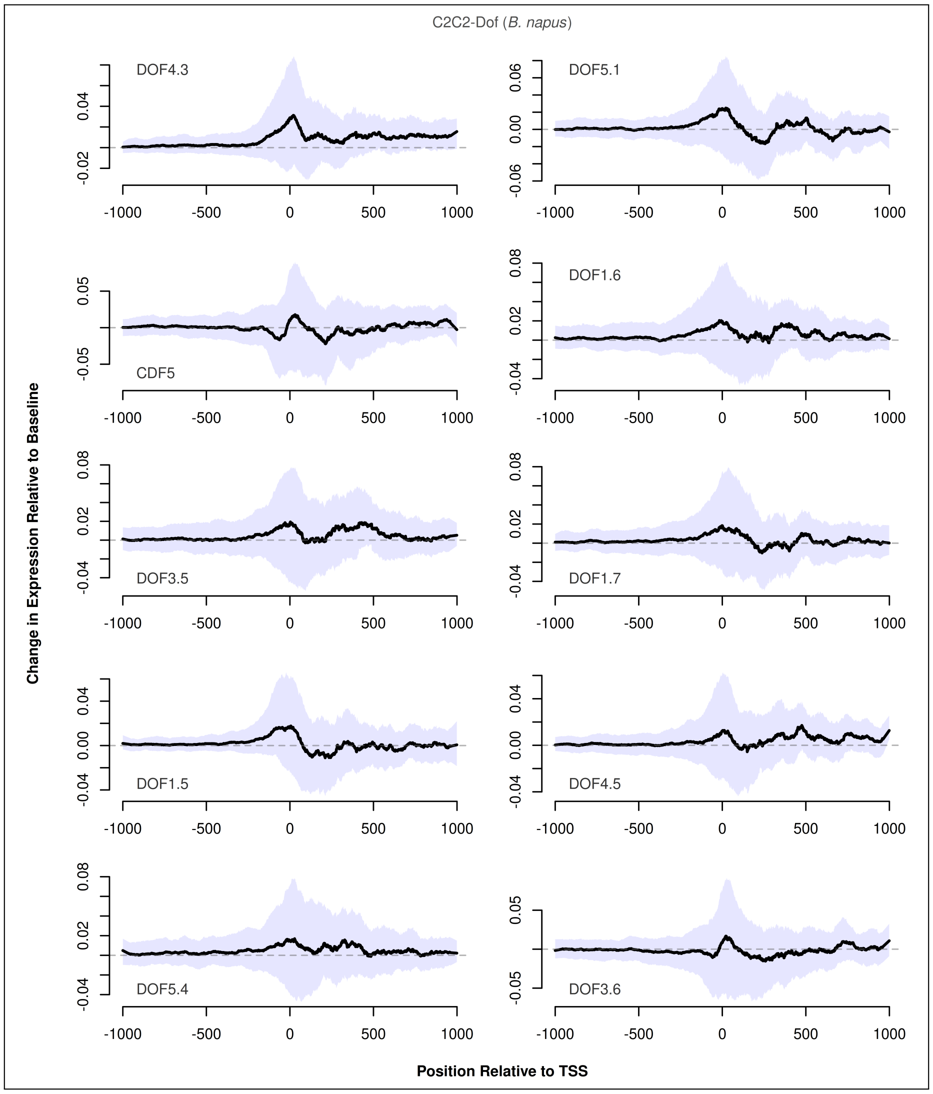


**Fig. S17.** Predicted changes in expression caused by insertion of C2C2-Dof TFs with respect to the insertion position in *B. napus*. The ten TFs for which the maximum absolute change in expression was predicted are shown, sorted by the maximum absolute change in expression from top-left to bottom-right. Black lines show the median change in expression at each position, while the shaded area shows the upper and lower quartiles. Dashed lines represent baseline expression.

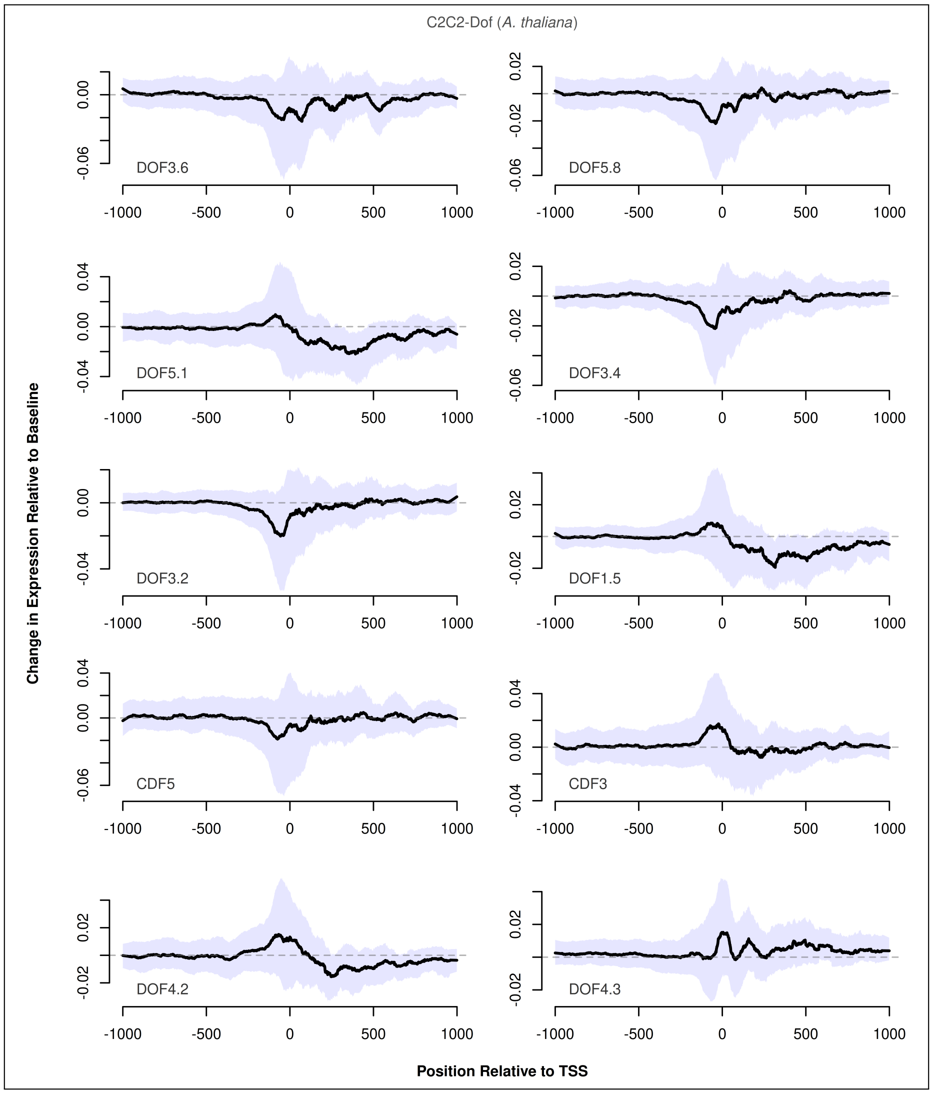


**Fig. S18.** Predicted changes in expression caused by insertion of C2C2-Dof TFs with respect to the insertion position in *A. thaliana*. The ten TFs for which the maximum absolute change in expression was predicted are shown, sorted by the maximum absolute change in expression from top-left to bottom-right. Black lines show the median change in expression at each position, while the shaded area shows the upper and lower quartiles. Dashed lines represent baseline expression.

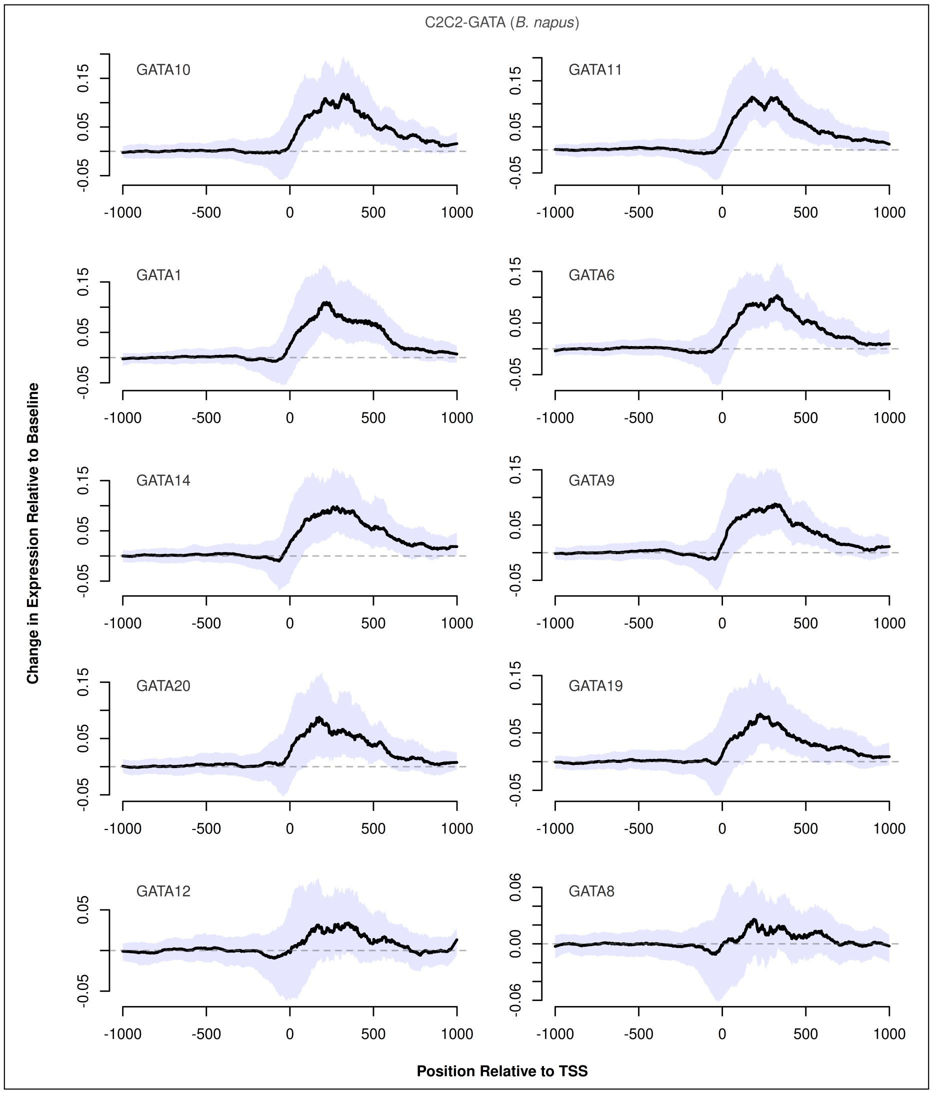


**Fig. S19.** Predicted changes in expression caused by insertion of C2C2-GATA TFs with respect to the insertion position in *B. napus*. The ten TFs for which the maximum absolute change in expression was predicted are shown, sorted by the maximum absolute change in expression from top-left to bottom-right. Black lines show the median change in expression at each position, while the shaded area shows the upper and lower quartiles. Dashed lines represent baseline expression.

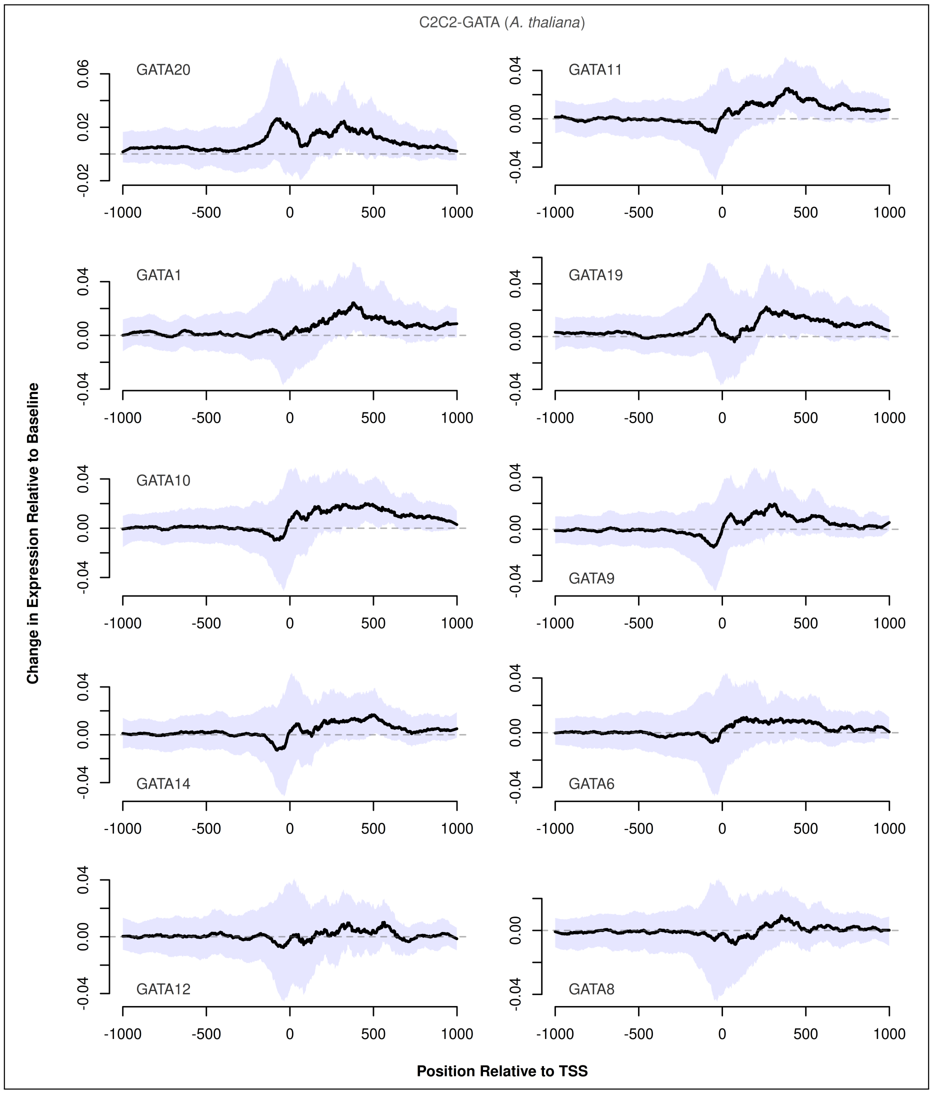


**Fig. S20.** Predicted changes in expression caused by insertion of C2C2-GATA TFs with respect to the insertion position in *A. thaliana*. The ten TFs for which the maximum absolute change in expression was predicted are shown, sorted by the maximum absolute change in expression from top-left to bottom-right. Black lines show the median change in expression at each position, while the shaded area shows the upper and lower quartiles. Dashed lines represent baseline expression.
