## Supplementary material for "A deep learning model captures position-specific effects of plant regulatory sequences and suggests genes under complex regulation": Fig S

### Supplemental figures 3: A deep learning model captures position-specific effects of plant regulatory sequences and suggests genes under complex regulation

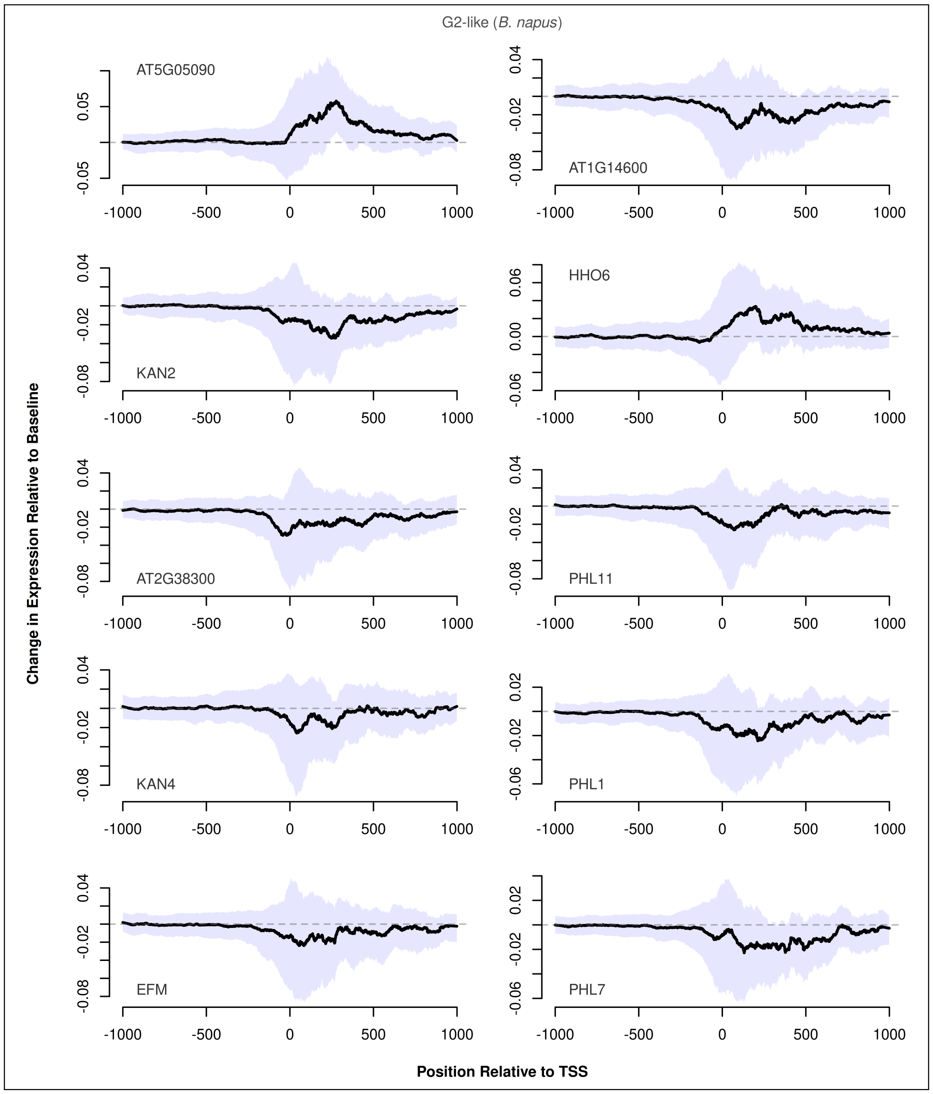


**Fig. S21.** Predicted changes in expression caused by insertion of G2-like TFs with respect to the insertion position in *B. napus*. The ten TFs for which the maximum absolute change in expression was predicted are shown, sorted by the maximum absolute change in expression from top-left to bottom-right. Black lines show the median change in expression at each position, while the shaded area shows the upper and lower quartiles. Dashed lines represent baseline expression.

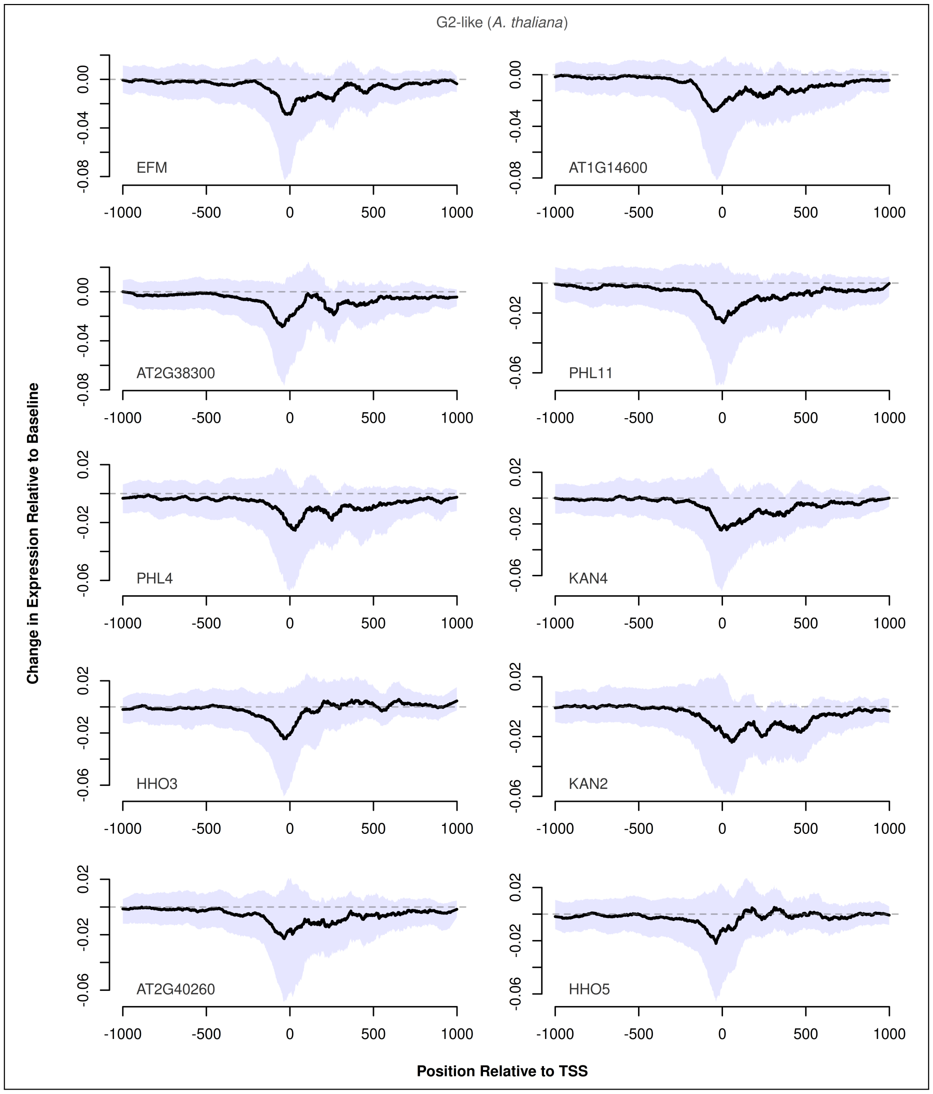


**Fig. S22.** Predicted changes in expression caused by insertion of G2-like TFs with respect to the insertion position in *A. thaliana*. The ten TFs for which the maximum absolute change in expression was predicted are shown, sorted by the maximum absolute change in expression from top-left to bottom-right. Black lines show the median change in expression at each position, while the shaded area shows the upper and lower quartiles. Dashed lines represent baseline expression.

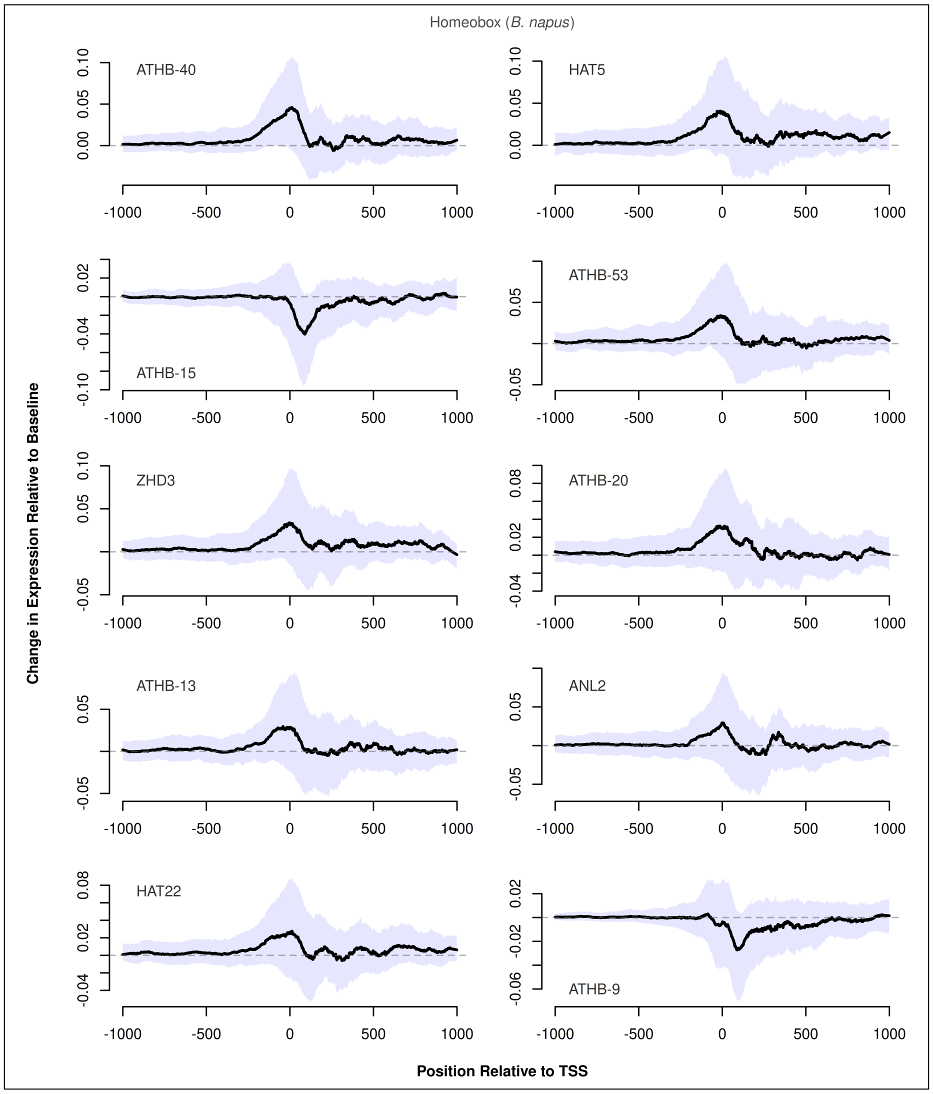


**Fig. S23.** Predicted changes in expression caused by insertion of Homeobox TFs with respect to the insertion position in *B. napus*. The ten TFs for which the maximum absolute change in expression was predicted are shown, sorted by the maximum absolute change in expression from top-left to bottom-right. Black lines show the median change in expression at each position, while the shaded area shows the upper and lower quartiles. Dashed lines represent baseline expression.

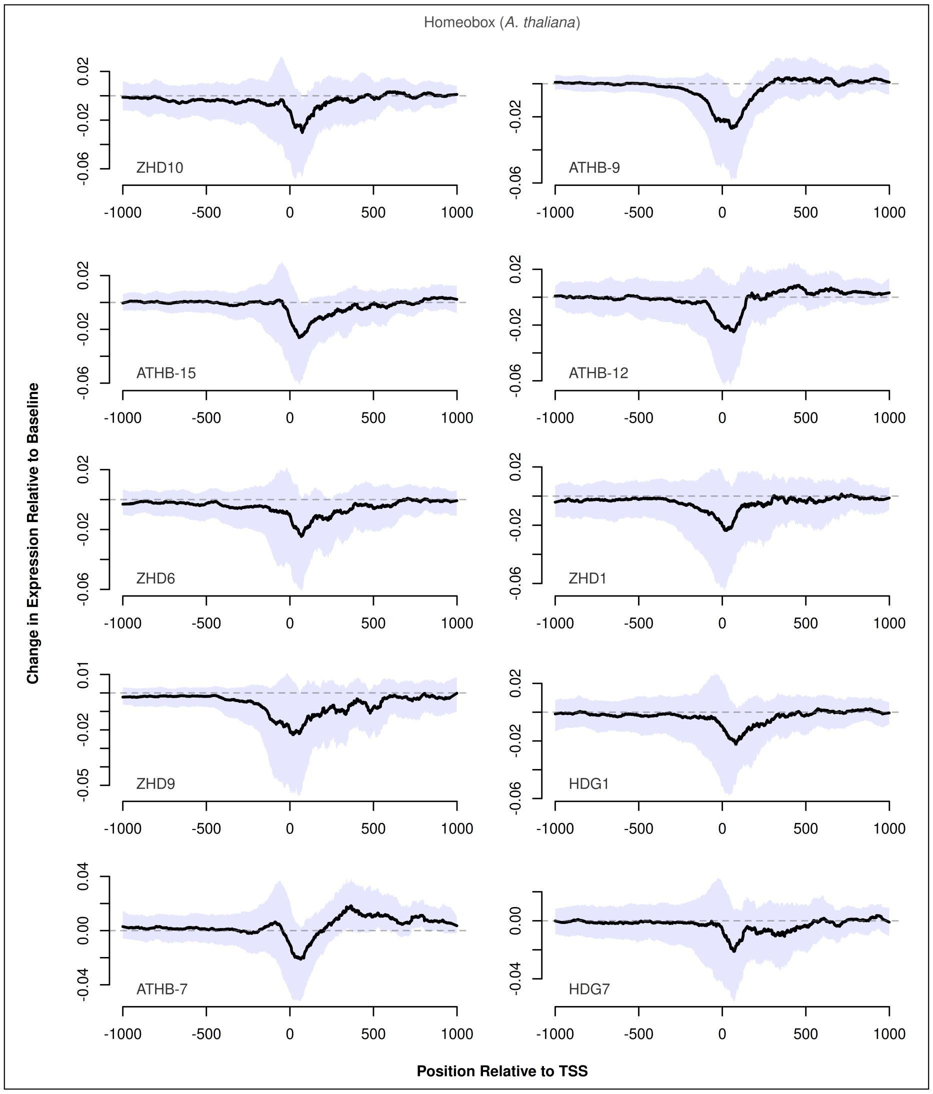


**Fig. S24.** Predicted changes in expression caused by insertion of Homeobox TFs with respect to the insertion position in *A. thaliana*. The ten TFs for which the maximum absolute change in expression was predicted are shown, sorted by the maximum absolute change in expression from top-left to bottom-right. Black lines show the median change in expression at each position, while the shaded area shows the upper and lower quartiles. Dashed lines represent baseline expression.

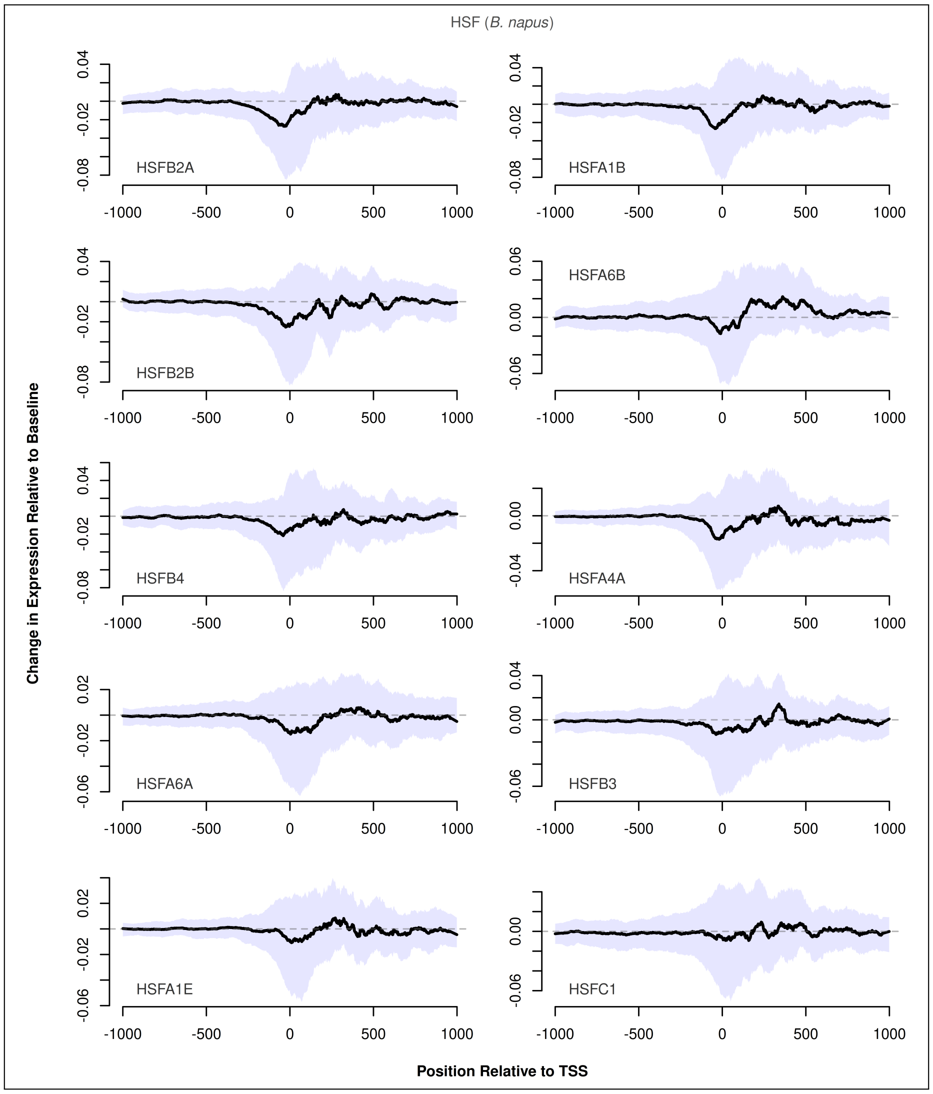


**Fig. S25.** Predicted changes in expression caused by insertion of HSF TFs with respect to the insertion position in *B. napus*. The ten TFs for which the maximum absolute change in expression was predicted are shown, sorted by the maximum absolute change in expression from top-left to bottom-right. Black lines show the median change in expression at each position, while the shaded area shows the upper and lower quartiles. Dashed lines represent baseline expression.

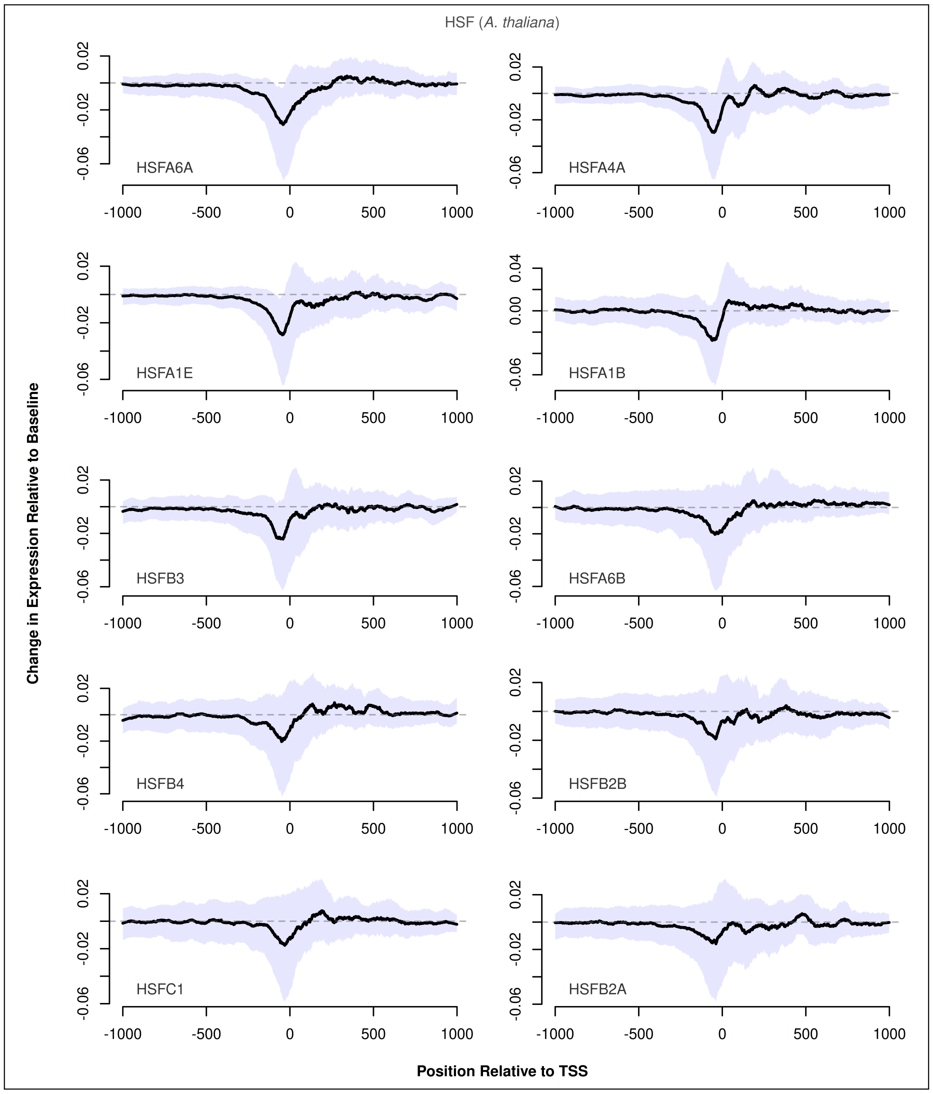


**Fig. S26.** Predicted changes in expression caused by insertion of HSF TFs with respect to the insertion position in *A. thaliana*. The ten TFs for which the maximum absolute change in expression was predicted are shown, sorted by the maximum absolute change in expression from top-left to bottom-right. Black lines show the median change in expression at each position, while the shaded area shows the upper and lower quartiles. Dashed lines represent baseline expression.

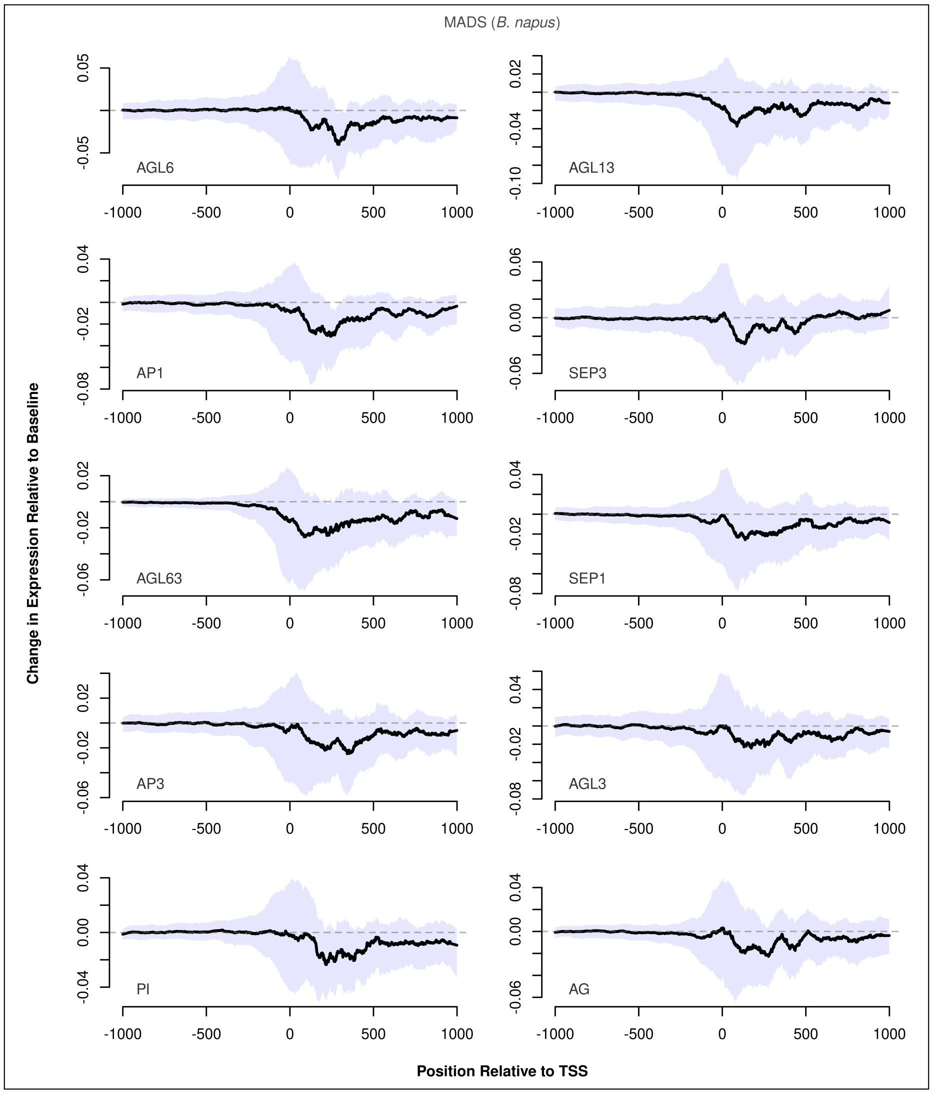


**Fig. S27.** Predicted changes in expression caused by insertion of MADS TFs with respect to the insertion position in *B. napus*. The ten TFs for which the maximum absolute change in expression was predicted are shown, sorted by the maximum absolute change in expression from top-left to bottom-right. Black lines show the median change in expression at each position, while the shaded area shows the upper and lower quartiles. Dashed lines represent baseline expression.

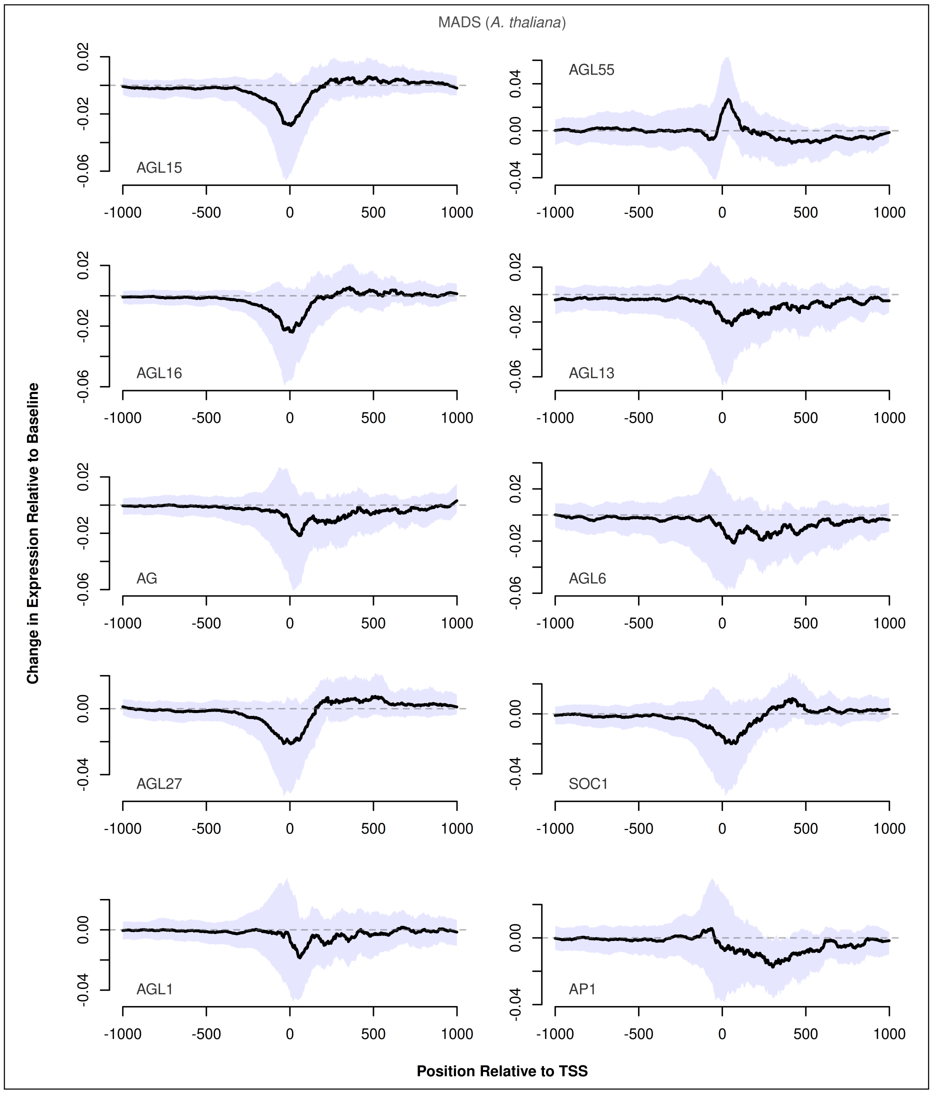


**Fig. S28.** Predicted changes in expression caused by insertion of MADS TFs with respect to the insertion position in *A. thaliana*. The ten TFs for which the maximum absolute change in expression was predicted are shown, sorted by the maximum absolute change in expression from top-left to bottom-right. Black lines show the median change in expression at each position, while the shaded area shows the upper and lower quartiles. Dashed lines represent baseline expression.

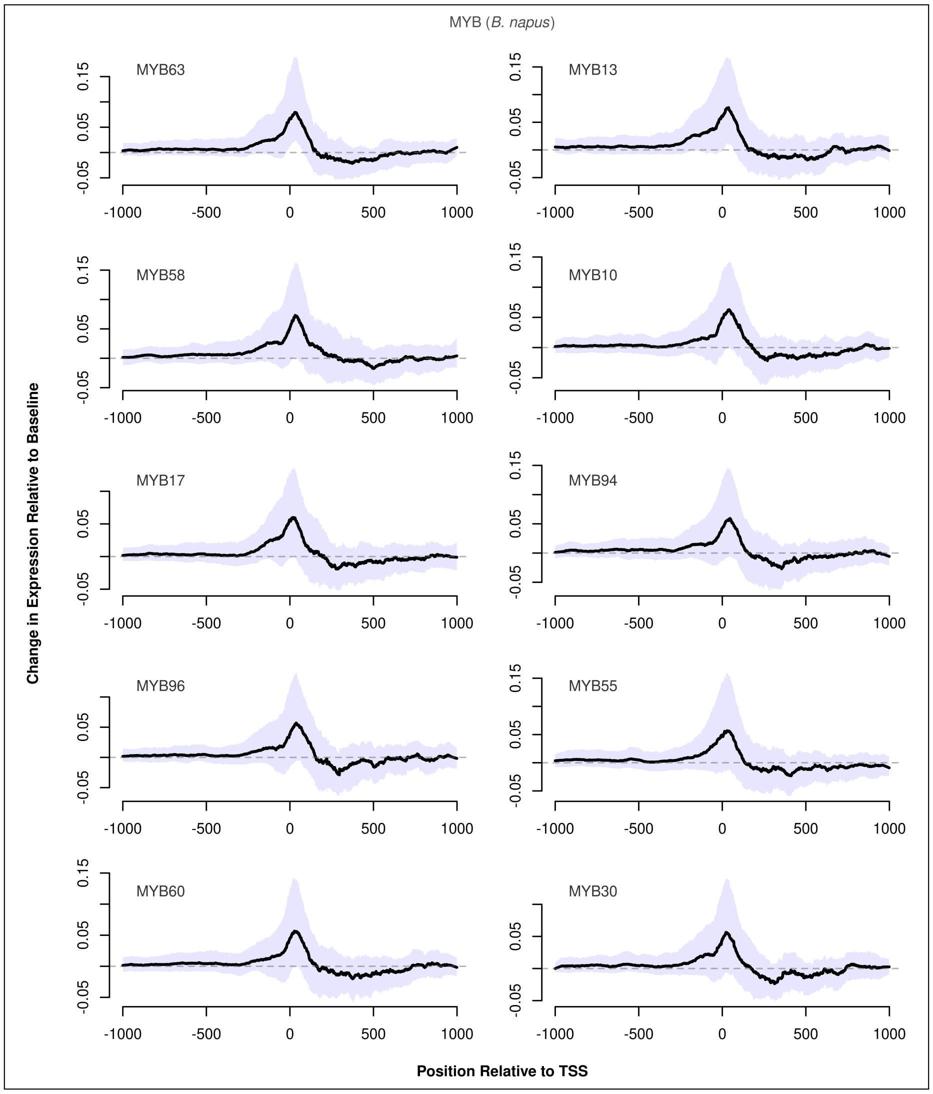


**Fig. S29.** Predicted changes in expression caused by insertion of MYB TFs with respect to the insertion position in *B. napus*. The ten TFs for which the maximum absolute change in expression was predicted are shown, sorted by the maximum absolute change in expression from top-left to bottom-right. Black lines show the median change in expression at each position, while the shaded area shows the upper and lower quartiles. Dashed lines represent baseline expression.

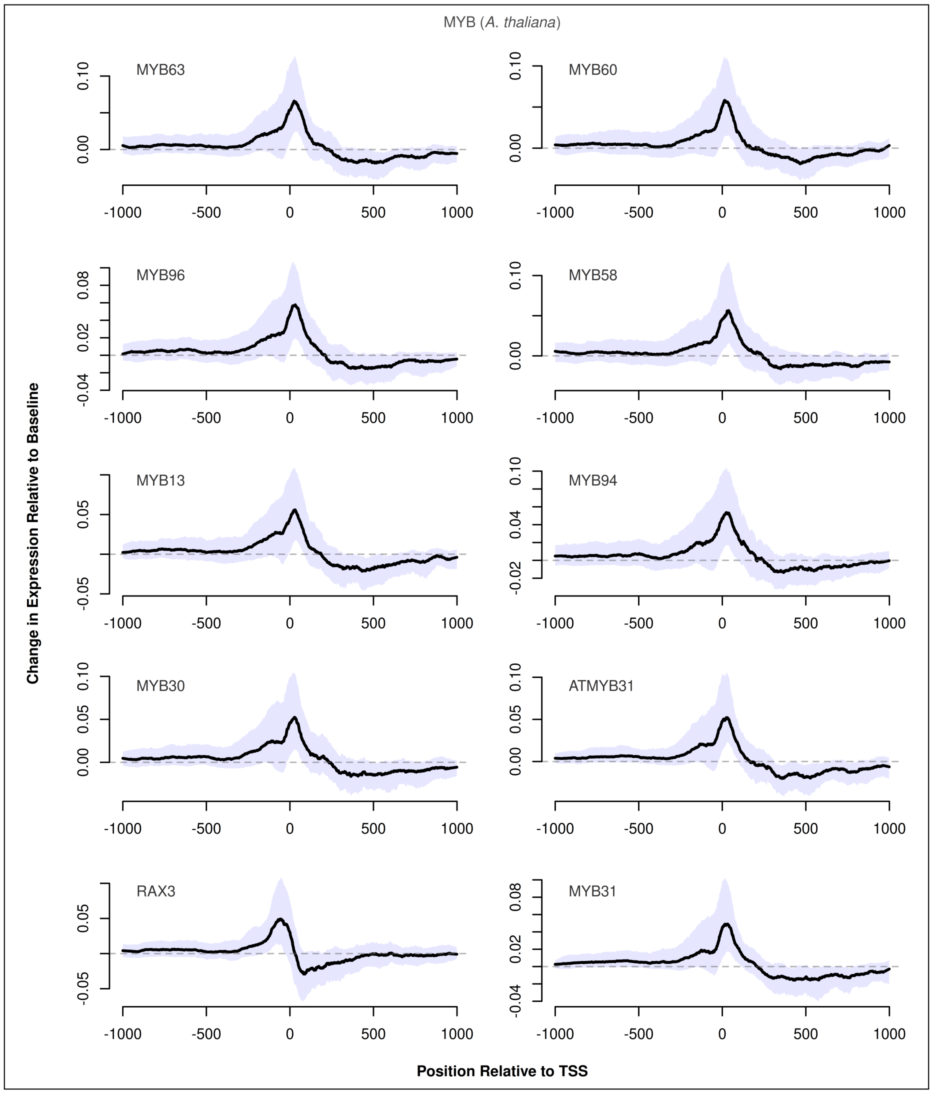


**Fig. S30.** Predicted changes in expression caused by insertion of MYB TFs with respect to the insertion position in *A. thaliana*. The ten TFs for which the maximum absolute change in expression was predicted are shown, sorted by the maximum absolute change in expression from top-left to bottom-right. Black lines show the median change in expression at each position, while the shaded area shows the upper and lower quartiles. Dashed lines represent baseline expression.
