## Supplementary material for "A deep learning model captures position-specific effects of plant regulatory sequences and suggests genes under complex regulation": Fig S

### Supplemental figures 4: A deep learning model captures position-specific effects of plant regulatory sequences and suggests genes under complex regulation

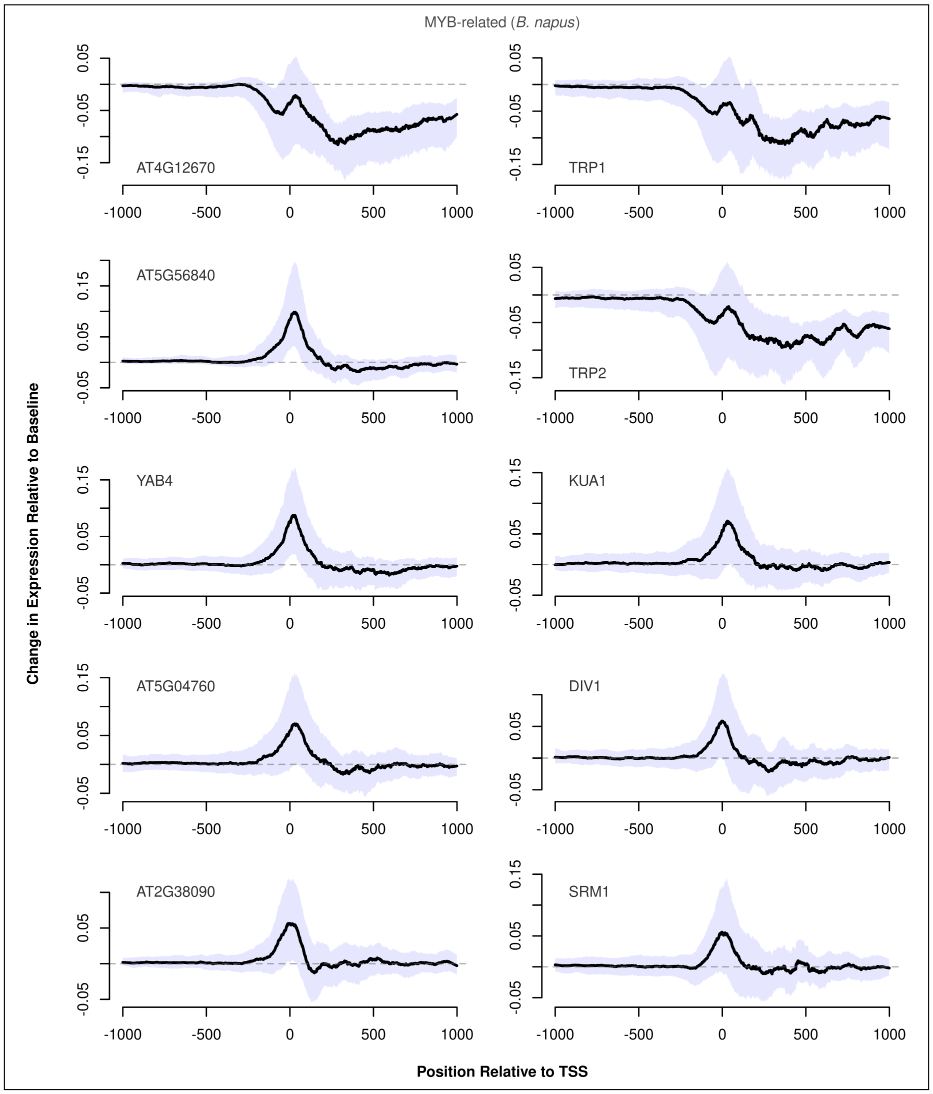


**Fig. S31.** Predicted changes in expression caused by insertion of MYB-related TFs with respect to the insertion position in *B. napus*. The ten TFs for which the maximum absolute change in expression was predicted are shown, sorted by the maximum absolute change in expression from top-left to bottom-right. Black lines show the median change in expression at each position, while the shaded area shows the upper and lower quartiles. Dashed lines represent baseline expression.

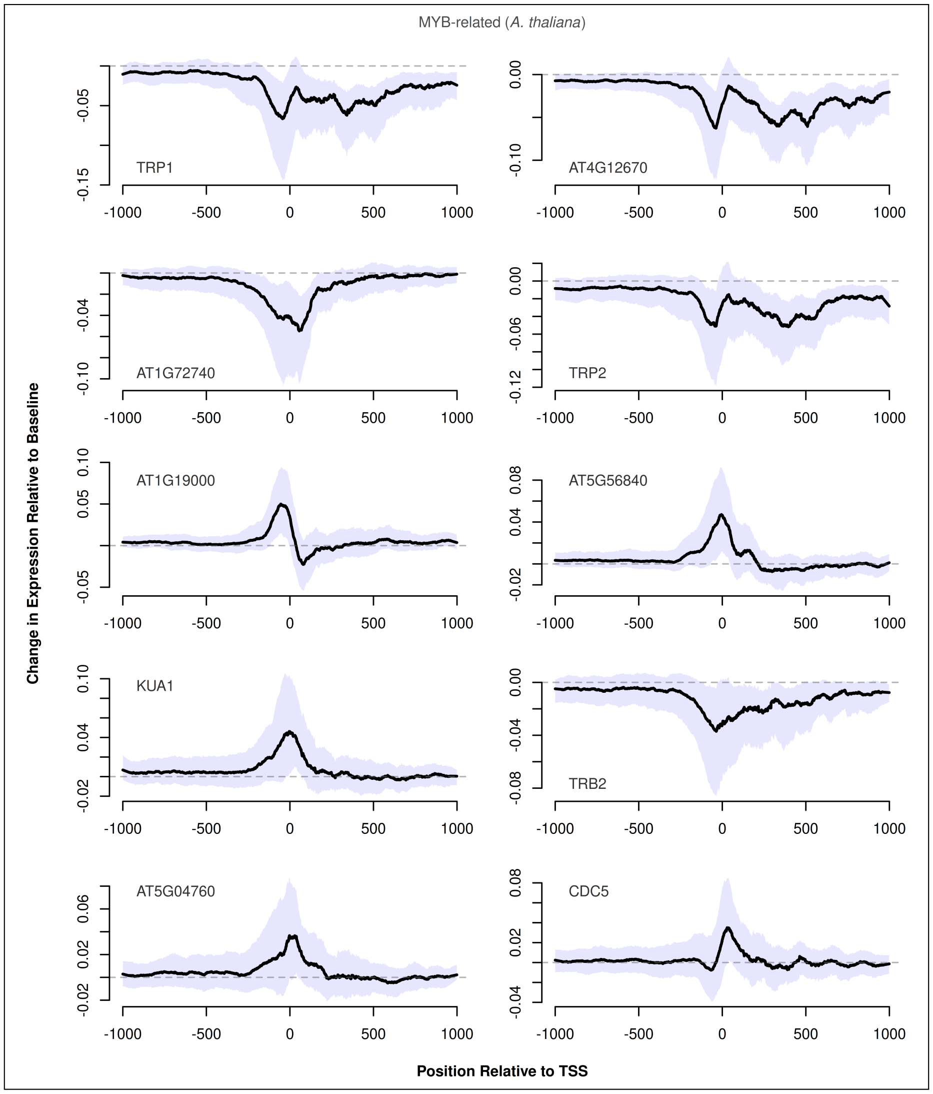


**Fig. S32.** Predicted changes in expression caused by insertion of MYB-related TFs with respect to the insertion position in *A. thaliana*. The ten TFs for which the maximum absolute change in expression was predicted are shown, sorted by the maximum absolute change in expression from top-left to bottom-right. Black lines show the median change in expression at each position, while the shaded area shows the upper and lower quartiles. Dashed lines represent baseline expression.

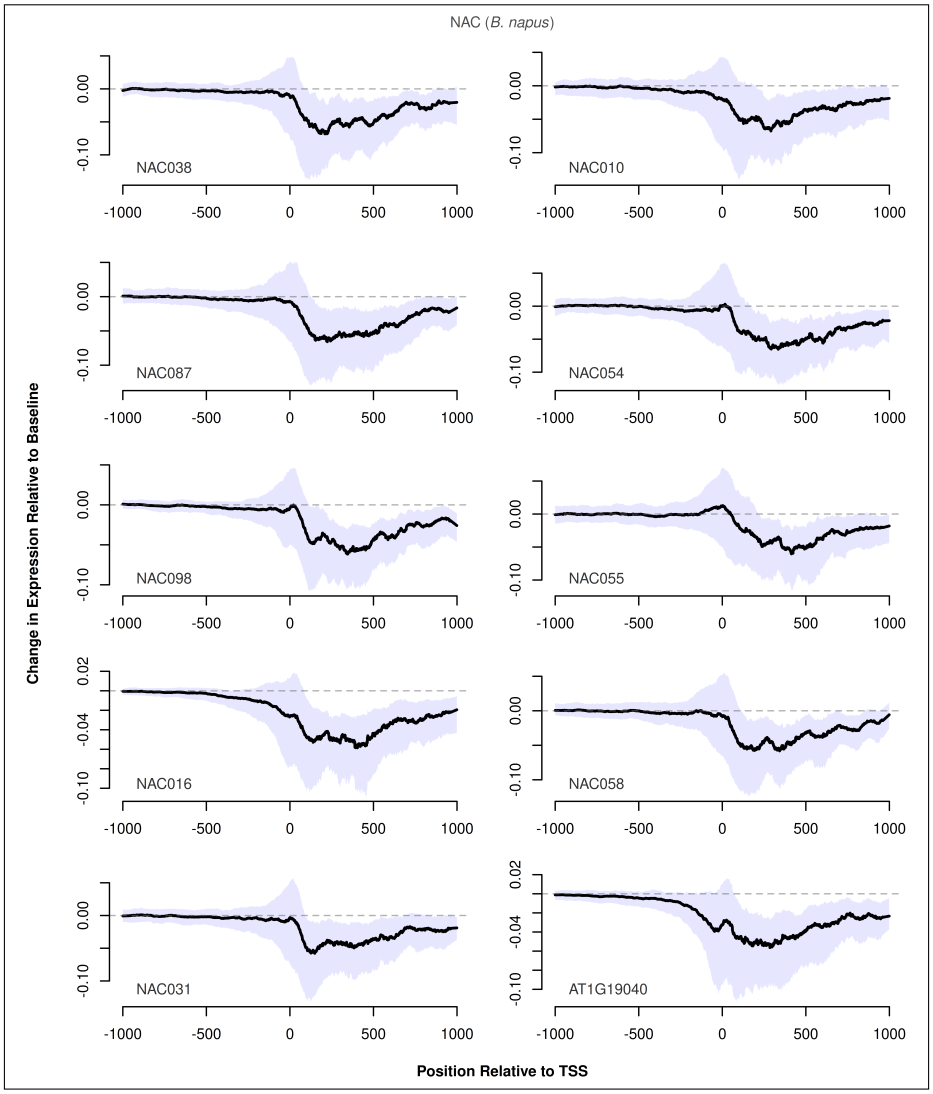


**Fig. S33.** Predicted changes in expression caused by insertion of NAC TFs with respect to the insertion position in *B. napus*. The ten TFs for which the maximum absolute change in expression was predicted are shown, sorted by the maximum absolute change in expression from top-left to bottom-right. Black lines show the median change in expression at each position, while the shaded area shows the upper and lower quartiles. Dashed lines represent baseline expression.

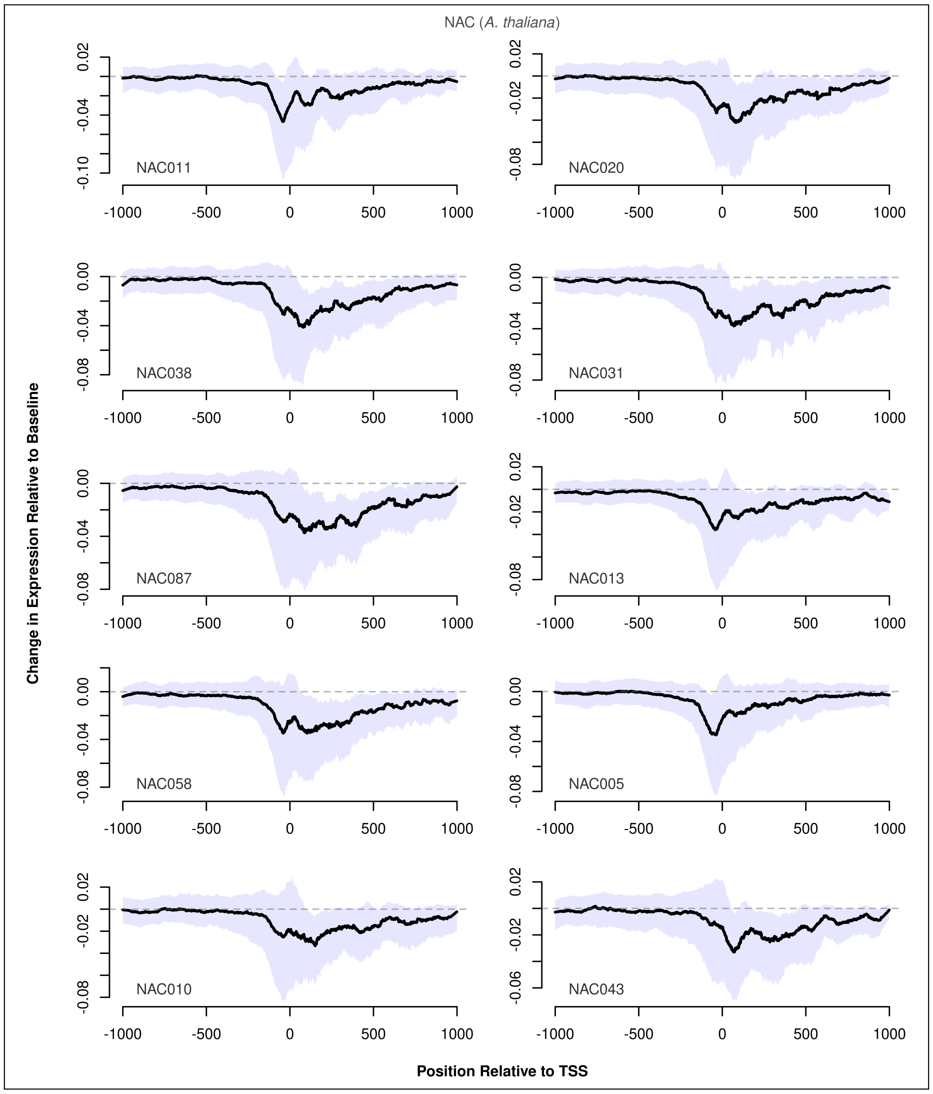


**Fig. S34.** Predicted changes in expression caused by insertion of NAC TFs with respect to the insertion position in *A. thaliana*. The ten TFs for which the maximum absolute change in expression was predicted are shown, sorted by the maximum absolute change in expression from top-left to bottom-right. Black lines show the median change in expression at each position, while the shaded area shows the upper and lower quartiles. Dashed lines represent baseline expression.

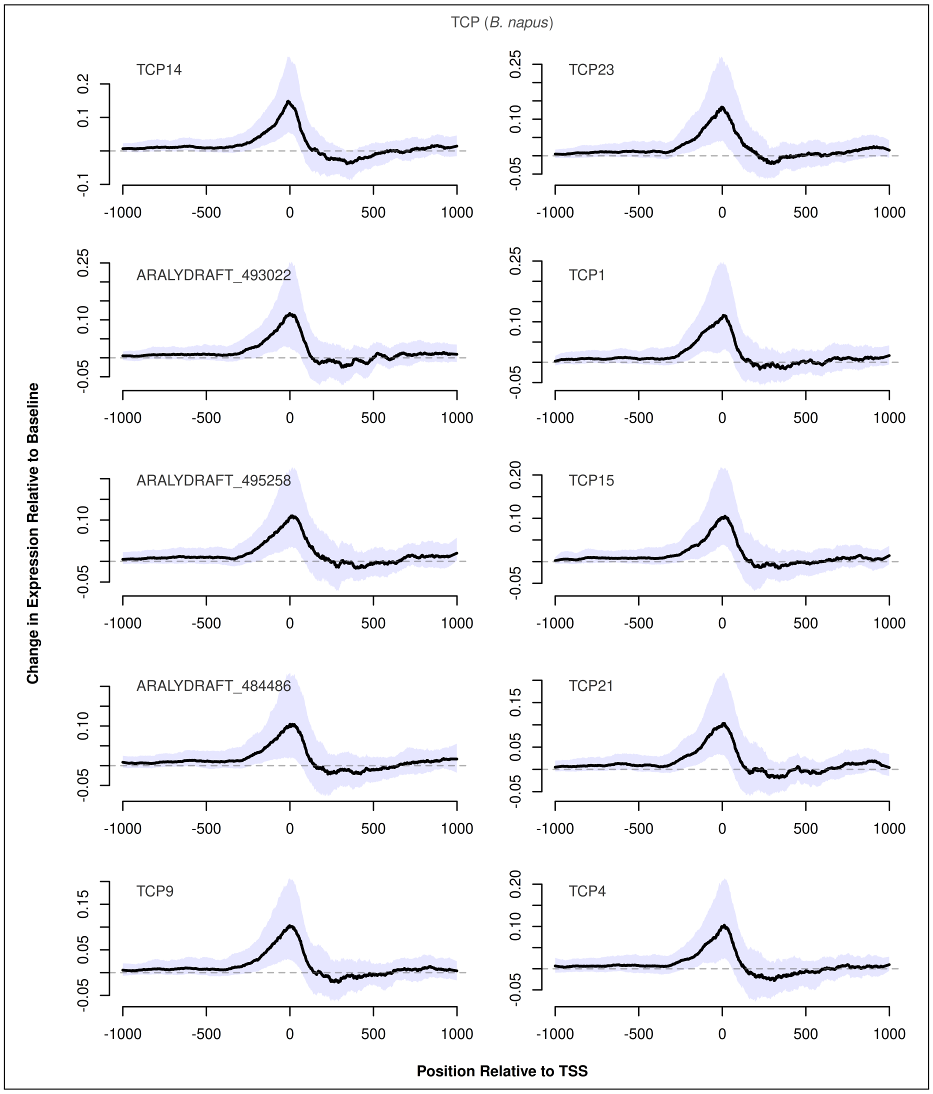


**Fig. S35.** Predicted changes in expression caused by insertion of TCP TFs with respect to the insertion position in *B. napus*. The ten TFs for which the maximum absolute change in expression was predicted are shown, sorted by the maximum absolute change in expression from top-left to bottom-right. Black lines show the median change in expression at each position, while the shaded area shows the upper and lower quartiles. Dashed lines represent baseline expression.

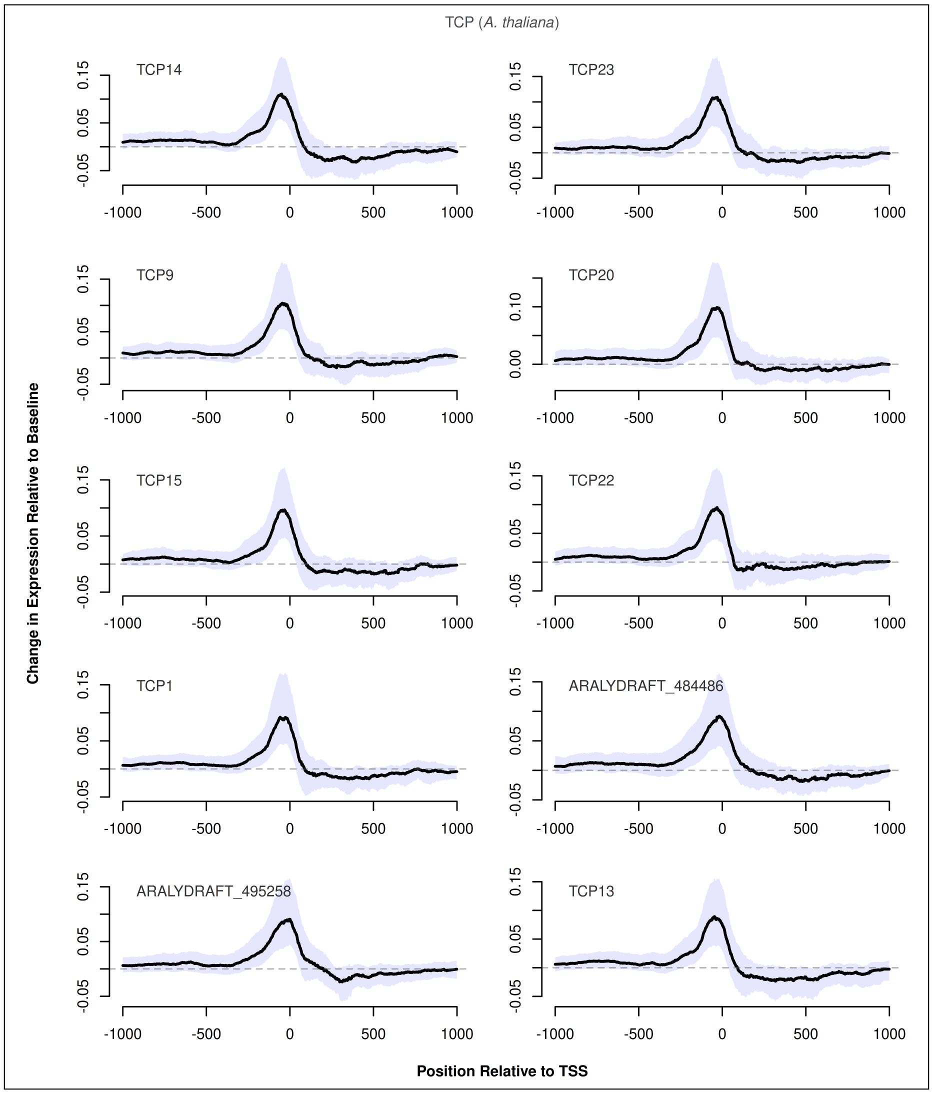


**Fig. S36.** Predicted changes in expression caused by insertion of TCP TFs with respect to the insertion position in *A. thaliana*. The ten TFs for which the maximum absolute change in expression was predicted are shown, sorted by the maximum absolute change in expression from top-left to bottom-right. Black lines show the median change in expression at each position, while the shaded area shows the upper and lower quartiles. Dashed lines represent baseline expression.

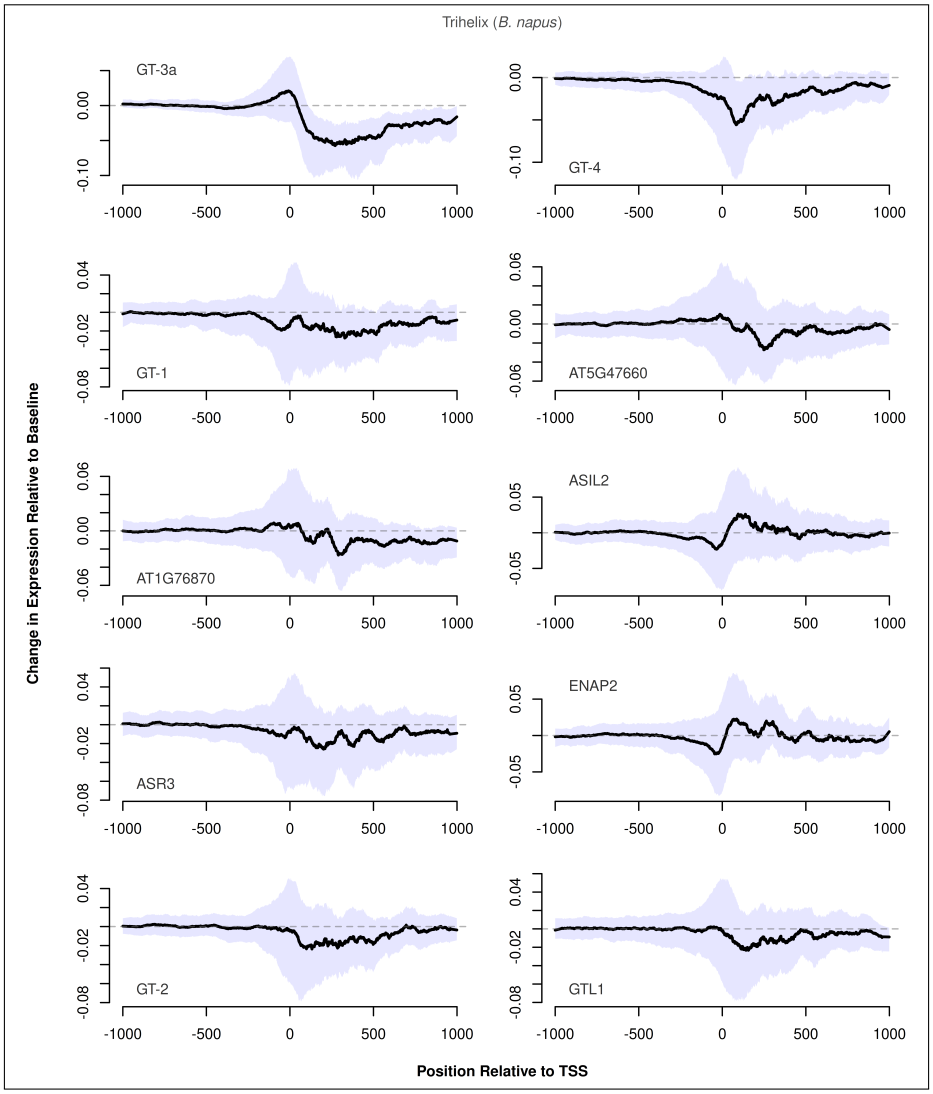


**Fig. S37.** Predicted changes in expression caused by insertion of Trihelix TFs with respect to the insertion position in *B. napus*. The ten TFs for which the maximum absolute change in expression was predicted are shown, sorted by the maximum absolute change in expression from top-left to bottom-right. Black lines show the median change in expression at each position, while the shaded area shows the upper and lower quartiles. Dashed lines represent baseline expression.

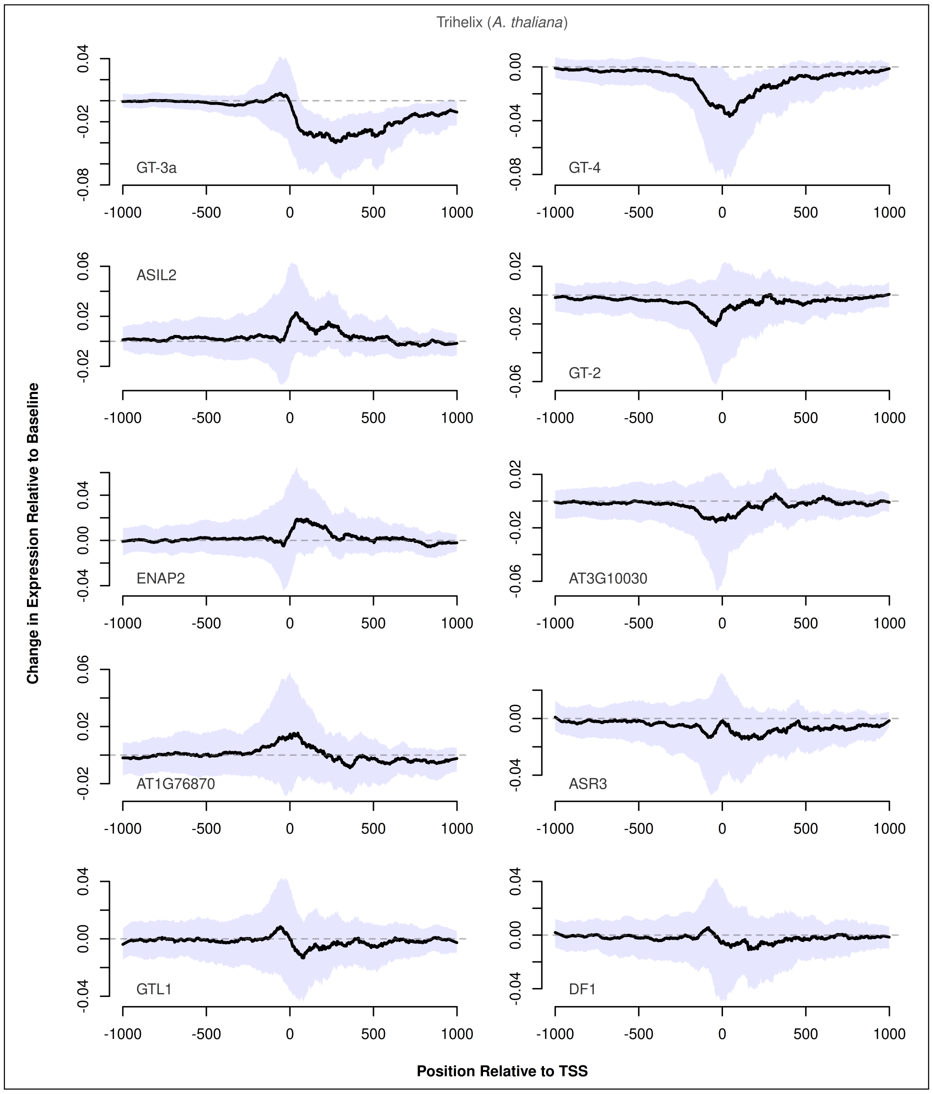


**Fig. S38.** Predicted changes in expression caused by insertion of Trihelix TFs with respect to the insertion position in *A. thaliana*. The ten TFs for which the maximum absolute change in expression was predicted are shown, sorted by the maximum absolute change in expression from top-left to bottom-right. Black lines show the median change in expression at each position, while the shaded area shows the upper and lower quartiles. Dashed lines represent baseline expression.

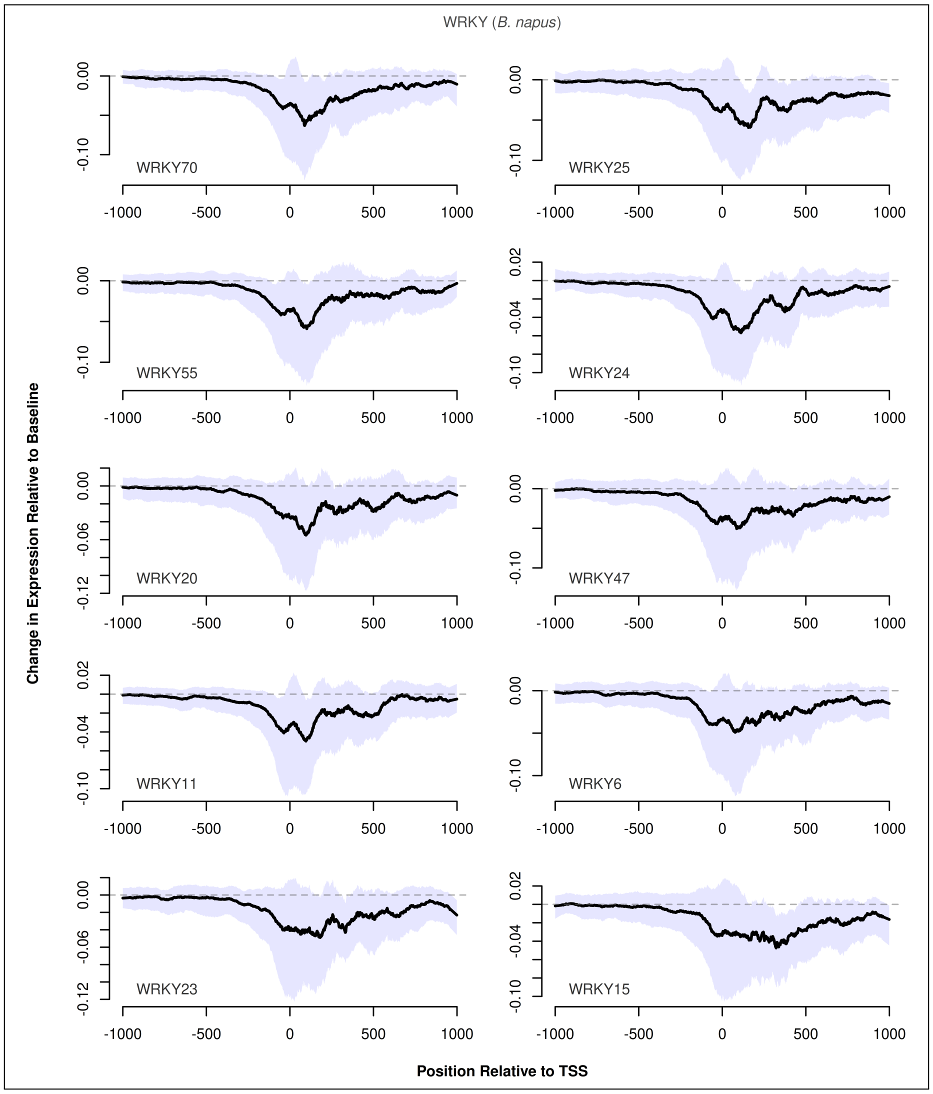


**Fig. S39.** Predicted changes in expression caused by insertion of WRKY TFs with respect to the insertion position in *B. napus*. The ten TFs for which the maximum absolute change in expression was predicted are shown, sorted by the maximum absolute change in expression from top-left to bottom-right. Black lines show the median change in expression at each position, while the shaded area shows the upper and lower quartiles. Dashed lines represent baseline expression.

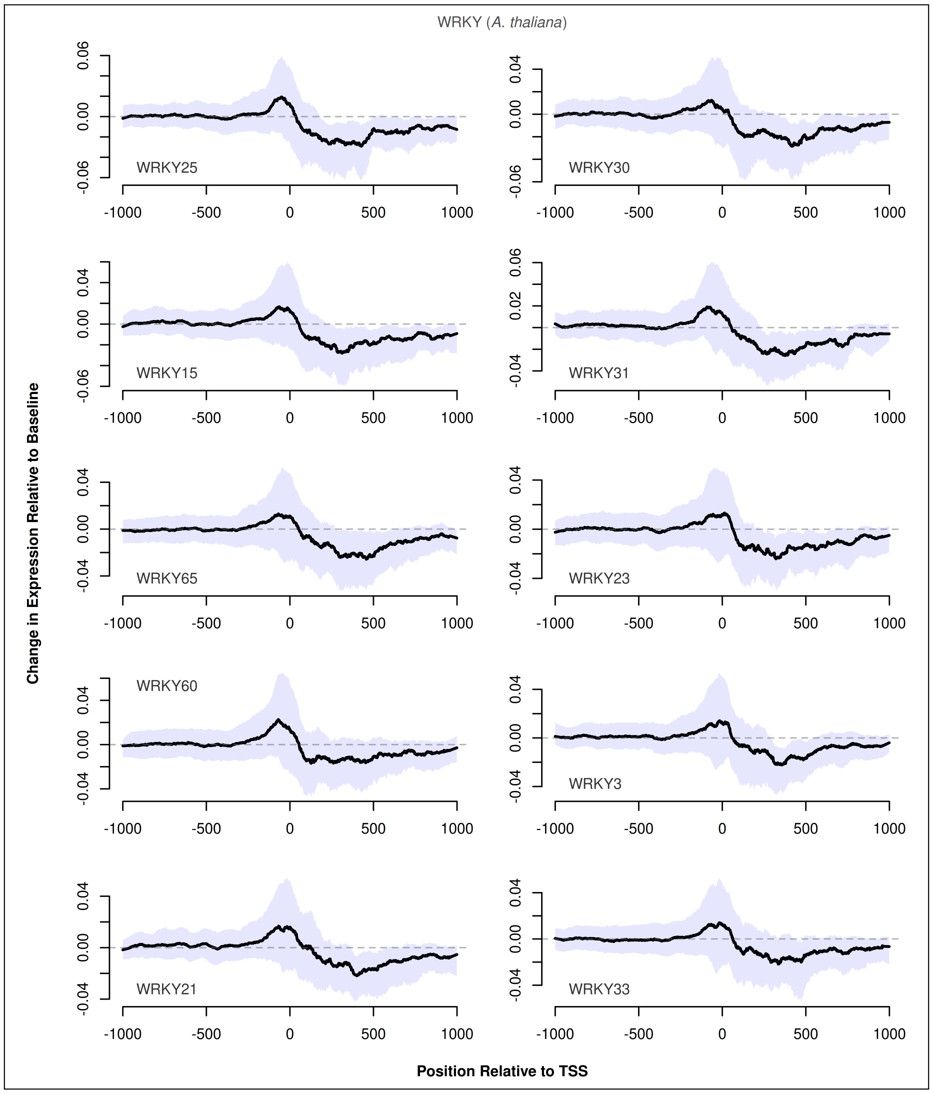


**Fig. S40.** Predicted changes in expression caused by insertion of WRKY TFs with respect to the insertion position in *A. thaliana*. The ten TFs for which the maximum absolute change in expression was predicted are shown, sorted by the maximum absolute change in expression from top-left to bottom-right. Black lines show the median change in expression at each position, while the shaded area shows the upper and lower quartiles. Dashed lines represent baseline expression.
