## Supplementary material for "A deep learning model captures position-specific effects of plant regulatory sequences and suggests genes under complex regulation": Table S

**Table S1.** Express617 RNA-seq sample information for all tissues, stages, and studies

| Tissue | Sample Name | Biological Replicate | Timepoint | Study | SRA |
|---|---|---|---|---|---|
| Leaf | ExL7 | 1 | 7 | This study | |
| Apex | ExA21a | 1 | 21 | This study | |
| Apex | ExA21b | 2 | 21 | This study | |
| Apex | ExA21c | 3 | 21 | This study | |
| Apex | ExA42a | 1 | 42 | This study | |
| Apex | ExA42b | 2 | 42 | This study | |
| Apex | ExA42c | 3 | 42 | This study | |
| Apex | ExA63a | 1 | 63 | This study | |
| Apex | ExA63b | 2 | 63 | This study | |
| Apex | ExA63c | 3 | 63 | This study | |
| Apex | ExA64a | 1 | 64 | This study | |
| Apex | ExA64b | 2 | 64 | This study | |
| Apex | ExA64c | 3 | 64 | This study | |
| Apex | ExA71a | 1 | 71 | This study | |
| Apex | ExA71b | 2 | 71 | This study | |
| Apex | ExA71c | 3 | 71 | This study | |
| Apex | ExA98a | 1 | 98 | This study | |
| Apex | ExA98b | 2 | 98 | This study | |
| Apex | ExA98c | 3 | 98 | This study | |
| Leaf | ExL21 | 1 | 21 | Calderwood et al. (2020) | SRS7070880 |
| Leaf | ExL42 | 1 | 42 | Calderwood et al. (2020) | SRS7070881 |
| Leaf | ExL63 | 1 | 63 | Calderwood et al. (2020) | SRS7070845 |
| Leaf | ExL64 | 1 | 64 | This study | |
| Leaf | ExL71 | 1 | 71 | This study | |
| Leaf | ExL98 | 1 | 98 | This study | |
| Anther | ExAnt1_1 | 1 | Sanders stage 12-13 | This study | |
| Anther | ExAnt1_2 | 2 | Sanders stage 12-13 | This study | |
| Anther | ExAnt1_3 | 3 | Sanders stage 12-13 | This study | |
| Bud | ExBud1_1 | 1 | Sanders stage 6-7 | This study | |
| Bud | ExBud1_2 | 2 | Sanders stage 6-7 | This study | |
| Bud | ExBud1_3 | 3 | Sanders stage 6-7 | This study | |
| Bud | ExBud2_1 | 1 | Sanders stage 10-11 | This study | |
| Bud | ExBud2_2 | 2 | Sanders stage 10-11 | This study | |
| Bud | ExBud2_3 | 3 | Sanders stage 10-11 | This study | |
| Gynoecia | ExGyn1_1 | 1 | Sanders stage 12-13 | This study | |
| Gynoecia | ExGyn1_2 | 2 | Sanders stage 12-13 | This study | |
| Gynoecia | ExGyn1_3 | 3 | Sanders stage 12-13 | This study | |
| Embryo | ExpG3_1 | 1 | Green stage | Woolfenden et al. (2025) | ERS23779109 |
| Embryo | ExpG3_2 | 2 | Green stage | Woolfenden et al. (2025) | ERS23779110 |
| Embryo | ExpG3_3 | 3 | Green stage | Woolfenden et al. (2025) | ERS23779111 |
| Endosperm | ExpE3_1 | 1 | Green stage | Woolfenden et al. (2025) | ERS23779112 |

| | | | | | |
|---|---|---|---|---|---|
| Endosperm | Exp_E3_2 | 2 | Green stage | Woolfenden et al. (2025) | ERS23779113 |
| Endosperm | Exp_E3_3 | 3 | Green stage | Woolfenden et al. (2025) | ERS23779114 |
| Seed coat | ExpSC3_1 | 1 | Green stage | Woolfenden et al. (2025) | ERS23779115 |
| Seed coat | Exp_SC3_2 | 2 | Green stage | Woolfenden et al. (2025) | ERS23779116 |
| Seed coat | Exp_SC3_3 | 3 | Green stage | Woolfenden et al. (2025) | ERS23779117 |
| Silique wall | ExpSW3_1 | 1 | Green stage | Woolfenden et al. (2025) | ERS23779118 |
| Silique wall | ExpSW3_2 | 2 | Green stage | Woolfenden et al. (2025) | ERS23779119 |
| Silique wall | ExpSW3_3 | 3 | Green stage | Woolfenden et al. (2025) | ERS23779120 |
| Gynoecia wall | ExpGW_1 | 1 | Pre-fertilisation (24h before anthesis) | Woolfenden et al. (2025) | ERS23779136 |
| Gynoecia wall | ExpGW_2 | 2 | Pre-fertilisation (24h before anthesis) | Woolfenden et al. (2025) | ERS23779137 |
| Gynoecia wall | ExpGW_3 | 3 | Pre-fertilisation (24h before anthesis) | Woolfenden et al. (2025) | ERS23779138 |
| Embryo | ExpH1_1 | 1 | Heart stage | Woolfenden et al. (2025) | ERS23779121 |
| Embryo | Exp_H1_2 | 2 | Heart stage | Woolfenden et al. (2025) | ERS23779122 |
| Embryo | Exp_H1_4 | 3 | Heart stage | Woolfenden et al. (2025) | ERS23779123 |
| Endosperm | ExpE1_1 | 1 | Heart stage | Woolfenden et al. (2025) | ERS23779124 |
| Endosperm | ExpE1_2 | 2 | Heart stage | Woolfenden et al. (2025) | ERS23779125 |
| Endosperm | Exp_E1_4 | 3 | Heart stage | Woolfenden et al. (2025) | ERS23779126 |
| Seed coat | ExpSC1_1 | 1 | Heart stage | Woolfenden et al. (2025) | ERS23779127 |
| Seed coat | Exp_SC1_2 | 2 | Heart stage | Woolfenden et al. (2025) | ERS23779128 |
| Seed coat | Exp_SC1_4 | 3 | Heart stage | Woolfenden et al. (2025) | ERS23779129 |
| Embryo | ExpM4_1 | 1 | Mature (not fully dry) | Woolfenden et al. (2025) | ERS23779130 |
| Embryo | ExpM4_2 | 2 | Mature (not fully dry) | Woolfenden et al. (2025) | ERS23779131 |
| Embryo | ExpM4_3 | 3 | Mature (not fully dry) | Woolfenden et al. (2025) | ERS23779132 |
| Seed coat | Exp_SCM_1 | 1 | Mature (not fully dry) | Woolfenden et al. (2025) | ERS23779133 |
| Seed coat | Exp_SCM_2 | 2 | Mature (not fully dry) | Woolfenden et al. (2025) | ERS23779134 |
| Seed coat | Exp_SCM_3 | 3 | Mature (not fully dry) | Woolfenden et al. (2025) | ERS23779135 |
| Ovules | ExpOV_1 | 1 | Pre-fertilisation (24h before anthesis) | Woolfenden et al. (2025) | ERS23779139 |
| Ovules | ExpOV_2 | 2 | Pre-fertilisation (24h before anthesis) | Woolfenden et al. (2025) | ERS23779140 |
| Ovules | ExpOV_3 | 3 | Pre-fertilisation (24h before anthesis) | Woolfenden et al. (2025) | ERS23779141 |
| Embryo | ExpT2_1 | 1 | Torpedo stage | Woolfenden et al. (2025) | ERS23779142 |
| Embryo | Exp_T2_2 | 2 | Torpedo stage | Woolfenden et al. (2025) | ERS23779143 |
| Embryo | Exp_T2_4 | 3 | Torpedo stage | Woolfenden et al. (2025) | ERS23779144 |
| Endosperm | ExpE2_1 | 1 | Torpedo stage | Woolfenden et al. (2025) | ERS23779147 |
| Endosperm | ExpE2_2 | 2 | Torpedo stage | Woolfenden et al. (2025) | ERS23779146 |
| Endosperm | Exp_E2_4 | 3 | Torpedo stage | Woolfenden et al. (2025) | ERS23779145 |
| Seed coat | ExpSC2_1 | 1 | Torpedo stage | Woolfenden et al. (2025) | ERS23779148 |
| Seed coat | Exp_SC2_2 | 2 | Torpedo stage | Woolfenden et al. (2025) | ERS23779149 |
| Seed coat | Exp_SC2_4 | 3 | Torpedo stage | Woolfenden et al. (2025) | ERS23779150 |
| Root | RNA_Rt1 | 1 | 9 | This study | |
| Root | RNA_Rt2 | 2 | 9 | This study | |
| Seedling | RNA_Sd1 | 1 | 4 | This study | |
| Seedling | RNA_Sd2 | 2 | 4 | This study | |
| Leaf | RNA_Lf1 | 1 | 28 | This study | |
| Leaf | RNA_Lf2 | 2 | 28 | This study | |
| Silique | RNA_Sq1 | 1 | size | This study | |
| Silique | RNA_Sq2 | 2 | size | This study | |
| Flower | RNA_Fl1 | 1 | size | This study | |
| Flower | RNA_Fl2 | 2 | size | This study | |

The hyperparameters were tuned in six stages, namely, optimizing the architecture, validating the architecture, fine-tuning the architecture, optimizing the training strategy, fine-tuning the training strategy and some final toches. Search spaces are given as a list of items "(item1, item2)" or a numeric range "[from, to]". An asterisk (*) symbol after the closing bracket signifies that the values were sampled from a logarithmic distribution. Hyperparameters that apply for multiple layers were individually optimized for each layer. The best values found are then given as a descending list from shallow layers to deeper layers. P and T signify values that apply to the promoter and terminator branches, respectively. In the first three stages, the hyperparameters were tuned to minimize the validation loss. For all stages the same 80% of data were used as a training set, 10% were used for validation and 10% were held out. In the first stage (Table S2), the Optuna TPE sampler was used to sample hyperparameters hierarchically, meaning that the higher order hyperparameters depend on one or more lower order hyperparameters. The respective conditional relationships are given in gray italics. The conditions correspond to a Boolean hyperparameter above, unless stated otherwise. In the second stage (Table S3) the Optuna grid sampler was used to test the optimized model architecture against simplified versions of the model with shared hyperparameters between layers and/or branches. In the third stage (Table S4) the architecture was fine-tuned using the TPE sampler with a narrow search space around the best values found in the first two stages and small step sizes. Before the fourth stage, a learning rate range test was performed to find the highest possible learning rate, using a modified version of a publicly available range finder (https://github.com/surmenok/keras_lr_finder). This was done to implement a one-cycle learning rate policy. For the learning rate range test and the subsequent hyperparameter optimizations, the batch size was fixed at a value of 130, which is the highest batch size found that did not lead to crashes, due to exceeding the memory capacity. In stages four five and six, hyperparameters relating to the training itself were optimized, including the loss function, therefore, instead of minimizing validation loss, hyperparameters were tuned to maximize the coefficient of determination on the validation set. While in stage four, the respective hyperparameters were tuned across a broad search space, they were tuned across a narrow space around the previously best found vales in stage five, using small step sizes, in order to arrive at the final model. In stage six, some final touches were made to three hyperparameters that potentially influence training efficiency.

**Table S2. Hyperparameter search space for optimizing the neural network architecture**
Optuna TPE sampler used, validation MSE minimized, 100 warm-up trials
Total number of trials: 1122
Number of pruned trials: 892
Number of complete trials: 209
Number of failed trials: 20

| $1^{st}$ order | $2^{nd}$ order | $3^{rd}$ order | $4^{th}$ order | search space | step | best value |
|---|---|---|---|---|---|---|
| outside sequence interval(1) | - | - | - | $[1000, 15000]$ | 1000 | 5000 |
| inside sequence interval(2) | - | - | - | $[100, 1500]$ | 100 | 1200 |
| batch size | - | - | - | $2^{[2,7]}$ | 1 | $2^5$ |
| number of convolutional layers | - | - | - | $[1, 6]$ | 1 | P:6; T:2 |
| number of LSTM layers | - | - | - | $[0, 2]$ | 1 | P:0; T:2 |
| use exponential activation(3) | - | - | - | $(True, False)$ | boolean | *False* |
| *if True* | first_layer_kernel_size | - | - | $2^{[4,5]}$ | 1 | - |
| convolutional kernel size | - | - | - | $2^{[2,5]}$ | 1 | P:$2^{(2,2,4,2,5,5)}$; T:$2^{(2,3)}$ |
| number of convolutional filters | - | - | - | $2^{[5,9]}$ | 1 | P:$2^{(9,8,7,7,9,6)}$; T:$2^{(5,5)}$ |
| pool size | - | - | - | $2^{[0,6]}$ | 1 | P:$2^{(2,2,4,2,5,5)}$; T:$2^{(2,3)}$ |
| pool type(4) | - | - | - | (max, avrg) | categorical | P:(m,a,a,m,a,m); T:(a,m) |
| fraction of pool size used as pool stride(5) | - | - | - | [0.5, 1.0] | 0.1 | P:(0.7,0.5,1.0,0.9,0.5,0.7); T:(0.7,1.0) |
| fraction of kernel size used as conv. stride(5) | - | - | - | [0.0, 0.5] | 0.1 | P:(0.0,0.2,0.2,0.5,0.5,0.3); T:(0.5,0.0) |
| activation function | - | - | - | (ReLU, LReLU, PReLU, SELU, GELU, ELU, Swish, SoftPlus, Mish) | categorical | GELU |
| *if LReLU or PReLU* | alpha parameter of ReLU function(6) | - | - | $[5 \cdot 10^{-6}, 5 \cdot 10^{-1}]$* | continuous | - |
| *if ELU* | alpha parameter of ELU function(7) | - | - | [0.2, 2]* | continuous | - |
| *if not (ReLU or LReLU or PReLU or SELU)* | initialization(8) | - | - | (He_normal, Glorot_normal) | categorical | Glorot_normal |
| 1 - μ (Batchnorm momentum)(9) | | | | $[1 \cdot 10^{-4}, 4 \cdot 10^{-1}]$* | continuous | 0.0823 |
| use dropout(10) | - | - | - | $(True, False)$ | boolean | *True* |
| *if True* | use dropout only after last layer | - | - | $(True, False)$ | boolean | *True* |
| - | *if True* | last layer dropout rate | - | $[10^{-5}, 1]$* | continuous | 0.0677 |
| - | *if False* | dense layer dropout rate | - | $[10^{-5}, 1]$* | continuous | - |
| - | *if False* | use dropout after conv. layers | - | $(True, False)$ | boolean | - |
| - | - | *if True* | conv. layer dropout rate | $[10^{-5}, 1]$* | continuous | - |
| - | *if False & nlstm > 0* | use dropout after LSTM layer | - | $[True, False]$ | boolean | - |
| - | - | *if True* | LSTM layer dropout | $[10^{-5}, 1]$* | continuous | - |
| *if nlstm > 0* | type of skip connection | - | - | (none, add, mult, concat)(11) | categorical | none |
| *if number of LSTM layers > 0* | LSTM units(12) | - | - | $2^{[5,9]}$ | 1 | T: $2^{(5,5)}$ |
| number of dense layers | - | - | - | $[1, 3]$ | 1 | 1 |
| number of dense layer neurons | - | - | - | $2^{[1, 10]}$ | 1 | $2^3$ |

(1) upstream of TSS and downstream of TTS; (2) downstream of TSS and upstream of TTS ;(3) after the first convolutional layer; (4) "max"/"m" = max-pooling, "avrg"/"a" = average pooling; (5) $stride = max(1, \lceil size \cdot fraction \rceil)$; (6) Slope for LReLU and initialization of the slope for PReLU; (7) smoothness of ELU function; (8) For ReLU-like activations, initialization was set to He_normal, for SELU it was set to LeCun_normal; (9) μ = Batchnormalization momenturm; (10) In case of SELU activation, alpha-dropout was used instead of regular dropout; (11) Implementation of skip-connection across LSTM layers. "none" = no skip connection; "add" = element-wise addition; "mult" = element-wise multiplication; "concat" = concatenation along feature dimension; (12) In case of an additive or multiplicative skip connection, the LSTM units of the last LSTM layer were set to the same dimension as the feature axis of the skip-connection, regardless of the hyperparameter value.

**Table S3.** **Hyperparameter search space for validating the architecture**
Optuna grid sampler used, validation MSE minimized
Total number of trials: 49
Number of pruned trials: 0
Number of complete trials: 47
Number of failed trials: 2

| hyperparameter | search space | best value |
|---|---|---|
| promoter branch(1) | [promoter hyperparameters, terminator hyperparameters] | promoter hyperparameters |
| terminator branch(1) | [promoter hyperparameters, terminator hyperparameters] | promoter hyperparameters |
| use default conv. stride(2) | $[True, False]$ | $True$ |
| use default pool stride(3) | $[True, False]$ | $False$ |
| (global) pool type(4) | [max, average, optimized] | average |

(1) Optimal hyperparameters for the promoter and terminator branch were validated by making them interchangeable and testing all possible combinations; (2) the default convolutional stride is 1; (3) the default stride of pooling layers is equal to the respective pooling window size; (4) global pool types (same type used for all pooling layers) were tested against the optimized set of pooling types that were optimized for each individual pooling layer.

**Table S4.** **Hyperparameter search space for fine-tuning the neural network architecture**
Optuna TPE sampler used, validation MSE minimized, 100 warm-up trials
Total number of trials: 1110
Number of pruned trials: 966
Number of complete trials: 135
Number of failed trials: 8

| $1^{st}$ order | $2^{nd}$ order | search space | step | best value |
|---|---|---|---|---|
| batch size | - | $[16, 32]$* | 1 | 18 |
| kernel size (conv. layers 1 and 2) | - | $[2, 8]$* | 1 | P:$(5, 8)$; T:$(4, 4)$ |
| kernel size (conv. layers 3 and 4) | - | $[4, 16]$* | 1 | P:$(9, 10)$; T:$(16, 15)$ |
| kernel size (conv. layers 5 and 6) | - | $[16, 64]$* | 1 | P:$(46, 62)$; T:$(20, 22)$ |
| number of conv. filters(1) | - | $[64, 512]$* | 1 | P:$(288, 106, 227, 187, 64, 152)$; |
| | | | | T:$(219, 449, 259, 154, 105, 152)$ |
| pool size(2) | - | $[1, 64]$* | 1 | $(1, 21, 1, 1, 18, 8)$ |
| fraction of pool size used as stride(3) | - | [0.5, 1.0] | 0.1 | (0.6, 0.7, 0.8, 0.6, 0.7, 0.7) |
| use cross-branch LSTM layer(4) | - | [True, False] | boolean | False |
| *if True* | number of LSTM units | [16, 128]* | 1 | - |
| number of dense layers | - | $[1, 2]$ | 1 | 2 |
| number of neurons (dense layer 1) | - | $[8, 1024]$* | 1 | 750 |
| *# dense layers > 1* | number of neurons (dense layer 2) | $[2, 8]$* | 1 | 3 |
| dropout rate after last layer | - | $[0.005, 0.5]$* | continuous | 0.01063 |
| $1 - \mu^{(5)}$ | - | $[0.01, 0.2]$* | continuous | 0.15784 |

(1) The number of filters for the last convolutional layer was only sampled once and used for both branches, in order to ensure that the output dimensions are identical between the two branches; (2) pool sizes were only sampled once and used for both branches, in order to ensure that the output dimensions are identical between the two branches; (3) $pool\_stride = max(1, \lceil size \cdot fraction \rceil)$; (4) LSTM layer applied to the concatenated outputs of both convolutional branches; (5) µ = Batchnormalization momenturm.

**Table S5.** **Hyperparameter search space for optimizing model training**
Optuna TPE sampler used, validation $r^2$ maximized, 100 warm-up trials
Total number of trials: 623
Number of pruned trials: 0
Number of complete trials: 608
Number of failed trials: 13

| hyperparameter | search space | step | best value |
|---|---|---|---|
| number of training epochs | [3, 30]* | 1 | 6 |
| $1-\beta_2{}^{(1)}$ | [$10^{-5}$,$10^{-1}$]* | continuous | 0.001793 |
| $\varepsilon^{(2)}$ | [$10^{-8}$,$10^{-6}$]* | continuous | $8.327 \cdot 10^{-7}$ |
| weight decay | [$10^{-6}$, $10^{-4}$]* | continuous | $3.209 \cdot 10^{-5}$ |
| learning rate peak shift$^{(3)}$ | [0.2, 0.8] | continuous | 0.7954 |
| final learning rate scale$^{(4)}$ | [0, 1] | continuous | 0.8513 |
| maximum learning rate$^{(5)}$ | [0.01584893, 0.37850893]* | continuous | $2.965 \cdot 10^{-2}$ |
| maximum $\beta_1{}^{(6)}$ | [0.9, 0.99] | continuous | 0.9288 |
| minimum $\beta_1{}^{(6)}$ | [0.8, 0.9] | continuous | 0.8630 |
| huber loss $\delta^{(7)}$ | [0.135, 13.5] | continuous | 11.267 |
| dropout rate after last layer$^{(8)}$ | [0.0, 0.01063] | continuous | $1.243 \cdot 10^{-3}$ |

(1) $\beta_2$ corresponds to the "beta_2" parameter of the Tensorflow implementation of the Nadam algorithm; (2) ε corresponds to the "epsilon" parameter of the Tensorflow implementation of the Nadam algorithm; (3) the relative position, at which the learning rate reaches its peak with respect to the complete cycle length; (4) relative scale of the final learning rate at the end of the cycle, with respect to initial learning rate, see 5; (5) the lower bound of the maximum learning rate is the learning rate at which, during the learning rate range test, the lowest training loss overall was reached, while the upper bound is the highest learning rate that was reached before the training loss began rapidly diverging; $initial\ LR = \frac{max(LR)}{25}$; (6) $\beta_1$ corresponds to the "beta_1" parameter of the Tensorflow implementation of the Nadam algorithm, which corresponds to momentum and is cycled in the opposite direction of the learning rate; (7) corresponds to the "delta" parameter of the Tensorflow implementation of Huber loss; (8) since high learning rates have a regularizing effect, dropout regularization is adjusted.

**Table S6.** **Hyperparameter search space for fine-tuning model training**
Optuna TPE sampler used, validation $r^2$ maximized, 10 warm-up trials
Total number of trials: 533
Number of pruned trials: 0
Number of complete trials: 511
Number of failed trials: 21

| hyperparameter | search space | step | best value |
|---|---|---|---|
| number of training epochs$^{(1)}$ | [6, 10] | 1 | 6 |
| final learning rate scale | [0, 0.9] | continuous | 0.8758 |
| maximum learning rate | [$2.965 \cdot 10^{-2}$, $5 \cdot 10^{-2}$] | continuous | $4.873 \cdot 10^{-2}$ |
| initial learning rate | [$\frac{max.\ learning\ rate}{25}$, $\frac{max.\ learning\ rate}{20}$] | continuous | $1.299 \cdot 10^{-3}$ |
| maximum $\beta_1$ | [0.9, 0.95] | continuous | 0.9274 |
| minimum $\beta_1$ | [0.83, 0.89] | continuous | 0.8899 |
| huber loss $\delta^{(2)}$ | [0.135, 13.5]* | continuous | 2.296 |

(1) The learning rate peak is shifted according to the number of epochs, to maintain a constant relative location with respect to the whole cycle length; (2) sampled from logarithmic distribution in this stage.

**Table S7. Hyperparameter search space for final touches to model training**
Optuna TPE sampler used, validation $r^2$ maximized, 10 warm-up trials
Total number of trials: 322
Number of pruned trials: 0
Number of complete trials: 311
Number of failed trials: 9

| hyperparameter | search space | step | best value |
|---|---|---|---|
| maximum learning rate | [0.01, 0.1] | continuous | $6.630 \cdot 10^{-2}$ |
| Batchnormalization Momentum ($\mu$)$^{(1)}$ | [0.7, 0.9] | continuous | 0.8167 |
| huber loss $\delta$ | [1.0, 3.0] | continuous | 2.9466 |

(1) The optimal value of $\mu$ depends on the batch size. Since it had previously been optimized at lower batch sizes, it needed to be adjusted to the high batch size used for the one-cycle learning rate policy.

**Table S8.** Final set of best hyperparameters

| hyperparameter | best value |
|---|---|
| batch size | 130 |
| kernel initialization | Glorot normal |
| activation function | GELU |
| length of outside sequence interval | 5000 |
| length of inside sequence interval | 1200 |
| number of convolutional layers | 6 |
| number of conv. filters | P:$(288, 106, 227, 187, 64, 152)$; T:$(219, 449, 259, 154, 105, 152)$ |
| size of conv. kernels | P:$(5, 8, 9, 10, 46, 62)$; T:$(4, 4, 16, 15, 20, 22)$ |
| pooling type | average pooling |
| size of pooling windows | P&T: (1, 21, 1, 1, 18, 8) |
| pooling strides | P&T: (1, 15, 1, 1, 13, 6) |
| number of dense layers | 2 |
| number of dense layer neurons | $(750, 3)$ |
| dropout rate after last layer | $1.243 \cdot 10^{-3}$ |
| number of epochs | 6 |
| learning rate peak shift | 0.7954 |
| initial learning rate | $1.299 \cdot 10^{-3}$ |
| maximum learning rate | $6.630 \cdot 10^{-2}$ |
| final learning rate scale | 0.8758 |
| initial Nadam $\beta_1$ | 0.9274 |
| minimum Nadam $\beta_1$ | 0.8899 |
| Nadam $\beta_2$ | 0.9982 |
| Nadam $\varepsilon$ | $1.133 \cdot 10^{-6}$ |
| Nadam weight decay | $3.209 \cdot 10^{-5}$ |
| Batchnormalization momentum ($\mu$) | 0.8167 |
| huber loss $\delta$ | 2.9466 |
